# Harnessing Landrace Diversity Empowers Wheat Breeding for Climate Resilience

**DOI:** 10.1101/2023.10.04.560903

**Authors:** Shifeng Cheng, Cong Feng, Luzie U. Wingen, Hong Cheng, Andrew B. Riche, Mei Jiang, Michelle Leverington-Waite, Zejian Huang, Sarah Collier, Simon Orford, Xiaoming Wang, Rajani Awal, Gary Barker, Tom O’Hara, Clare Lister, Ajay Siluveru, Jesús Quiroz-Chávez, Ricardo H. Ramírez-González, Ruth Bryant, Simon Berry, Urmil Bansal, Harbans S. Bariana, Malcolm J. Bennett, Breno Bicego, Lorelei Bilham, James K.M. Brown, Amanda Burridge, Chris Burt, Milika Buurman, March Castle, Laetitia Chartrain, Baizhi Chen, Worku Denbel, Ahmed F. Elkot, Paul Fenwick, David Feuerhelm, John Foulkes, Oorbessy Gaju, Adam Gauley, Kumar Gaurav, Amber N. Hafeez, Ruirui Han, Richard Horler, Junliang Hou, Muhammad S. Iqbal, Matthew Kerton, Ankica Kondic-Spica, Ania Kowalski, Jacob Lage, Xiaolong Li, Hongbing Liu, Shiyan Liu, Alison Lovegrove, Lingling Ma, Cathy Mumford, Saroj Parmar, Charlie Philp, Darryl Playford, Alexandra M. Przewieslik-Allen, Zareen Sarfraz, David Schafer, Peter R. Shewry, Yan Shi, Gustavo Slafer, Baoxing Song, Bo Song, David Steele, Burkhard Steuernagel, Phillip Tailby, Simon Tyrrell, Abdul Waheed, Mercy N. Wamalwa, Xingwei Wang, Yanping Wei, Mark Winfield, Shishi Wu, Yubing Wu, Brande B.H. Wulff, Wenfei Xian, Yawen Xu, Yunfeng Xu, Quan Yuan, Xin Zhang, Keith J. Edwards, Laura Dixon, Paul Nicholson, Noam Chayut, Malcolm J. Hawkesford, Cristobal Uauy, Dale Sanders, Sanwen Huang, Simon Griffiths

## Abstract

Breeding crops resilient to climate change is urgently needed to help ensure food security. A key challenge is to harness genetic diversity to optimise adaptation, yield, stress resilience and nutrition. We examined the genetic and phenotypic diversity of the A.E. Watkins landrace collection of bread wheat (*Triticum aestivum*), a major global cereal, through whole-genome re-sequencing (827 Watkins landraces and 208 modern cultivars) and in-depth field evaluation spanning a decade. We discovered that modern cultivars are derived from just two of the seven ancestral groups of wheat, leaving five groups as previously untapped sources for breeding. This provides access to landrace-specific functional variations using structured germplasm, genotyping and informatics resources. Employing complementary genetic populations and approaches, we identified thousands of high-resolution quantitative trait loci (QTL) and significant marker–trait associations for major traits, revealing many Watkins-unique loci that can confer superior traits in modern wheat. Furthermore, we identified and functionally verified causative genes for climate-change adaptation, nutritional enhancement and resistance to wheat blast. Finally, we assessed the phenotypic effects of 44,338 Watkins-unique haplotypes, introgressed from 143 prioritised QTL in the context of modern cultivars, bridging the gap between landrace diversity and current breeding. This study establishes a framework for systematically utilising genetic diversity in crop improvement to achieve sustainable food security.

## Main text

The world population is projected to increase by two billion people over the next 30 years, placing greater demands on wheat (currently 20% of global diets) as a vital source of calories, protein, minerals, and fibre^1^. Our ability to meet this growing demand is threatened by climate change and geopolitical instability, which have a multiplying effect when they disrupt the narrow global wheat export base^2^. Just five countries (Russia, USA, Canada, France and Ukraine) dominate wheat exports, while major population centres such as China require adequate imports to satisfy internal demands for wheat^3^. To further compound these challenges, yield gains in wheat and other crops have slowed, in part due to the narrowing genetic diversity of modern cultivars.

Historically, farmers relied on locally adapted domesticated crop cultivars known as landraces, which could withstand numerous severe environmental hardships, but compared to modern cultivars, were relatively low yielding^4^. Compared to modern cultivars, landraces have been less exposed to historical and geographical founder effects, making them a rich, albeit underutilised, source of genetic diversity. Despite the potential of landraces for developing more resilient and nutritious crops, the full extent of their value is not yet understood, and how this value could be utilised in breeding programmes remains unclear.

Integrating new beneficial alleles from landraces into modern cultivars poses multiple scientific, technical and economic obstacles^5^. These include a lack of genetic resources with associated sequence information needed to identify rare useful alleles. Moreover, bread wheat possesses distinctive characteristics, including a large genome (16 Gbp), recent allohexaploidy which favours dominant gain-of-function alleles^6^, a complex genetic architecture of quantitative traits and a convoluted domestication history with frequent introgressions from wild relatives^7^. Consequently, the opportunity to deliver beneficial alleles precisely, particularly rare alleles, is hindered by the limited availability of resources for their discovery, deployment and selection.

In this study, we present the collaborative work of an international consortium for implementing systematic gene discovery for breeding that has overcome these obstacles and harnessed untapped wheat landrace diversity (Extended Data Fig. 1). Our strategy capitalised on the rich genetic, geographic and phenotypic diversity within the A.E. Watkins landrace collection of bread wheat, comprising 827 accessions collected from 32 countries in the 1920s and 1930s^8^. We implemented a pre-breeding strategy^9^ to decode, dissect, discover, design and deliver progress in breeding. To do this, we combined gene discovery populations^10^ developed from the Watkins collection with a genomic variation matrix, haplotype map and in-depth field phenotyping for quantitative traits segregation within Watkins-derived structured populations. This comprehensive approach generated an integrated set of tools that provides the research and breeding communities with access to new, beneficial diversity (Extended Data Fig. 2).

### WHEAT GENOMIC COMPARISONS REVEALS RICH FUNCTIONAL DIVERSITY

To discover novel genetic variations available in the Watkins bread wheat landrace collection (hereafter ‘Watkins’), we conducted 12.73X whole-genome re-sequencing of its 827 accessions (Supplementary Table 1 and Supplementary Fig. 1). We aligned these sequences to the IWGSC RefSeq v1.0 bread wheat reference genome^11^. Based on the identified SNPs (Supplementary Tables 2–4 and Supplementary Fig. 2), Watkins was categorised into seven ancestral groups (AGs, individual groups designated as AG1 to AG7, Fig. 1a and 1b, Extended Data Fig. 3, Supplementary Fig. 3-4 and Supplementary Table 5). The geographical collection points of the landrace accessions for each AG represent all the major wheat-growing areas of the world in the 1920s (Fig. 1a, Supplementary Table 6). To explore the relationships of modern wheat with the AGs, we selected a set of 208 20^th^ century wheat cultivars for whole-genome sequencing, including 15 previously described accessions^12^, which embody the maximal diversity contained in the 1169 cultivars from 25 countries (Extended Data Fig. 4) genotyped using the Axiom 35K Array^13^ (CerealsDB database), hereafter referred to as ‘modern wheat’ (Supplementary Table 1).

**Fig. 1.**
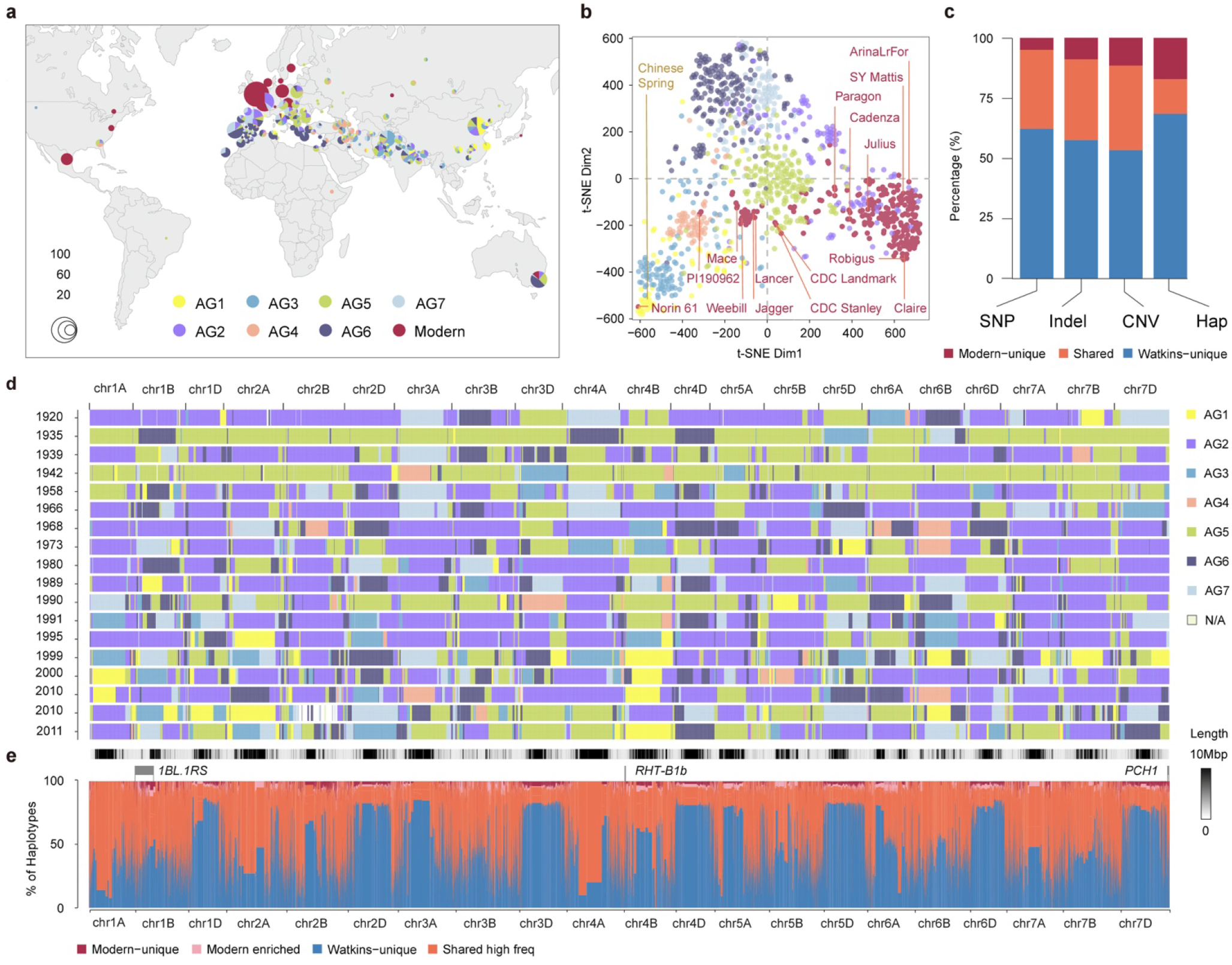
Genomic variants in Watkins landraces compared to modern wheat. **a,** Geographical distribution of all accessions, including the entire Watkins collection (*n* = 827) and modern wheat cultivars (*n* = 224; comprising 208 cultivars sequenced in this study and 16 previously described modern wheat cultivars including Chinese Spring). The seven ancestral groups (AG1–7) derived from Watkins and modern wheat are colour-coded. **b,** t-SNE plot based on the 10M SNPs shared by different ancestral groups. The SNPs were stringently controlled by linkage disequilibrium (LD-based; see Methods), with AG1–7 and modern wheat colour coded as in panel **a**. The distribution of the 10+ Genome lines and Chinese Spring are shown. **c,** Percentage of Watkins-unique (shown in blue), shared (orange) and modern-unique (red) variants for SNPs, short Indels (< 50 bp), gene copy number variations (CNV) and haplotypes (Hap). **d,** *k*-mer based IBSpy long-range haplotype analysis of 18 representative modern wheat cultivars (released from 1920 to 2011, see Source Data). IBS regions between modern wheat and landraces are shown as coloured blocks (top 100 Watkins accessions; see Methods, Supplementary Table 10). **e,** Genomic distribution and comparison of haplotypes between Watkins and modern wheat along the 21 chromosomes, including the proportion of haplotypes that are absent (Watkins-unique; blue), shared with high frequency (orange), modern-enriched (pink) or unique (red) to modern wheat. The haplotypes were identified based on linkage disequilibrium by Plink (Methods), with single-base-resolution based on the IWGSC v1.0 reference genome.

Taking Watkins and modern wheat together, we identified ∼262 million high-quality single nucleotide polymorphisms (SNPs; Supplementary Tables 2–4). The SNP composition of modern wheat largely overlaps with that of AG2 and AG5 (Fig. 1b), which have Western and Central European origins, respectively, suggesting that these AGs supplied the founder lines of modern wheat. The AG2/5 hypothesis for the origins of contemporary wheat cultivars is well supported by independent wheat genomics data sets^14^ (Extended Data Fig. 3c) and other non-PCA-based methods such as ‘identity by state’ (IBS) distance and population differentiation analyses^15,16^ (Supplementary Fig. 5). Watkins contains variants that are absent in modern wheat. Specifically, we identified 162 million SNPs (62%), 9.7 million insertions or deletions (57%) and 57,000 copy number variants (CNVs, 53%) that are unique to Watkins (Fig. 1c, Supplementary Figs. 6–8 and Supplementary Tables 7–9). These variants are predominantly carried by five AGs (1, 3, 4, 6 and 7), showing almost no overlap with modern wheat (Supplementary Figs. 9–10). These data indicate that the five phylogenetically isolated AGs are highly diverse and represent a reservoir of previously untapped genotypic diversity for contemporary wheat breeding.

To further explore the landrace origins of modern wheat, we used long-range haplotypes to visualise the mosaic of IBS regions across their genomes (Fig. 1d, Supplementary Table 10). These IBS segments are signatures of the close relatives of Watkins landrace accessions that were the founder lines of modern wheat cultivars, which have retained high chromosome-level identity with AG2/5 landraces, often across multi-megabase tracts extending across the majority of a chromosome’s length (Supplementary Fig. 11). On average, IBS segments remained intact along a length of 159.78 Mbp in centromeric regions, whereas they were shorter in the more recombinogenic distal regions (using the R1–3 chromosome segment designations^17^: 23.96 and 38.36 Mbp for R1 and R3 regions, respectively; Supplementary Fig. 12). In addition to confirming that AG2 and AG5 were the most significant contributors to modern wheat, the IBS analysis provided insight into the very small effective population size of modern wheat. As few as 26 Watkins accessions could be modelled as virtual donors of IBS segments to reconstitute >50% of the modern wheat genomes included here (Fig. 1e and Supplementary Fig. 13).

To map the genomic locations of variants that are absent from modern wheat, we used linkage disequilibrium (LD)-based haplotype analysis^18^ (Extended Data Fig. 5, Supplementary Fig. 14). This identified 71,282 haploblocks, of which 69.6% (49,626) only contain the Watkins-unique haplotypes (median 5 and mean 11.85 haplotypes/haploblock; Supplementary Figs. 15–16 and Supplementary Tables 11–12). We aligned these haploblocks to the IBS chromosome map of modern wheat, revealing the potential for new arrangements of chromosomal segments to enrich the current IBS structure of elite wheat germplasm with Watkins-unique haplotypes (Fig. 1e). However, in addition to the unique variants identified in Watkins, 2.5% of the unique haplotype variants were found in modern wheat, several of which were associated with introgressions from wheat wild relatives made by breeders (e.g., *1BL/1RS*^19^, *RHT1*^15^, *Pch1*^20^), after the AG2/5 landrace foundation of modern wheat (Fig. 1e).

To assess the potential for Watkins-unique genetic diversity to influence functional traits, we studied variations occurring in or around genes. Among all the Watkins-unique SNPs, 325,915 were predicted to have functional significance (Supplementary Tables 4 and 13–14). Particularly noteworthy among these are the Watkins-unique functional SNPs found near the 13,902 genes that are monomorphic in modern wheat, meaning that there is no standing variation available to improve the associated traits through traditional breeding (Supplementary Fig. 17; Supplementary Tables 15–17). According to ontology term analysis, these genes control diverse biological processes that could affect important agronomic traits such as yield, stress tolerance, nutritional quality and disease resistance (Supplementary Tables 15–17). To leverage these genomics resources (Extended Data Fig. 1, 2, 6) and to search for useful trait variations associated with these variants, we undertook a programme for high-throughput quantitative trait locus (QTL) discovery.

### TARGETING LANDRACE QTL ALLELES FOR BREEDING

Traits required for climate change adaptation and mitigation are often controlled by multiple genetic loci in a quantitative manner^21,22^. Thus, structured populations combined with specialised statistical analyses^23^ are required for their study. We used the 827 Watkins accessions as a genome-wide association panel for phenotypic datasets recorded in the UK and China (Supplementary Fig. 18 and Supplementary Tables 18–19<). We also used 73 Paragon x Watkins recombinant inbred line (RIL) populations^10^ (Fig. 2a), representing the seven AGs (Supplementary Fig. 18 and Supplementary Table 20), resulting in 6,762 RILs for which whole-genome imputation was performed. We recorded phenotypic data for 137 traits (Supplementary Tables 21–22), covering the major categories of grain yield, nutritional quality, adaptation, and abiotic and biotic stress tolerance (Fig. 2b). These field experiments were conducted over ten years in ten environments, resulting in over 2 million observations and data points (Fig. 2c and 2d, Extended Data Fig. 1, 2, 7 and Supplementary Table 23). The structured populations allowed us to capitalise on the complementary strengths of joint linkage and association studies for complex traits by nested association mapping (NAM) studies, as well as classical QTL analysis in biparental populations and gene discovery strategies such as genome-wide association studies (GWAS) (Methods, Extended Data Fig. 8 and Supplementary Fig. 19 and Supplementary Tables 24– 28).

**Fig. 2.**
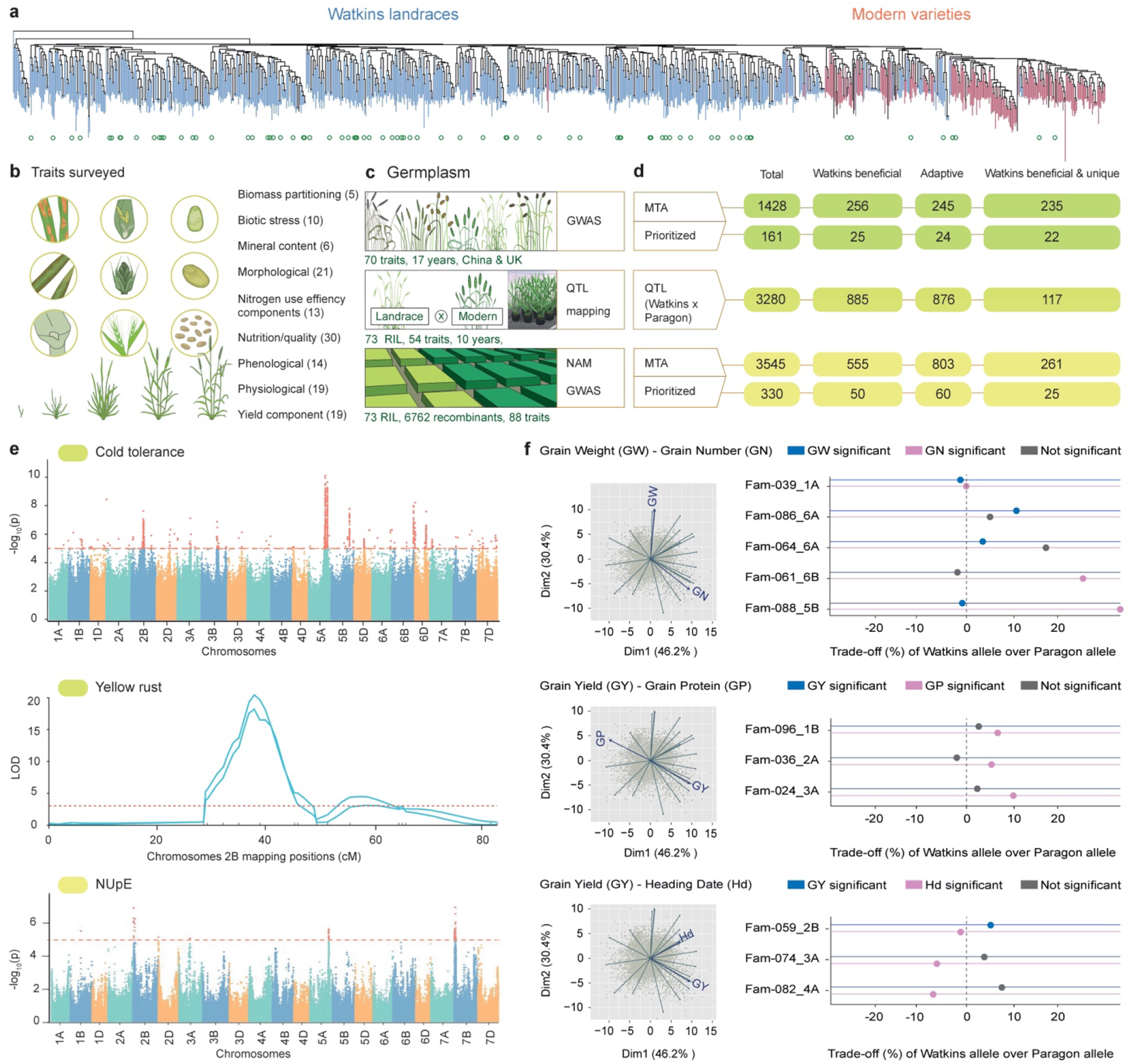
Genetic dissection of new and useful traits from Watkins. **a,** SNP-based phylogeny of Watkins landraces (blue) and modern varieties (pink). The Watkins parents of 73 RIL populations used here are marked with empty green circles. **b,** Overall summary and schematic illustration of the phenotypic trait data collected in this study categorised into nine high-level trait classes. The total number of sub-traits for each category is shown in parenthesis. **c,** Field experiments and trait data collected from the Watkins natural population for GWAS analysis (top panel), QTL analysis from individual segregating mapping populations (RIL; middle panel), and combined analysis of Nested Association Mapping (NAM) GWAS with RILs from the imputation of the NAM populations (bottom panel). **d,** Prioritisation of detected genetic effects. The total number, and the number of prioritised marker–trait associations (MTAs) detected from GWAS (top panel) and from the NAM population (bottom panel). The total number of QTL identified using biparental mapping populations (Paragon x Watkins) is summarised in the middle panel. Watkins beneficial are the number of allelic effects from QTL, GWAS and NAM GWAS in which the Watkins allele exceeded the Paragon allele for traits under directional selection. Adaptive are effects that can be beneficial in either direction depending on the breeding objective. Watkins beneficial/adaptive and unique are alleles for which the haplotype under the peak marker/genomic interval was absent or present at extremely low frequency in modern wheat. **e,** Examples of genetic effect detection using GWAS (cold resistance), biparental QTL analysis (yellow rust resistance) and NAM GWAS for Nitrogen Uptake Efficiency (NUpE) using all data collected from experiments in panel **c**. **f,** Representative principal component analysis (PCA) plots (left) of Watkins NAM data from a single environment. highlighting three general trait trade-off relationships (Grain Weight vs. Grain Number, Grain Yield vs. Grain Protein and Grain Yield vs. Heading Date). Plots on right show the percentage of phenotypic differences for the labelled traits between NILs carrying the Watkins allele over the Paragon allele across a target QTL region. Each QTL region is represented by a NIL pair or family (Fam; Supplementary Table 48). Significant effects (*P* < 0.05) for traits are represented by coloured/grey circles.

Combining the mapping populations with sequence-based haplotypes, NAM-GWAS captures both historical and RIL population recombinations as well as common alleles from the natural population and rare useful alleles with amplified frequency in the segregating populations. Using this approach, we calculated robust QTL effects at haplotype resolution and determined the distribution of useful QTL alleles between Watkins landraces and modern wheat (Fig. 2e and Supplementary Tables 29–32). Useful QTL alleles are those that increase a phenotypic value for traits always selected in the same direction, such as yield and disease resistance, or those that provide new adaptability for selection in either phenotypic direction (Extended Data Fig. 9), such as heading date. Nitrogen Use Efficiency (NUE) is a complex quantitative trait that is dependent on multiple biological processes and agronomic traits whose genetic architecture and regulatory network remain largely unknown. Here, we measured NUE components NUtE (Nitrogen Utilisation Efficiency) and NUpE (Nitrogen Uptake Efficiency) in multiple field trials with high (200 kg N ha^−1^) and low (50 kg N ha^−1^) nitrogen inputs conducted over the past decade and identified 26 significant associated loci across several chromosomes (Extended Data Fig. 10, and Supplementary Table 27). Of particular note, we detected a significant signal with candidate genes underlying the target genomic interval on chromosome 5A, including genes encoding a transcription factor involved in regulating senescence (*NLP9*, *TraesCS5A02G349500*), an NPF nitrate transporter (*NPF2.2*, formerly known as *NRT1.2*, *TraesCS5A02G388000*) and an ammonium transporter (*AMT 2.2*, *TraesCS5A02G388100*). These genes showed novel allelic diversity, with 38 new haplotypes that were only present in Watkins (Supplementary Table 33).

In total, we identified 8,253 genetic effects (3,280 QTL, 1,428 GWAS and 3,545 NAM-GWAS marker–trait associations, Extended Data Fig. 8 and Supplementary Tables 25–27). Based on the direction of the allelic effects, 1,696 have the potential to improve modern cultivars such as Paragon, and 36% (613) of the most significantly associated SNPs are located within haplotypes that are absent in modern wheat (Fig. 2d and Supplementary Table 24). Despite reduced genetic resolution compared to NAM and GWAS, the use of single biparental populations was essential for detecting QTL from very rare Watkins haplotypes. For example, just 33 Watkins accessions exhibit resistance to the ‘Warrior’ race of *Puccinia striiformis* (the causal agent of yellow rust disease; Fig. 2e, Supplementary Table 34), a recently emerged, highly aggressive race with increased pathogenicity at elevated temperatures^24^. Iran is the dominant country of origin for these resistant accessions (14 out of 33). GWAS did not identify significant MTAs for these resistance loci, but biparental QTL mapping identified 15 new loci conferring yellow rust resistance in the UK and Australia (Supplementary Tables 35–36). Twelve of these resistance loci are carried by accessions outside of the modern wheat AG2/5 founder complex, again with the majority from Iran (5).

Complementary to bi-parental QTL mapping, GWAS of Watkins accessions maximised the allelic diversity captured in a single experiment. In ShanDong province (North China Plain), we detected an association of the *CBF* (C-repeat binding factor) gene cluster with tolerance to lethal cold winter temperatures (LD75; Fig. 2e). *CBF* diversity is more pronounced in Watkins accessions than in modern wheat, with 40% novel allelic diversity and 16% AG-specific presence-absence variations or copy number variations identified in these accessions (Supplementary Tables 37–40 and Supplementary Figs. 20–22). The increased depth of functional diversity described for the common large-effect *CBF* gene family was also found in genes controlling many other quantitative traits such as flowering time in Watkins, for which again NAM GWAS demonstrates its power. We identified novel allelic diversity for the central floral regulator *FT1*, including indels in its promoter and changes in CNV important for environmental adaptation. This novel diversity appears to be specific to particular AGs and is often absent in modern wheat (Supplementary Fig. 23). Similarly, for *PPD1*, a major photoperiod gene in wheat, most alleles used in modern breeding are present in the B and D genomes. We conducted photoperiod sensitivity screens of Watkins, followed by QTL analysis of derived RIL populations and NIL confirmation, and identified new A genome alleles that are absent in modern wheat (Supplementary Figs. 24–26). These results highlight the potential of using the large set of genetic effects identified here to help deliver new traits for adaptation to and mitigation against the effects of climate change in wheat.

To elucidate the potential utility of these alleles for practical wheat improvement through breeding, we investigated the relationships between multiple key traits (Fig. 2f). This approach is highly relevant, as QTL for one agronomic trait can often be antagonistically coupled to other breeding targets^25^. Thus, uncoupling these relationships, often thought of as trade-offs, is crucial for accelerating the breeding process. We developed near isogenic lines (NILs) to test the extent to which these trait relationships (e.g., grain weight vs. grain number, grain yield vs. grain protein content) were upheld for individual QTL effects (Fig. 2f and Supplementary Table 41). For each locus-trait combination, we found a range of penetrance for these mainly antagonistic relationships, with several positive-effect QTL for one trait being either neutral or positive for the other trait, reversing the general relationship and providing a route for selection without the trade-offs that the founding breeders of modern wheat could not avoid (Fig. 2f).

This understanding of trade-offs helped us form a strategy for the recovery of beneficial traits that have been lost in modern breeding due to preferential selection pressure for one of the paired traits. For example, breeding for reduced crop height to avoid lodging and increase yield via harvest index^25^ culminated in the development of the semi-dwarf Green Revolution wheat cultivars^26^ (Fig. 3a). Their low stature was likely achieved at the expense of crop biomass, which is also an important physiological component of grain yield^25^. Of the 291 QTL identified for plant height, 187 conferring reduced height were derived from the cultivar Paragon (Fig. 3b and Supplementary Table 25). We focused on a height-increasing Watkins haplotype on chromosome arm 7BL with no effect on harvest index, whereas other height-increasing QTL alleles (e.g., chromosome 6A) had consistent negative effects on harvest index. The local linkage disequilibrium block corresponding to the chromosome arm 7BL target region contains contiguous haplotypes that are either absent or present at low frequency in modern wheat (Fig. 3c-f). Upon testing in NILs, we determined that these haplotypes were associated with a height increase of 9.37 cm (*P* = 0.002) and also a grain yield increase of 0.39 t ha^−1^ (*P* < 0.002) compared to Paragon (Fig.3g, Supplementary Fig. 27). This novel QTL on chromosome arm 7BL is currently being bred into modern commercial cultivars.

**Fig. 3.**
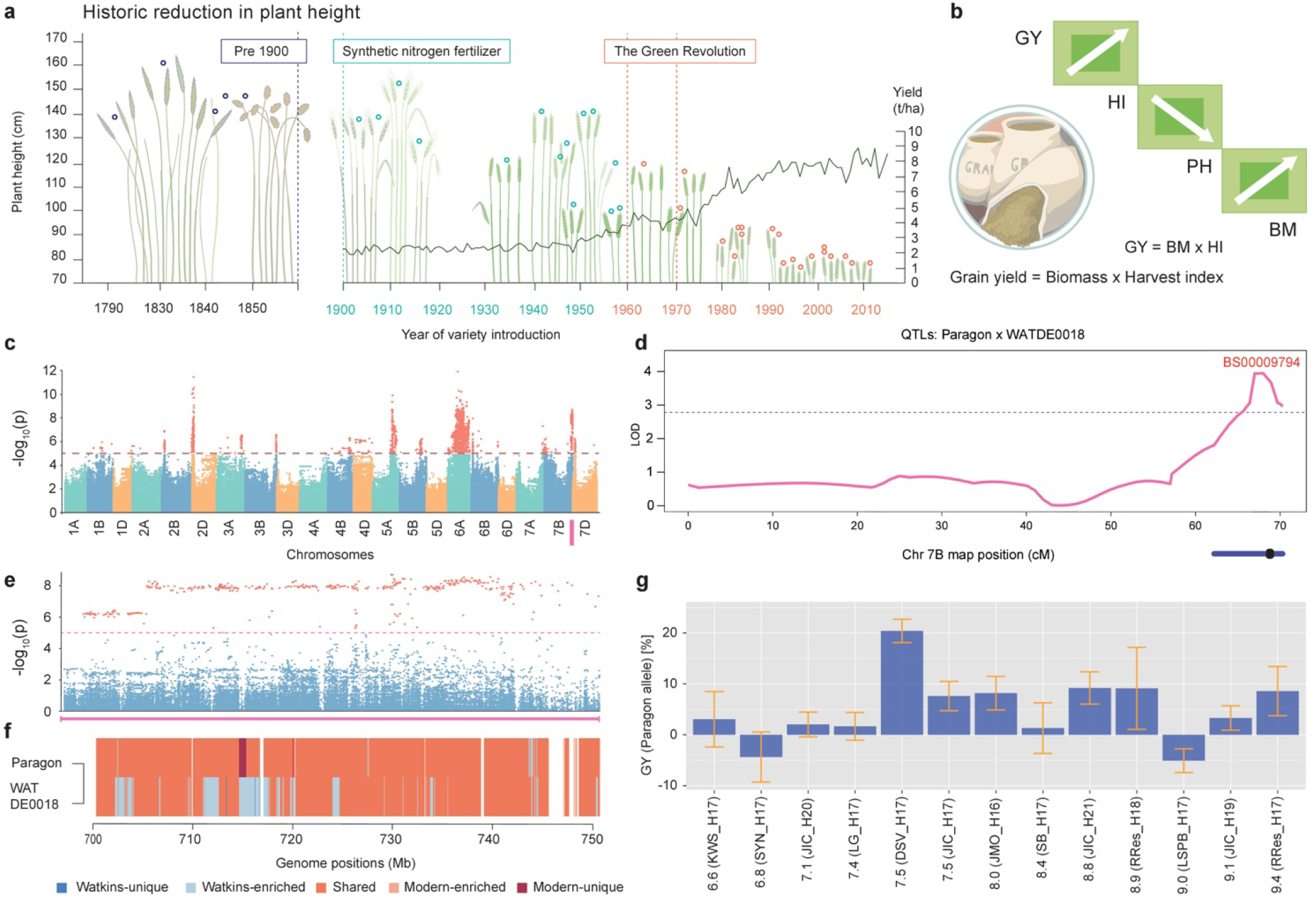
Recovery of useful diversity left behind by the Green Revolution, exemplified by the changes in plant height and grain yield during modern breeding history. **a,** Historical reduction in plant height of wheat cultivars. Empty circles represent height from heritage accessions cultivated and cultivars released from 1790 to 2010^48^. Yields (t ha^−1^) from 1900 to the present are shown. Agricultural milestones are highlighted. **b,** Illustration of trait relationships between Grain Yield (GY), Harvest Index (HI), Plant Height (PH) and Biomass (BM), where GY = HI x BM, and BM is positively correlated with PH, which is negatively correlated with HI. **c,** Dissection of the genetic architecture of plant height performed by Watkins NAM GWAS. **d,** Plant height QTL on chromosome arm 7BL and **e,** local association plot across the chromosome arm 7BL region based on NAM GWAS data shown in **c**. **f,** Frequency distribution of haplotypes carried by Paragon and WATDE0018 (chromosome arm 7BL QTL carrier) in the NAM GWAS target marker–trait association locus (**e**). Comparisons for haplotype frequency were performed between Watkins-unique (dark blue), Watkins enriched (light blue), shared (orange), modern enriched (light pink) and modern-unique (dark red). **g,** Effect of the WATDE0018 segment on Grain Yield (GY) with respect to the Paragon haplotype in NILs. Numbers at the start of each label represent mean GY in t ha^−1^. H: year harvested; the harvest locations include KWS: KWS, SYN: Syngenta, JIC: John Innes Centre-Church Farm, LG: Limagrain, DSV: DSV, JMO: John Innes-Morley, SB: Sutton Bonington, RRes: Rothamsted, and LSPB: LS Plant Breeding. Error bars denote standard errors of the mean.

By exploring the genetic basis of the antagonism between grain yield and component traits, we identified modern applications for allelic effects that were left behind early in the history of wheat breeding, highlighting the potential of this approach for decoupling other key antagonistic trait relationships (Fig. 2f).

### WATKINS GENOMIC RESOURCES ACCELERATE GENE DISCOVERY

In addition to mining novel QTL alleles, targeting individual genes with major effects within the polyploid wheat genome context is useful for exploiting wheat landrace diversity. Thus, we used Watkins genomics and genetics resources to identify causal genes for traits that are increasingly relevant given changes in climate (e.g., heat adaptation, emerging diseases) and dietary patterns (e.g., increased nutritional density).

Many of the world’s major wheat-growing regions are facing increasingly warm, dry summers with negative consequences for wheat production^27^. This is exacerbated by the presence of *RHT1* semi-dwarfing Green Revolution alleles that have negative pleiotropic effects, such as reduced early crop vigour in very dry summers typical of Mediterranean climates^28^. We capitalised on historic adaptation processes of crops in these regions to guide the selection of genes (such as the semi-dwarfing *RHT8* gene) that hold promise for adaptation to future climate scenarios^28^. Using NILs, we confirmed the beneficial effects of *RHT8* in Mediterranean environments (reduced height, increased yield) compared to its negative antagonistic effects (including reduced height and reduced yield) in more temperate UK environments (Fig. 4a and Supplementary Table 42). *RHT8* was introduced into Europe from the Japanese landrace ‘Akakomugi’ ^29^ and was genetically defined using the Akakomugi-derived cultivar ‘Mara’. Although *RHT8* has been mapped at high genetic resolution^30^, it has not yet been cloned. Whereas previous studies have suggested that a closely linked gene (*TraesCSU02G024900, RNHL-D1*) also affects plant height^31^, this gene is located outside of the genetic interval, and the proposed diagnostic/causative SNP is absent in Mara. We fine mapped *RHT8* to a 0.82 Mbp interval containing 23 gene models (Supplementary Tables 43–45). We identified 36 Watkins accessions with the Mara-like haplotype and used markers derived from novel Watkins SNPs, reducing the *RHT8* mapping interval to a physical distance of 6.7 kbp (Fig. 4b). This interval contains two annotated genes in IWGSC RefSeq v1.0, *TraesCS2D02G057800* (unknown function) and *TraesCS2D02G057900* (Photosystem 1 assembly 2), positioned in a head-to-head orientation with only 6 bp between their respective 5’ untranslated regions and 250 bp between their translational start codons. The expression of the two genes is tightly correlated (Fig. 4c): when *TraesCS2D02G057800* expression is high (as it is until mid-stem extension), *T*raesCS2D02G057900 expression is low, and vice versa (Fig. 4c; Pearson correlation r = −0.979; *P* < 0.0001). These findings demonstrate the potential of using sequence variations in landraces to identify candidate genes for key agronomic traits.

**Fig. 4.**
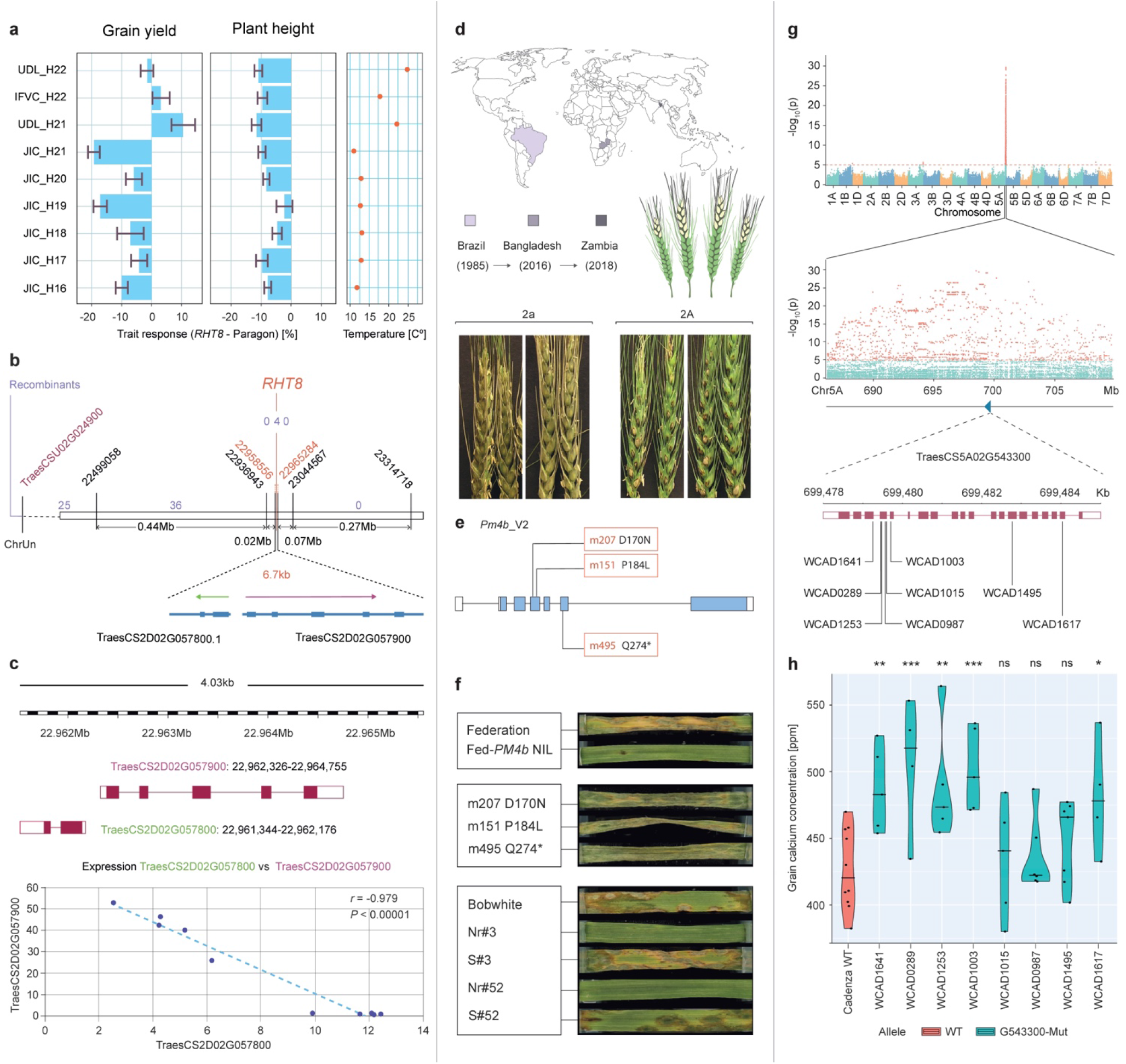
Gene discovery and functional validation for next-generation wheat traits. **a,** Effect on plant height and grain yield (%) of NILs carrying the *RHT8* (Mara) allele compared to the recurrent NIL parent Paragon grown in the UK, Spain and Serbia. H: year harvested, JIC: John Innes Centre (UK), UDL: University of Lleida (Spain), IFVC: Institute of Field and Vegetable Crops (Serbia). Right panel shows the mean temperatures during the growing season at each location. **b,** High-resolution genetic map of the *RHT8* locus showing delimitation to a 6.7 kb interval in the reference genome. Marker positions (bp) are indicated in black, with flanking markers shown in red. Independent recombination lines between the markers and *RHT8* phenotype are indicated in purple. **c,** Two genes (*TraesCS2D02G057800* and *TraesCS2D02G057900*) are present within the 6.7 kb *RHT8* interval in head-to-head orientation. The gene structure is depicted by rectangles (filled: exon; unfilled: untranslated regions) and lines (introns). RNA-seq expression data for the two genes in developing wheat tillers/stems (eight timepoints, three biological replicates per timepoint) reveal a significant negative regulatory relationship (Pearson correlation *r* = −0.979; *P* < 0.0001). **d,** Global spread of wheat blast disease since the 1980s. Bottom panels show the phenotypic effect of *Pm4* resistance (2A) compared to the wild-type non-resistant line (2a). **e,** Diagram of the gene structure of *Pm4* (as in **b**), including the names (red) of the EMS-derived mutants and their predicted effects on PM4_V2 protein (black). **f,** Effects of the *Pm4b* allele on blast resistance in a detached leaf assay in Federation NILs (top). Representative leaves of EMS-derived susceptible mutants (middle) and transgenic lines (Nr#3; Nr#52) alongside non-transgenic wild-type Bobwhite and null controls (S#3, S#52) inoculated with an isolate of the Bangladesh/Zambia B71 lineage of *MoT*. **g,** Watkins NAM GWAS (top panel) for calcium content, identifying a significant signal on chromosome 5A. The most significant SNP corresponds to a haploblock spanning 24 Mb, with the local Manhattan plot highlighted (middle). Detailed map, combined with functional annotation and differential gene expression, showing the structure of the candidate ATPase transporter gene (*TraesCS5A02G543300*), along with EMS-derived mutants in cv. Cadenza (WCAD; bottom). **h,** Grain calcium concentration (ppm) of TILLING mutants (blue) of *TraesCS5A02G543300* along with wild-type Cadenza (pink).

Changing weather patterns have also been linked to increased disease incidence and spread in wheat^32^ and other crops. To identify novel sources of disease resistance, we searched for marker–trait associations for major diseases, including yellow rust, stem rust, *Septoria* and take-all disease (Supplementary Fig. 28). Of the haplotypes extracted from the most significant blocks within the target genomic intervals associated with these diseases, 65% (15,280 out of 23,507) are unique to Watkins landraces (Supplementary Table 46). We focused on wheat blast, caused by *Pyricularia oryzae* (syn. *Magnaporthe oryzae*) pathotype *Triticum* (*MoT*), which poses a significant threat to global wheat production^33^. First identified in Brazil in 1985, the pathogen subsequently spread to neighbouring countries and more recently to Bangladesh (2016) and Zambia (2018), with the 2016 Bangladesh outbreak resulting in over 50% yield losses^34,35^ (Fig. 4d). Therefore, the discovery and deployment of resistance genes against this pathogen are critical for mitigating its threat. Very few sources of wheat blast resistance are available^36^ and despite its importance, no gene has been identified that confers resistance to *MoT*. We screened the core set of the Watkins wheat collection using two *MoT* isolates carrying the *AVR-Rmg8* effector (present in Bangladesh/Zambia isolates; see Methods). An association was observed between resistance to these isolates and a locus on the distal portion of chromosome 2A. O’Hara et al. (https://doi.org/10.1101/2023.09.26.559489) performed a detailed examination of this region using the Watkins data and long-range IBS haplotypes to show that this resistance originated from a wild wheat relative similar to *Triticum turgidum*. The causal gene was identified as *Pm4*, conferring resistance against *Blumeria graminis* f. sp. *tritici*, the powdery mildew pathogen of wheat^37^. The recognition of *AVR-Rmg8* by *Pm4* was confirmed in multiple independent loss-of-function mutants of *Pm4* in the resistant wheat line Federation-*Pm4b* (Fig. 4e) and by the gain of resistance by overexpressing *Pm4* in the susceptible cultivar Bobwhite (Fig. 4f). This is the first report of the identification of genetic variation in a wheat gene that confers resistance to the Bangladesh/Zambia B71 lineage of *MoT*.

Given that wheat is a major staple across many societies, improving nutritional content is a major breeding target for ensuring nutritional food security^38^. To identify nutrition-related genetic variations in the Watkins accessions, we focused on the levels of iron, zinc, potassium, magnesium and calcium in wheat grains: these essential minerals are widely deficient in the human diet. Using three biparental Watkins x Paragon populations, we identified QTL for variation in mineral content which are described elsewhere. Given that grain iron and zinc contents have been extensively studied^39^, we focussed on calcium content in grains, as calcium deficiency is widespread and likely to increase due to the preference for plant-based diets. Calcium is the most abundant mineral in the human body and is required for bone health, healthy pregnancies, cancer prevention, and reducing cardiovascular disease^39^. We located a major gene for grain calcium content on chromosome arm 5AL (Fig. 4g), likely corresponding to a previously identified QTL^40^. We identified loss-of-function mutants for the two most likely candidate genes, *TraesCS5A02G543300* (encoding a cation transporter/plasma membrane ATPase) and *TraesCS5A02G542600* (encoding a major Facilitator Superfamily transporter), with both genes likely carrying loss-of-function mutations in RILs with high grain calcium contents (Fig. 4g). Induced mutations in *TraesCS5A02G543300* (*n* = 8 independent mutants) resulted in a greater than 10% increase in grain calcium content in five mutants (*P value* between 0.03 and 0.002), whereas mutations in *TraesCS5A02G542600* (*n* = 2 mutants) produced no significant changes (*P* = 0.15 and 0.91) compared to control plants (Fig. 4h). This suggests that *TraesCS5A02G543300* and its homoeologues could be manipulated to increase grain calcium content in new wheat cultivars.

By pinpointing *RHT8*, cloning the first resistance gene against the Bangladeshi wheat blast isolate and discovering a novel gene target for increasing grain calcium content, we demonstrated the use of the new Watkins genomic resources for accelerating gene discovery, which will have a direct impact on the breeding of improved wheat cultivars.

### NEW TOOLS FOR THE DEPLOYMENT OF WATKINS VARIATION IN BREEDING

To quantify and accelerate the impact of Watkins diversity on future breeding endeavours, we systematically introduced potentially beneficial Watkins QTL alleles into the Paragon modern wheat genetic background. We successfully introgressed 143 prioritised QTL, 107 of which originate from AGs 1, 3, 4, 6 and 7, which we showed are underrepresented in modern wheat (Fig. 1c, Fig. 5a and Supplementary Table 48). This included 44,338 Watkins-unique haplotypes across all chromosomes available in a stable modern genetic background, thus bridging the gap between landrace diversity and modern breeding. We collected detailed phenotyping data for 11 traits from NIL families (encompassing 127 of these QTL alleles) in multiple field experiments, including evaluation at seven commercial breeding stations for 3–5 consecutive years (Fig. 5b and Supplementary Table 41). Adding allelic effects (*P* < 0.05; Supplementary Table 41) shows the potential to improve various traits in Paragon including (mean Paragon values for each trait are shown in brackets): a 4.5 t ha^−1^ increase in grain yield (9.2 t ha^−1^), an increase of 11,500 grains m^2^ (17,819 grains m^2^), an increase in thousand grain weight of 55.6 mg (44.8 mg), a 32 day variation in heading date, a 52 cm reduction in height (88.4 cm), and a 3.2% increase in protein content (12.4%).

**Fig. 5.**
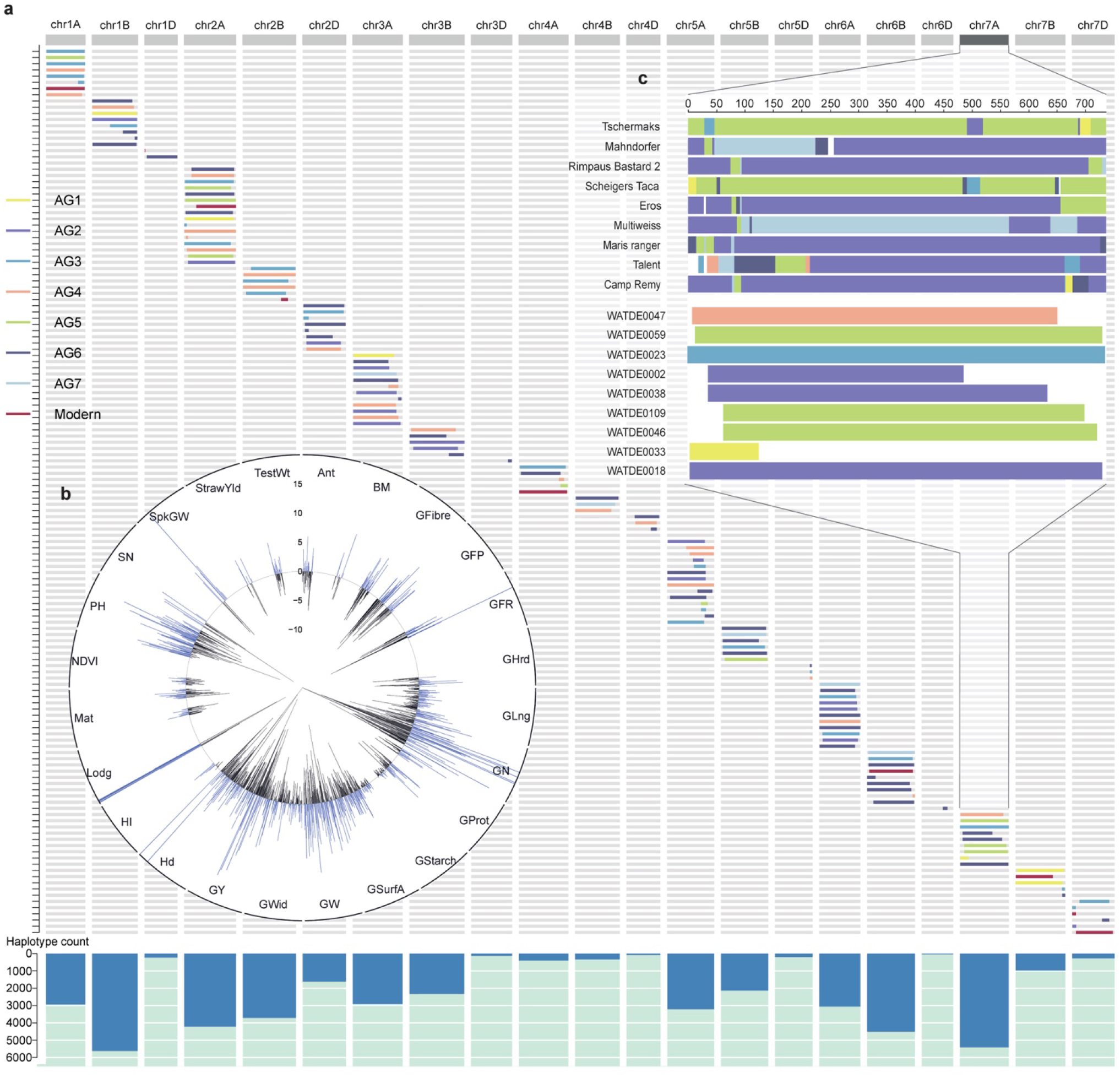
Validation of the breeding value and delivery of target segments. **a,** Introgressed target segments of Watkins in 143 Paragon Near Isogenic Lines (NILs) based on QTL prioritisation. Colours of segments represent the corresponding ancestral groups (AG1–7) of the donor parents. Bar plots at the bottom show the total number of introgressed haplotypes for each chromosome that were previously absent in modern wheat. **b,** Phenotypic effects (AMMI means) of the genetic substitution derived from multi-environment screening for the 127 introgressed loci on 24 traits (marked on the perimeter; for trait abbreviations see Supplementary Table 21). Percentage increase or decrease in Watkins alleles compared to Paragon are shown according to the vertical scale at 12 o’clock. **c**, Inset highlighting one example for chromosome 7A and the NILs (top) with their AG groups and their extent across the chromosome shown. Details on each segment are provided in Supplementary Table 48.

Our comprehensive characterization of modern wheat as a mosaic collection of Watkins IBS segments, combined with the identification of LD haplotypes between Watkins and modern wheat, offers a unique framework for designing and developing new wheat varieties. To enable the selection of any haplotype for this purpose, we identified diagnostic ‘tag’ SNPs along with the corresponding molecular markers for each of the 1.7 million haplotypes (Supplementary Table 49). To demonstrate the efficacy and utility of these markers, we selected tag markers for two QTL: 5A awn inhibitor (*B1*)^41^ and 7A coleoptile colour (*Rc*) (Supplementary Fig. 29). Using KASP assays, we genotyped 382 independent accessions from the BBSRC small grain cereals collection (Supplementary Table 50). These markers proved to be highly effective for enriching target haplotype selection, resulting in correct predictions of the two phenotypes in 87.4% and 93.7% of accessions, respectively.

In addition to marker-assisted selection, the combined Watkins/modern dataset and haplotype analysis maximise the discriminatory power of high-throughput genotyping arrays. This is particularly valuable for germplasm that has not undergone full sequencing, as would usually be the case for genomic selection (GS) or GWAS within breeding programmes. We therefore developed a *Triticum aestivum* Next Generation genotyping array (*Ta*NG). To test the performance of this new array, we compared the Watkins GWAS results for four traits: heading date, stem rust, coleoptile colour and awn length, using (i) 10 million core SNPs generated from our whole-genome sequencing, (ii) the existing 35K Axiom array and (iii) the *Ta*NG array (Supplementary Fig. 30). Notably, the *Ta*NG array identified three MTA not identified using the 35K Axiom array, showcasing the enhanced capability of the *Ta*NG array for trait-gene discovery (Supplementary Table 51).

## DISCUSSION

We cannot overstate the urgency of breeding climate-resilient crops for future food security and therefore the importance of discovering (or re-discovering) and characterizing genetic resources that improve crop performance under challenging environmental conditions. For wheat, Arthur Ernest Watkins first described the bread wheat landraces used here in “The wheat species: a critique” published in 1930^42,43^. Although nearly a century has passed, we can now use genomics to fully realise the potential of these invaluable genetic resources. Indeed, our whole-genome re-sequencing of the Watkins landrace collection showed that five of the seven Watkins AGs are phylogenetically isolated from modern wheat varieties (Fig. 1b). By combining structured gene discovery populations, genomic data and extensive field trials with in-depth phenotyping, we confirmed that many of the haplotypes restricted to phylogenetically isolated Watkins AGs have beneficial effects on yield potential, adaptation, human nutrition and disease resistance. Development of the first wheat haplotype map and haplotype–phenotype association study allowed us to estimate frequencies of these alleles and assess their functional significance. To introduce novel and useful landrace diversity into the landscape of modern wheat breeding, we transferred 44,338 Watkins-unique haplotypes from Watkins into a single elite wheat variety. The confirmed breeding values help guide the stacking of beneficial Watkins haplotypes into cultivars for further evaluation across environments. This comprehensive strategy will enable breeders to leverage this untapped genetic diversity in crop improvement to help secure sustainable food production, which is achievable because genomic variants of global wheat varieties directly affect soil, water, air, and nutrition.

Our findings establish the Watkins resources as a powerful, unique platform for isolating genes controlling in-demand traits and exploring genetic regulation in modern varieties. We envision a future where wheat crops will be genetic mosaics that include AGs 1, 3, 4, 6, and 7 (Fig. 5c), and thus have increased yield potential and reduced reliance on nitrogen fertilisers, thereby lowering the main source of greenhouse gas emissions from agriculture (Fig. 3g and 5b). Moreover, breeders can deploy new arsenals of disease resistance genes to combat evolving pathogen populations driven by the changing climate. For example, here we identified new sources of resistance to aggressive yellow rust isolates (Supplementary Table 46) and the Watkins resources helped clone a novel gene for *Septoria* resistance^44^. We also provide novel mechanistic insights into the genetic control of wheat blast through the cloning of the first wheat gene conferring resistance to the devastating Bangladesh/Zambia *MoT* isolate (Fig. 4d-f). In addition, nutritious wheat varieties will facilitate dietary shifts towards plant-based diets, with the associated reduction in livestock greenhouse gas emissions (Fig. 4h). Moreover, the phenotypic versatility endowed by novel variation in canopy architecture and phenology traits offers the potential for new wheat varieties that are more resilient to heat and drought (Fig. 4b and Supplementary Figs. 23-26). Ultimately, these discoveries demonstrate the transformative potential of these resources, which will enable breeders to address the urgent challenge of adaptation to climate change in wheat.

To empower the global community to accelerate breeding in wheat, we have adhered to the collaborative spirit of the Human Genome Project by making our resources, including germplasm and genomic and phenotypic data, publicly available through the Watkins Worldwide Wheat Genomics to Breeding portal (https://wwwg2b.com/). We aim to promote openness and collaboration that will enable the full potential of this work to be realised, providing resources for extending the use and further development of the Watkins resource, for example, through developing pan-transcriptome maps, gene regulatory atlases or pan-structural variation studies.

Our analysis of this remarkable genetic resource from the 1900s underscores the enduring value of collecting and preserving genetic diversity *ex-situ*. The Watkins collection, assembled from across the globe a century ago, is now reverting to its global role once more. The profound legacy left by Watkins and others has inspired our international collaboration and commitment to open data sharing and knowledge exchange, recognizing the collective benefits to the global community^45^. Although recent policies have restricted international germplasm exchanges^46,47^, it is crucial to remember that the challenges posed by climate change transcend these artificial boundaries.

## Data availability

All whole-genome sequence data has been deposited at the National Genomics Data Center (NGDC) Genome Sequence Archive (GSA) (https://ngdc.cncb.ac.cn/gsa/), with BioProject accession number PRJCA019636. Variation matrix and annotations, wheat HapMap, phenotyping data, association genetics analyses, the developed tagSNPs and KASP markers were deposited in WWWG2B breeding portal (https://wwwg2b.com). IBSpy variations tables, haplotypes, long-range tilling paths, variant files (VCF) and all raw phenotypic data are available online (https://opendata.earlham.ac.uk/wheat/under_license/toronto/WatSeq_2023-09-15_landrace_modern_Variation_Data/). All the Source Data to support all the Figures and provided in the attached file as well as in: https://doi.org/10.5281/zenodo.8351964. Publicly available sequencing data were obtained from SRA accessions SRP114784, PRJNA544491, PRJEB37938, PRJNA492239, PRJNA528431, PRJEB39558, PRJEB35709 and from the NGDC database project CRA005878.

## Germplasm availability

The 827 Watkins single seed derived accessions and their 827 predecessor landrace populations, 208 modern cultivars, 73 RIL mapping populations and 143 NIL families are all available from the John Innes Centre Germplasm Resources Unit (https://www.seedstor.ac.uk/) and the Agricultural Genomics Institute at Shenzhen, Chinese Academy of Agricultural Sciences (https://wwwg2b.com/).

## Code availability

Code associated with this project is available at Github: https://github.com/ShifengCHENG-Laboratory/WWWG2B; https://github.com/Uauy-Lab/IBSpy; https://github.com/JIC-CSB/WatSeqAnalysis/; and https://github.com/pr0kary0te/GenomeWideSNP-development.

## Acknowledgements

We thank Prof. Graham Moore and Prof. Mike Bevan for providing valuable feedback at multiple stages of the project. We thank colleagues for assistance in Watkins field trial and phenotyping work from five experimental stations across China, Dr. Zhanwang Zhu (Hubei (ErZhou) Academy of Agricultural Sciences), Dr. Qiang Wang (HeiLongJiang (HaerBing) Academy of Agricultural Sciences), Yinming Song (Hebei (Quzhou) agricultural experimental station), Prof. Yan Zhu (Nanjing Agricultural University) and Prof. Xiansheng Zhang (Shandong Agricultural University). We thank the John Innes Centre (JIC) NBI Computing Infrastructure for Science group, the JIC Field Trials and Horticultural Services teams for support in field and glasshouse experiments, Tobin Florio at Flozbox studio (https://flozbox-science.com/) for Figure visualisation, and the Rothamsted Research farm team and Analytical Chemistry unit for support in field experiments and analytical mineral analyses.

This work was supported by the Program for Guangdong “ZhuJiang” Introducing Innovative and Entrepreneurial Teams (2019ZT08N628), the National Natural Science Foundation of China (32022006), the Agricultural Science and Technology Innovation Program (CAAS-ASTIP-2021-AGIS-ZDRW202101), the Shenzhen Science and Technology Program (AGIS-ZDKY202002) to S.Cheng, and the Guangdong Basic and Applied Basic Research Foundation (2020A1515110677) to L.M. The UK work was possible due to the long term investment of the UK Biotechnology and Biological Sciences Research Council (BBSRC) in wheat research through Institute Strategic Programme (ISP) grants and longer larger grants: BBSRC LOLA ‘Enhancing diversity in UK wheat through a public sector prebreeding programme’ (BB/I002545/1); BBSRC ISP ‘JIC WISP ISP - Wheat Institute Strategic Programme’ (BB/J004596/1); BBSRC ISP ‘BBSRC Strategic Programme in Designing Future Wheat (DFW)’ (BB/P016855/1); BBSRC ISP ‘BBSRC Institute Strategic Programme: Delivering Sustainable Wheat (DSW)’ (BB/X011003/1) and for wheat germplasm conservation and global distribution through the Germplasm Resources BBSRC National Capability award (BBS/E/J/000PR8000). S.G. and C.L. also received support from the UK Department for Environment, Food and Rural Affairs (Defra) as part of WGIN phases 3 and 4 (CH0106 and CH0109). This work was also supported by the European Research Council (ERC-2019-COG-866328), the Sustainable Crop Production Research for International Development (SCPRID) programme (BB/J012017/1), the Mexican Consejo Nacional de Ciencia y Tecnología (CONACYT; 2018-000009-01EXTF-00306), the Science, Technology & Innovation Funding Authority (STDF), Egypt-UK Newton-Mosharafa Institutional Links award, Project ID (30718) and EG - US cycle 19 - Project ID (42687).

## Author contributions

S.Cheng and S.G. conceived, designed, coordinated and managed the project; S.Cheng led the genomics, bioinformatics, association genetics analysis and phenotypic activities in China; C.F. led population genomics, association genetics and bioinformatics analyses under S. Cheng’s supervision, including variants calling and quality control, construction of the LD-based haplotype map, GWAS, NAM imputation and NAM GWAS, integrated association genetics and quantification of genetic effects and haplotype-phenotypic associations, the development of HAPPE pipelines and the WWWG2B portal, tagSNP and KASP markers development; L.U.W. led quantitative genetics analyses including genetic mapping, QTL mapping, statistical analysis of field trials including AMMI and trait ontology; L.U.W., A.B.R., M.L-W., M.J.H. and S.G. led the field trial, phenotyping activities and data analyses in the UK; H.C. and M.J. assisted in the bioinformatics analyses including variant calling, LD-based haplotype construction, GWAS and NAM GWAS; Z.H. built the WWWG2B portal and provided additional computational support; S.Collier, S.O., and R.A. developed germplasm, performed marker assisted selection and curated seed stocks; G.B., A.P-A, M.W. and K.J.E. designed the TaNG genotyping array; T.O. and P.N. designed, performed and analysed wheat blast experiments; C.L., A.K. and S.G. led the fine-mapping of *RHT8*; B.B., A.K-S., G.S. and S.G. conducted field trials for *RHT8*; A.S., N.C, and L.U.W. led the grain calcium content phenotyping; J.Q-C., R.H.R-G. and C.U. developed IBSpy and with Xiaoming Wang performed the long-range haplotype analyses; R.B. and S.B. coordinated field and glasshouse disease phenotyping in commercial settings; L.D and A.G. performed flowering time gene analyses; L.B. extracted DNA for sequencing; A.B., M.L-W. and K.J.E. conducted high density genotyping; K.G. and Xingwei Wang supported GWAS analyses; H.C., X.L. and Bo Song performed population structure analyses; W.X. led CNV identification; Ba.Song and H.L. provided bioinformatics support; R.Horler and S.T. digitised germplasm, genomic and phenotypic data; B.Steuernagel led bioinformatics work in UK; U.B., H.S.B., J.K.M.B., C.B., M.B., L.C., A.F.E., P.F., D.F., A.N.H., M.S.I., M.K., W.D., J.L., S.L., D.Schafer, P.T., M.N.W., B.B.H.W. and Q.Y. conducted disease phenotyping and data analyses; M.J.B., M.C., B.C., J.F., O.G., R.Han, J.H., Z.H., L.M., C.M., S.P., C.P., D.P., Z.S., Y.S., D.Steele, A.W., Y.Wei, S.W., Y.Wu, Yawen Xu, Yunfeng Xu and X.Z. conducted phenotyping experiments; A.L. and P.R.S. conducted grain phenotyping experiments; N.C. managed germplasm and coordinated phenotyping of grain traits; D.Sanders and S.H. provided leadership and coordination roles; S.Cheng, C.F., L.U.W, Xiaoming Wang, H.C., M.J., C.U. and S.G. prepared the Figures, Extended Data Figures, Supplementary Figures and Supplementary Tables; S.Cheng., C.F., L.U.W., C.U. and S.G. wrote the manuscript, with additional help from N.C and A.B.R.; All authors read and approved the final manuscript.

## Additional information

Correspondence and requests for data and materials should be addressed to.

## Online Methods

### Statistical analysis

Statistical analyses were conducted in R software suite (version 4.2, https://www.r-project.org/) unless otherwise stated. The Linkage disequilibrium (LD) and haploblocks were calculated by PLINK (version 1.90 beta)^49,50^, the haplotype clustering was performed by HAPPE^18^. The phenotypic effects observed in the NILs were assessed using ANOVA (Analysis of variance) or by AMMI (Additive Main-effects and Multiplicative Interaction) mean values. Where relevant, statistical tests were two-sided, randomised experimental units were used as replications, multiple measurements of single experimental units were treated as subsamples, and all data was tested for assumptions and corrected accordingly, and described.

### Germplasm collection and glasshouse growing conditions

The 827 accessions from the entire A.E. Watkins Collection of landraces were used, alongside 208 modern wheat lines selected to represent worldwide diversity and to include parents of publicly available populations^13^. Seeds were obtained from the John Innes Centre Germplasm Resource Unit (JIC GRU, http://www.jic.ac.uk/germplasm/). Single seeds were sown in 3.8 cm diameter pots containing peat and sand mixture (85% fine grade peat, 15% washed grit, 4 kg m^-3^ Mag Lime (powdered limestone containing 90% CaCQ3), 2.7 kg m^-3^ slow release fertiliser (Osmocote, 3–4 months), 1 kg m^-3^ PG Mix 14-16-18 + TE 0.02% and wetting agent) for 3 weeks growth at 20/17.5 °C day/night temperature with 16 h day length, before transfer to 6 °C day & night temperature with 8 h day length for 8 weeks.

Following vernalization, plants were transferred to outdoor glasshouse conditions with automated watering and following 2 weeks further growth, transplanted into 2 L pots containing cereals mixture (40% medium grade peat, 40% sterilised loam (soil), 20% washed horticultural grit, 3 kg m^-3^ Mag Lime (powdered limestone containing 90% CaCQ3), 1.3 kg m^-3^ PG mix 14%-16%-18%, N-P-K + TE base fertiliser, 1 kg m^-3^ ‘Osmocote mini’ 16%-8%-11% 2 mg + TE 0.02%, and wetting agent) for continued development. During plant growth, spikes were bagged to prevent cross pollination and plant material dried naturally. Seeds were deposited in JIC GRU and transferred to the Agricultural Genomics Institute at Shenzhen (AGIS), Chinese Academy of Agricultural Sciences.

### DNA Extraction

Genomic DNA was extracted from approximately 50 mg wet weight young leaf tissue of three-week stage seedlings. Extractions for the Watkins collection used the DNeasy 96 Plant Kit protocol (Qiagen) and extractions for remaining lines used the oKtopure^TM^ automated plant-based system (LGC Biosearch Technology) following tissue desiccation with silica for 48 h. A bespoke maxi prep protocol was used with the following volumes per sample: 250 µl lysis buffer, 170 µl Binding buffer, 20 µl sbeadex^TM^ suspension, 300 µl PN1 wash buffer, 300 µl PN2 wash buffer, 300 µl PN2 wash buffer (x3 wash cycles) using 75 µl final Elution buffer.

### Whole Genome Resequencing and Quality Control

DNA was used for sequencing library construction following the manufacturer’s protocols (Illumina Inc.). Libraries were 150 bp paired-end with insert size ∼500 bp and were sequenced on an Illumina NovaSeq 6000 at Berry Genomics (954 accessions), Beijing, in 2018, and also on DNBSEQ Platform at BGI group (90 accessions). A total of over 200 TB raw data was generated, producing, on average, 193 Gb raw reads for each accession (Supplementary Fig. 1 and Supplementary Table 1). Then, we used the following parameters (fastp^53^ v0.20.0: -f 9 -F 9 -l 80 -g) to filter the raw data: (1) Remove adaptor sequences; (2) Discard reads of N number >=5; (3) Discard reads of base quality <=15 exceed 40%; (4) Trimmed 9 bp in the front of reads; (5) Retain the reads length >=80bp; After removing the low-quality and adaptor-containing reads, an average of ∼185.14 Gb of clean data (∼12.73X coverage) was retained for each accession.

### Reads Mapping, Variant discovery, quality control and SNP annotation

The clean reads were mapped to IWGSC RefSeq v1.0 using BWA-MEM (v0.7.17)^51^ with default parameters. Non-unique mapped and duplicated reads were excluded using SAMtools (v1.9)^52^ and Picard (v 2.20.3-SNAPSHOT)^53^, respectively. SNP and InDel calling were performed by GATK (v4.1.2)^54^. A total of 720,048,179 raw variants (SNP/InDel) were identified from GATK, including 668,764,660 SNPs and 51,283,519 InDels.

For variants filtering and Quality Control process, there are four main steps for both SNPs and InDels, corresponding to Supplementary Table 2. Firstly, only the bi-allelic variants were retained, including 634,873,707 SNPs and 44,525,746 InDels. Secondly, variants were filtered based on the parameters recommended by GATK. SNPs Filtering criteria: QD < 2.0 || FS > 60.0 || MQ < 40.0 || MQRankSum < -12.5 || ReadPosRankSum < -8.0 || SOR > 3.0. Indels Filter criteria: QD < 2.0 || low_QD || FS > 200.0 || high_FS || ReadPosRankSum < -20.0 || low_ReadPosRankSum. A total of 411,400,604 SNPs and 42,415,907 InDels were retained using such filtering criteria. Third, variants were filtered by Inbreeding Coefficient (F). F is computed as: F= 1-(Hobs/Hexp). Hobs is the frequency of heterozygous calls, and Hexp = 2p(1-p), in which p is the frequency of non-reference allele (or reference allele). The median value of F (Fmedian) for each chromosome is calculated using the SNP site of F>0 and minor allele frequency (MAF) >0.05. The max observed heterozygous frequency (Hobs_max) is computed as: Hobs_max = 10*(1-Fmedian)*Hexp. The sites of Hobs>Hobs_max were discarded. A total of 261,659,890 high-quality SNPs (dataset 1) and 17,279,131 high-quality InDels were retained, which is typically for association genetics study. Finally, the SNPs with missing rate >20% and MAF < 0.01 were discarded. A total of 90,750,089 common SNPs (high frequency) were retained as high-quality SNP dataset (dataset 2), which is specifically for population LD-based analyses.

SnpEff (v4.3t)^55^ was used for annotating and predicting the genome structural position and functional effects of identified SNPs and InDels. SNPs were annotated as exonic, intronic, splicing region, up/down stream, intergenic and 3’/5’ prime untranslated region (UTR) variants. Exonic variants can be further divided into synonymous and nonsynonymous variants, and the latter included missense variants, stop loss, stop gain, start loss, start gain, and stop retained. Intron variants can be categorised as splicing donors, splicing acceptors and others.

### Identification of gene Copy Number Variations (CNVs)

Considering the limitations of using short reads for CNV identification, only the CNVs of the wheat protein-coding genes were calculated in this study. Five steps were implemented for the identification of gene CNVs. (1) Calculation of read depth for each gene, sequenced in each accession, based on the properly mapped reads. This is referred to as the absolute read depth for each gene. (2) Optimal correction of the absolute value for read depth variation (RDV). To account for highly similar genes in the reference genome, such as paralogues arising from recent duplication events, we performed an all-vs-all CDS alignment using BLASTN. Genes that meet specific criteria (fewer than 5 gaps and fewer than 5 mismatches) are classified as recently duplicated. For depth calculations, these highly similar genes are collapsed into a single representative gene in the reference genome. Specifically, the depth values of all the duplicated genes are summed together. This approach aims to minimise the depth bias introduced by recent gene duplications in the reference genome^56^. (3) Normalization for each accession. Considering the slight variation for the sequencing depth for each accession, we divided the corrected read depth for each gene in step (2) by the average sequencing depth for each accession. This is the relative read depth variation. (4) GC bias correction. There were GC bias in Illumina short-read sequencing technology despite all libraries being PCR-free. We gave a read depth distribution for all the GC content within the wheat genome (1,000 windows for GC content from 0% to 100%) and corresponded the estimated read depth for each level of GC content for each gene to this distribution, which was divided by the overall read depth of the overall GC content. This resulted in the GC content correction value for each gene, to avoid and correct the GC bias for read depth^57^. (5) Correction of read depth variation for genomic regions with Insertions or Deletions in the genome reference. To explore the population-unique and shared CNVs, the number of accessions with different copy numbers, such as [0-0.25), [0.25-0.75), [0.75-1.25), were calculated for each gene in both Watkins and modern populations.

### Ensuring correspondence of sequenced accessions with existing public genotypic data

To ensure consistency of sequenced samples, sequence data was compared to publicly available genotyping data using the Axiom 35k Breeders’ array^58^. The probe set of the Breeders’ array (https://www.cerealsdb.uk.net/cerealgenomics/CerealsDB/Excel/35K_array/35k_probe_set_IWGSCv1.xlsx) was used to generate two allele-specific sets of *k*-mers (*k*=31) for each SNP of the array. We also built *k*-mer databases from the 12-fold sequence data of each Watkins line. The allele for each SNP in each Watkins line was determined using presence or absence of allele specific *k*-mers in the *k*-mer databases. This resulted in a genotype profile for each Watkins line based on the 12-fold sequence data. Next, the new genotype profile was compared to existing data generated using the 35k Breeders’ array, which provides an indication if samples we sequenced are consistent with accessions genotyped in previous studies. From this analysis, we obtained 100% match between the sequenced accessions and the existing public genotypic data. Scripts for the procedure and details on the pipeline are available at https://github.com/JIC-CSB/WatSeqAnalysis/tree/master/qc_vs_iselect.

### Linkage disequilibrium (LD) analysis and Construction of the Wheat Haplotype map (HapMap)

To obtain a core SNP set from the filtered SNP set described above, a two-step LD pruning procedure were conducted as previously done in rice^59^. First, SNPs were removed by LD pruning with a window size of 10 kb, window step of one SNP and r^2^ threshold of 0.8 using PLINK^49^. Second, another round of LD pruning with a window size of 50 SNPs, window step of one SNP and r^2^ threshold of 0.8 was performed. About 10 Mb SNPs retained after two-step LD pruning (dataset 3). The construction of wheat HapMap included two parts, the population linkage disequilibrium (LD) based haplotype map, and the Identity-by-State in python (IBSpy) *k*-mer based large-scale long-range haplotypes segments^16^.

*Population LD-based haplotype analysis*. First, the SNP dataset 2 was phased by Beagle (v 21Apr21.304)^60^. Using this phased dataset, haplotype blocks were identified using PLINK^49^ with the parameters (--blocks no-pheno-req --blocks-max-kb 1000 --geno 0.1 --blocks-min-maf 0.05). To merge the adjacent blocks that might still maintain strong linkage disequilibrium into larger ones, the D’ statistic value were calculated for all of the SNPs (dataset 1) of every two adjacent blocks. If the lower quartile (Q1) was larger than 0.98, the two adjacent blocks would be merged. HAPPE^18^ that we developed previously was used to identify haplotype clusters (haplogroups) for each block based on SNPs dataset 1.

*k-mer based IBS long-range haplotype analysis*. We devised a systematic approach for conducting *k*-mer based IBS approach for long-range haplotype analysis and reconstructed modern wheat genomes using the Watkins lines. The methodology comprises the following steps:

1. Generation of *k*-mer matrices and variation analysis: We initiated the analysis by utilizing the kmerGWAS pipeline^61^ (https://github.com/voichek/kmersGWAS) to produce a *k*-mer matrix (*k* = 31) for our dataset containing 1,051 wheat accessions. Concurrently, we integrated the chromosome-level genome assemblies from the 10+Genome project, including Arina*LrFor*, Chinese Spring, Jagger, Julius, LongReach Lancer, CDC Landmark, Mace, Norin 61, CDC Stanley, and SY Mattis. Employing the IBSpy pipeline (https://github.com/Uauy-Lab/IBSpy), we computed the *k*-mer variation matrix (*k* = 31) for each reference assembly vis-à-vis the 1,051 accessions, utilizing non-overlapping 50 kb step windows.
2. Transformation of *k*-mer variations to IBSpy values and haplotype assignment: Next, we converted the *k*-mer variation matrix into IBSpy *variation* values. We then conducted haplotype assignment for distinct non-overlapping 1 Mb windows. We applied the affinity propagation clustering technique^62^ (with a window size of 1 Mb) based on IBSpy *variation* values computed from 20 consecutive 50 kb windows. To enrich the IBSpy values utilised in clustering, we incorporated IBSpy values derived from syntenic regions across the 10 genome references.
3. Pedigree tracking and genome reconstruction: Pedigree tracking was employed to track the ancestry of each modern cultivar within the Watkins collection. Building upon the haplotype assignments in non-overlapping 1 Mb windows, we reconstructed each cultivar’s genome using extended matched haplotype blocks from the Watkins lines, employing an *in silico* strategy featuring a minimum tiling path approach.

**a.** Haplotype Comparison and Identification of Progenitor: On a per-chromosome basis, we began by comparing the haplotypes of a cultivar’s first window with the corresponding chromosome’s haplotypes in each Watkins line. The length of Matched Haplotype Windows (MHW) from each Watkins line was noted. The Watkins line with the longest MHW was identified as the potential progenitor contributing to that genomic region. Subsequent MHWs from other Watkins lines were discarded.
**b.** Iterative Process and Longest MHW Recording: The comparison process was iterated from the second window onward. At each step, we identified the longest MHW and its associated Watkins line. This iterative comparison continued window by window until covering the entire chromosome.
**c.** Refinement and Minimum Tiling Path Construction: We then aligned the physical positions of identified MHWs from each window. Overlapping MHWs were removed, retaining those forming the minimum tiling path for reconstructing the chromosome. The reconstructed path allowed observation of the Mosaic composition originating from the Watkins lines. This strategic process delineates the closest Watkins relatives at a 1 Mb resolution for any given genomic region within a modern cultivar. The contribution percentage of each Watkins line to a cultivar was quantified as the ratio of its total MHWs to the entire genome windows.

### Population structure analysis

*Phylogenetic tree and ADMIXTURE.* For the Phylogenetic genetic analysis, neighbor-joining trees was constructed for the genome-wide 4DTv (fourfold degenerate synonymous Site), and tagSNPs (described below) using rapidnj^63,64^. 1000 bootstrap were performed for each tree. Interactive Tree of Life (iTOL) was used to visualise and annotate these trees (https://itol.embl.de/)^65^. To explore and obtain the accurate population structure of Watkins collection, a new pipeline was developed with a view to pervasive introgression in wheat genome. First, we calculated the genetic distance of each pairwise accessions in 5 Mb sliding window (dataset 2) using vcf2dis software (https://github.com/BGI-shenzhen/VCF2Dis). We also calculated the geographic distance of each pairwise accessions based on the longitude and latitude using R package geosphere. Then, the correlation between the genetic and geographic distances was calculated for each 5 Mb window using R package corr. R value of 88.97% of introgression block with known resource is less than 0.07, indicating that R value = 0.07 will be powerful to exclude the introgression. Therefore, the SNPs located in the windows with R value less than 0.07 were discarded to reduce background noise. The remaining SNPs were used to quantify the genome-wide population structures using ADMIXTURE^66^.

*t-SNE and PCA analyses*. For the haplotype matrix, we first imported the data into a Python environment and then transformed the matrix into a one-hot encoded format using the OneHotEncoder class from the sklearn. preprocessing module. Subsequently, we used the PCA class and the TSNE class from the sklearn.decomposition and sklearn.manifold modules, respectively, to perform PCA and t-SNE. For the SNP matrix, VCF datasets were converted into a numerical matrix. In this conversion, a value of ’0’ represents a reference allele, ’1’ represents heterozygosity, ’2’ indicates an alternative allele, and ’-1’ is used for missing data. We then applied the PCA and TSNE methods as described above.

### Genetic diversity and population differentiation

In consideration of the effect of the missing rate and the MAF on genetic diversity, dataset 1 (without missing rate and MAF filtering) was used in the following analyses. We calculated the number of SNPs, number of InDdels and π of accessions in different populations and countries. These calculations were performed in non-overlapping 2 Mb windows across the whole genome using PLINK. For genic diversity, we calculated the number of total SNPs, population-unique SNPs and allele frequency among different populations within each gene. To evaluate the populations differentiation among Watkins groups and modern variety, we further used plink--cluster to calculate the identity by state distance of each pairwise accessions ^49^. The allelic diversity, haplotype clustering and cataloguing, and CNV diversity according to the read depth variations were analyzed and visualization with HAPPE^18^.

### Field Experiments for Watkins Collection

We conducted field experiments to assess the phenotypic diversity of the Watkins collection (the natural population, Supplementary Fig. 19).

*UK Field Experiments for Watkins Collection.* The Watkins collection was grown at the JIC Field Experimental Station (JI) (Bawburgh, Norfolk; 52.628 N, 1.171 E) in 2006 in unreplicated 1 m^2^ plots under low nitrogen input as previously reported^67^. Experiments were repeated at JIC in 2010 and 2014. Phenotypes measured were: 2006: heading date, plant height and growth habit; 2010: presence of awns, heading date, thousand grain weight, grain width and grain length; 2014: heading date, kernel hardness and coleoptile colour. Details on phenotype measurements are given in crop ontology format in Supplementary Table 21.

*Chinese Field Experiments*. The Watkins collection, alongside 208 contemporary cultivars, were grown and phenotyped in diverse geographic locations throughout China. These sites were: Shenzhen city (22.597N, 114.504E, seasons 2021, 2022 and 2023), Guangdong Province, southern China; Ezhou (30.386N, 114.656E, season 2023), Hubei Province, middle of China; Nantong (32.268N, 120.759E, season 2023), Jiangsu Province, southeast China; Tai’an (35.987N, 116.875E, season 2023), Shandong Province, northern China; Quzhou (36.863N, 115.016E, season 2022), Hebei Province, the northern; and Harbing (45.830N, 126.853E, season 2023), HeiLongJiang Province, northern China. All trials were hand planted in the autumn (mostly November), with the exception of Harbing, where sowing took place in March. Plants were grown in 1.2 m or 2 m long rows using an augmented plot design with 50 or 100 plants per block with three check varieties, with the exception to Shenzen 2020 and Ezou 2022 where a factorial split-block design with two Nitrogen treatments and two replicates were grown. Plants were phenotyped for a broad range of traits spanning lodging, height, tillering, phenology, disease resistance, and various morphological traits, alongside yield and biomass components. Following harvest, yield component traits including spikelet number and grain morphometric traits were measured.

*Egyptian Field Experiments for Watkins Collection.* A diverse subset of 300 Watkins bread wheat landraces and 20 modern lines were grown at four agricultural research stations in Egypt: Sakha (31.0642 N, 30.5645 E), Nubaria (30.6973 N, 30.66713E), Gemmiza (30.867 N, 31.028 E) and Side (29.076 N, 31.097 E). Growing season was from November to the end of May with harvest in the years 2020, 2021 and 2022. The wheat lines were grown in 3.5 m rows and hand harvested. Fertiliser applications were before sowing: phosphorus (200 kg P ha^-1^) and potassium sulphate (50 kg K ha^-1^); and three doses of Urea (in total 300 kg N ha^-1^) at sowing, 30 days post-sowing, and at the tillering stage. Phenotypes recorded (as the mean of ten plants) were: growth habit, plant height, heading date, number of kernels per spikes, grain weight, maturity date. Rust scores were taken at the early dough stage as host responses and rust severity.

### RIL Population Development and Analysis

*Construction of bi-parental populations.* Bi-parental populations with diverse Watkins landrace parents were developed as described^8^. Initial crosses of Paragon (female) to Watkins landrace (male) plants were advanced to F4 using single seed descent. In total, we developed 109 populations using 107 different Watkins accessions, resulting in 9,771 recombinant inbred lines (RILs). Of these, 73 RIL populations have been phenotyped to date (Supplementary Table 20 and https://www.seedstor.ac.uk/search-browseaccessions.php?idCollection=47).

*Genotyping RIL Populations*. Early in the project, genotyping was conducted with KASP and SSR markers followed by genetic map construction as described. This was the case for the majority of the populations, see Supplementary Table 20, column ‘Genetic Map Type’. Later, genotyping was done with the 35k Axiom Wheat Breeders’ array^13^.

*Genetic Map construction.* Genetic map construction was carried out using the R package “ASMap” (version 1.6) as described^68^. The genotype scores can be retrieved from CerealsDB (https://www.cerealsdb.uk.net/cerealgenomics/CerealsDB/array_info.php)

*QTL discovery.* QTL mapping was conducted in the R software suite (v3.6.1) using package “qtl” (v1.5)^69,70^ as described taking cross type (RIL) and generation number (F4 or F5) into account. The QTL model used a significance threshold calculated from the data distribution. A first QTL scan, using Haley-Knott regression, determined co-factors for the second scan. The second scan by composite interval mapping (CIM) identified QTL at a significance level of 0.05 taking the co-factors into account. The resulting QTL with a LOD score equal or larger than 2.0 are listed in Supplementary Table 25.

### Field experiments using RIL populations

Trials were drilled around mid-October and harvested end of July to late August and grown with standard farm management^71,72^. 73 bi-parental RIL populations were grown in field experimentation trials over ten years between 2011 and 2020 at the JIC Field Experimental Station (JI) (Bawburgh, Norfolk; 52.628 N, 1.171 E) in randomised un-replicated 1 m^2^ multiplication trials with low Nitrogen input (approx 50 kg N * ha^-1^). A subset of 18 populations were grown in a randomised block design (3 replicates, 1 m^2^ plots) at Rothamsted Experimental Farm (RH) in south-east England (51.8100 N, -0.3764 E) at two different Nitrogen levels over 7 years (2012-2018). Nitrogen supply was taken to be the sum of the amount of mineral nitrogen in the top 90 cm of soil measured late winter each year (before the first fertiliser N application) and the amount of N fertiliser applied (either as ammonium nitrate or ammonium sulphate). Nitrogen fertiliser applications (50 or 200 kg N ha^-1^) were made late-February to May, with a split application to plots receiving 200 kg N ha^-1^. A set of fifteen populations were grown at Bunny Farm, University of Nottingham, Nottinghamshire, (SB) (52.8607 N, -1.1268 E) in randomised replicated 1 m^2^ plots under two different Nitrogen conditions as in RH over four years (2012-2015). Details on which population was grown in which season are given in Supplementary Table 23. Seed sources for the JI trials were glasshouse seed, and for the other trials the JIC field multiplied seeds. Field experiments at RH were targeted to specifically assess grain yield and NUE. Above-ground nitrogen uptake was calculated from the sum of nitrogen in straw and grain at final harvest. These data were calculated from grain and straw dry matter yield; both recorded by plot combine harvester at Rothamsted, grain by combine and straw from a pre-harvest grab sample at Nottingham. Harvest index (ratio of grain dry matter to above-ground dry matter) was also calculated, and grain and straw nitrogen concentration, measured on samples taken at final harvest and measured by near Infrared spectroscopy using in-house calibrations. Nitrogen use efficiency (NUE) was calculated using published methods^72^. Overall, phenotypes recorded were: JI: heading date, plant height, grain yield, grain weight, grain length, grain width, and grain surface area; RH and SB: anthesis date^73^, crop height, grain yield, straw yield, grain and straw nitrogen concentration. RH also carried out canopy maturity, grain and straw moisture content and grain weight measurements and on targeted populations mineral analysis by Inductively coupled plasma optical emission spectrometry (ICP-OES). Further phenotypes were calculated from the direct measurements, e.g. grain number, harvest index, nitrogen uptake, NUE, grain fill rate, grain protein concentration and grain protein deviation. Details on phenotype measurements are given in crop ontology format in Supplementary Table 21. Four RIL populations targeted for yellow rust resistance were grown in six experiments by commercial partners as winter drilled, unreplicated 1 m^2^ plots and scored for yellow rust incidence. Locations were: Limagrain UK (Rothwell Cherry Tree Top, 53.4882N, -0.25437E), RAGT (Ickleton 52.063481N, 0.170783E and Gasworks 52.081978N, -0.169570E), Elsoms Wheat Ltd (Weston, 52.842426N, -0.079539E), KWS (Fowlmere, 52.0533N, 0.03551E). Details on seasons and populations in Supplementary Table 23.

### NIL development

*Construction of the NIL library.* QTLs for putative advantageous alleles from landrace parents were targeted for NIL development. For each QTL, a RIL was selected that carried the landrace allele at all markers of the QTL confidence interval (CI) and that carried a maximum number of Paragon alleles at the remaining loci. Selected RILs were crossed with cultivar ‘Paragon’, followed by two backcross steps also to ‘Paragon’, ensuring the heterozygous state of the CI region in the selected parent for the next crossing step by marker assisted selection (MAS). All crosses were conducted with Paragon as pollen donor and plants were grown under standard glasshouse conditions. For the final step of the NIL development, BC2F1 lines were self-pollinated and BC2F2 lines homozygous for the CI region for both, either the landrace parent or Paragon, were selected by MAS to be used as NIL pairs or families for the QTL validation (an overview over the crossing scheme is given in Supplementary Fig. 31). A more complete genotype of the BC2F2 lines was determined using the 35k Axiom Wheat Breeders’ array^68^. NIL development progressed in annual sets, called Toolkit (TK) sets, starting in 2012. We report here on sets developed up to 2017 (TK1 to TK5, details of selected QTL, number of NIL families and individual NILs are given in Supplementary Table 48). BC2F2 NIL seeds are available from https://www.seedstor.ac.uk/search-browseaccessions.php?idCollection=40.

### Field Experiments for NIL families

The performance of NIL pairs, or families of NILs coming from one cross with opposing parental alleles in the targeted QTL region, were compared (Supplementary Fig. 31). After initial multiplication trials at JI (1 m^2^ plots) further field trials were conducted as yield trials (replicated 6 m^2^ plots in randomised block design) at JI, RH and SB under the same conditions as the RIL trials. The NIL families were grown in yearly TK sets between 2015 and 2022 (see details in Supplementary Table 23). Phenotypes recorded were similar to the RIL trials, and raw data can be downloaded from https://grassroots.tools/fieldtrial (Search term: “DFW Academic Toolkit Trial”). NIL family performance for measured phenotypes were assessed using simple ANOVA in individual years (Supplementary Table 41). The performance over all seasons and environments was assessed using AMMI (Supplementary Table 41). Specific NIL field experiments to assess the grain yield potential were conducted for NIL lines WL0019 and WL0026 at 15 different locations. Of those trials, six were the general NIL trials reported above; another six trials were part of grain yield germplasm trials by commercial UK breeders, grown in triplicated randomised 6 m^2^ plots, harvested in 2017 (details and raw data can be downloaded from https://grassroots.tools/fieldtrial, search term: DFW-BTK-H2017) and three similar trials at JI with harvest in 2018, 2019 and 2020.

In total, six trials for *RHT8* NILs were conducted with a NIL carrying *RHT8* and the recurrent NIL parent Paragon, with two seasons at each of the three locations: JI, the University de Lleida, Spain (41.36464N 0.4819.7E) and IFVC, Novi Sad, Serbia (45.20N, 19.51E). All trials were grown in a randomised block design together with other varieties with three replicates (JI) or five replicates (the other sites). All trials were autumn drilled and harvested in July at JI (2020, 2021) and in June at the other sites (2021, 2022). Plot size was 6 m^2^ at JI and 5 m^2^ at the other site. The average temperature was recorded at all three sites.

### Quantification of Trait Relationships

The trait-trait relationship analysis was performed using the phenotypic dataset of the TK trials, collected at multiple locations in multiple years. For individual years and locations, the Pearson correlation coefficient (r) was calculated using R package corr. Then, the median and mean of the r values over several trials was used as the final result of correlation coefficient. These results were visualised using R package corrplot.

Trait trade-off plots were created for all traits together as PCA plots and in individual plots and for the trait pairs: grain weight (GW) x grain number (GN); grain yield (GY) x grain protein (GP); and GY x heading date (Hd). The individual plots are based on the effect direction and effect size of the introgressed alleles from the 127 NIL families, where the majority of introgressed alleles come from Watkins and are compared to the Paragon allele. The trait effects were calculated using the AMMI method as described. In the trade-off plots, the traits effects are shown as filled circles on a horizontal bar, which represents a sliding scale of the allele effect as percentage of the mean trait value. A positive allele effect will result in a positive value. For the trade-off plots, two traits are shown together, with the bar for the first trait (blue) being on top and for the second trait (pink) at the bottom. The circles are shown in colour (blue, respectively pink) if the effect was statistically significant, and in grey otherwise (Fig. 2f). The PCA bi-plots on the left of the individual trade-off plots are based on a PCA calculated using the phenotype estimators from the AMMI, employing the package princomp in R. Specific trait pairs are highlighted by bold arrows in the PCA plots to show their overall relationship. This contrasts the individual trade-off plots, which show exception to this overall trend.

### Genome-wide Association Study from Watkins collection

The markers used for GWAS of Watkins collection were ∼10 Mb core SNPs in dataset 3. Extreme outlier values of phenotypic data were removed. Based on these, we performed GWAS using GEMMA (v0.98.1)^74^ with parameters (gemma-0.98.1-linux-static -miss 0.9 –gk -o kinship.txt) and gemma-0.98.1-linux-static -miss 0.9 -lmm -k kinship.txt). In-house scripts programmed in R were used to visualise these results.

### NAM imputation and NAM GWAS

*Pre-processing for skeleton marker*: The accessions of NAM/RIL populations were genotypes by 35 kb SNP array^58^. To obtain a high-quality SNPs dataset, we used marker flanking sequences to align to the reference genome (IWGSC V1.0)^11^. Positions and alleles of SNPs that were consistent with the resequencing data were retained and then only markers with polymorphisms between parents were retained.

*NAM Imputation*: The HapMap constructed in this study was the base for NAM imputation. The SNPs of RIL parent were extracted from SNP dataset 1, since the rare alleles in the natural population will not be rare in the RILs. Overall, we used the 35 kb array results of RIL offspring as the skeleton to predict the genotype of each site in each accession. For each RIL population, the detailed methods are as follows:

Step 1:

1. Traverse the SNP sites of each parent.
2. The SNP locus is in a haploblock: If there is one or more skeleton markers in the block, the offspring genotypes will be filled according to the nearest skeleton marker in the block; If there is no skeleton marker in the block, the progeny genotypes are filled according to the nearest skeleton marker on the chromosome.
3. The SNP locus is not in haploblock: the offspring genotype is filled according to the nearest skeleton marker on the chromosome.
4. The method of filling offspring genotypes according to skeleton marker: for example, marker typing exists A or B or H or -, where A represents from parent 1, B represents from parent 2, H represents heterozygosity, and - represents missing. Then a SNP site can select which parent is in the SNP matrix genotype according to the marker typing.

Step2: Go through each RIL and do the same as step 1 for each RIL.

Step3: Use the bcftools merge command to merge all RIL groups to generate vcf files.

Step4: Two steps of LD pruning were carried out (plink --indep-pairwise 10kb 1 0.8; plink --indep-pairwise 50 1 0.8)

*Percentage of the environmental mean*: To standardise phenotypic values across different environments, we calculated the ’percentage of the environmental mean’ for each trait. For each individual trait, the raw mean values were first used to calculate the environmental mean of that trait. Next, the individual trait values were converted into a percentage of this environmental mean. The formula used was: percentage of environmental mean = (individual trait value / environmental mean of the trait) *100. By doing so, each trait’s value was expressed as a percentage relative to the mean trait value in that environment. Importantly, before the calculation, controls and outliers were excluded from the data. This approach allowed us to compare traits on a similar scale, effectively reducing potential bias introduced by the raw numerical values of different traits.

*NAM GWAS*: Based on the SNP sets generated after NAM imputation, two-step LD filtering was performed: plink --indep-pairwise 10kb 1 0.8; plink --indep-pairwise 50 1 0.8. Finally, 19,253,188 SNPs were retained. We then collated the following data for each phenotype used to perform NAM GWAS:

1. Phenotypic value of a single year, in which the phenotypic of the offspring of the RIL population involved in this year is directly integrated.
2. Phenotypic values of years with high nitrogen content. the phenotypic values of progeny of all RIL populations in years with high nitrogen content were integrated, using percentage of the environmental mean value.
3. Phenotypic values of low nitrogen years: the phenotypic values of offspring of all RIL populations in all years involved in low nitrogen environments were integrated, using the percentage of the environmental mean value.
4. The phenotypes of all years and all environments are combined, using the percentage of the environmental mean value.

### Discovery and functional verification

*RHT8 fine mapping and gene discovery.* Seventy two recombinant lines within the *RHT8* locus (Supplementary Table 43) were used for genetic mapping^30^. These are from the cross: RIL4 (from the Cappelle Desprez x Cappelle Desprez (Mara 2D) population described^75^ with Cappelle Desprez. Seven new KASP markers were designed and used as described^76^ based on Watkins SNPs (Supplementary Table 44).

*Blast Pm4 gene discovery and Proof of Function.* Seedling leaves of a set of 300 diverse Watkins genotypes were screened for resistance to two isolates of *Magnaporthe oryzae* pathotype *triticum* (*MoT*) that carry the effector *AVR-Rmg8* using the methods described^77^. Spike resistance to these isolates was assessed using the methods described^77^. Validation of the candidate gene recognising *AVR-Rmg8* was undertaken using the loss of function mutants and overexpressing transformant lines described for *Pm4* (ref^37^).

*Calcium NAM GWAS and Proof of Function.* Wheat lines carrying induced mutations in either of two candidate genes for grain calcium content (GrnCaCnc), discovered in the NAM analysis (Fig. 4g), *TraesCS5A02G542600* and *TraesCS5A02G543300*, were identified in the Cadenza TILLING population^78^, following the method described in Adamski et al. 2020^79^. Only mutations that were predicted to lead to a gained stop codon, to a missense variant or to a splice donor variant were selected. Two independent mutations were selected for *TraesCS5A02G542600* and eight for *TraesCS5A02G543300*. For each mutation, 24 seeds of the TILLING lines were grown under standard glasshouse conditions. Ten wildtype Cadenza plants were grown as control. Plants were genotyped with KASP markers specific for the presence/absence of the mutations and only homozygous mutant plants were taken forward (in total 53 plants, between 4 to 7 individual plants for each tilling mutation, mean 5.3 plants per mutation). From each of these plants, all grains were harvested and grain number per plant (GNplant), grain yield per plant (GYplant) and the seed characteristics GW, GLng and GWid were measured using a Marvin seed analyser. Grain moisture content was measured using DA 7250 Near-infrared spectrometer. GrnCaCnc was measured using X-Supreme8000, a benchtop X-ray fluorescence spectrometer, equipped with XSP-Minerals’ Package and calibrated with data collected using an ICP-OES using 187 data points with GrnCaCnc levels ranging from 242.4 ppm to 726.7 ppm. No outliers for GrnCaCnc were detected and the average over the three technical reps was calculated. This data set (GrnCaCnc range 380.0-564.2 ppm, mean 461.3 ppm) was used to statistical compare the GrnCaCnc between the Cadenza wildtype and the independent mutants in a linear model (ANOVA). The data was also visualised in a violin plot (Fig. 4h). Of the mutant lines for gene *TraesCS5A02G543300*, all lines showed a higher level of GrnCaCnc in comparison to the Cadenza wildtype (1.7 % – 18.7% higher). Of these, five lines (WCAD1641, WCAD0289, WCAD1253, WCAD1003) showed statistically highly-significant increased levels of GrnCaCnc (14.1-18.7 % increase, *P* value < 0.005) and one line (WCAD1617) statistically significant higher levels of GrnCaCnc (11.7 % increase, *P* value < 0.03) (Supplementary Table 47). The two mutant lines for gene *TraesCS5A02G542600* did not show significantly higher levels of GrnCaCnc. Of the four lines with highly-significant increased GrnCaCnc, three showed no significant change in GW (Supplementary Table 47), demonstrating that the GrnCaCnc increase is not simply a result of a reduced grain weight.

*Genetic dissection for yellow rust.* Seedlings of the Watkins lines were screened with single isolates of *Puccinia striiformis* under cold glasshouse conditions in April 2018. Two single isolates (UK 16/342 and 19/501) belonging to the ‘Warrior’ race, were tested separately as described^80^. Field screening was conducted in 1 m^2^ untreated plots. Yellow rust scores were: 1, no infection observed; 2, stripe per tiller; 3, 2 stripes per leaf; 4, most tillers infected but some top leaves uninfected; 5, all leaves infected but leaves appear green overall; 6, leaves appear ½ infected and ½ green; 7, Leaves appear more infected than green; 8, Very little green tissue left; 9, leaves dead-no green tissue left. In Egypt, we evaluated a diverse set of 300 Watkins bread wheat landraces and 20 additional control lines for yellow rust disease resistance and agronomic traits under natural field conditions at the Sids, Sakha, Nubaria and Gemmeiza Agricultural Research Center stations (Egypt) during the 2019/20, 2020/21 and 2021/22 growing seasons.

### Development of haplotype-based diagnostic SNPs and design of KASP markers

For each haploblock, we selected SNPs being able to differentiate all haplotypes within this block, defined as tagSNPs. The detailed process is as follows.

Step 1: Distance calculation of all the SNPs between each pairwise haplogroups. The genotype encoding rules used during the distance calculation are as follows: Reference allele (homozygous): -1; Alternative allele (homozygous): 1 ; Missing: 0; Heterogeneous: 0.9.

For each SNP site, we first calculated the state of this SNP within the haplogroups. Then the average genotype state for all accessions within its respective haplotype cluster was computed. Let’s denote the states of the SNP in haplotype clusters 1 and 2 as *s*_*h*1_ and *s*_*h*2_, respectively:

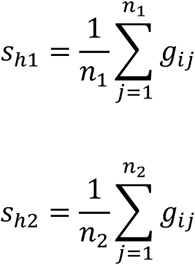

Here, *g_ij_* is the genotype of the *j*-th sample in haplotype cluster *i*, and **n**_1_ and *n*_2_ are the counts of samples in haplotype clusters 1 and 2, respectively.

Step 2: Calculation of the distance of this SNP position between haplotypes. The Euclidean distance between the average genotype states of two haplotype clusters is then calculated:

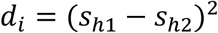

If the SNP falls within a coding region, its weight is quadrupled:

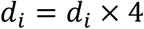

Step 3: Sort of distances of all the SNPs. The SNPs are sorted based on distances, and the SNP with the maximum distance is chosen as the tagSNP. Within a haploblock, a tagSNP is selected for each pairwise haplogroups. The union of these selected tagSNPs forms the tagSNP set for the haploblock.

Step 4: This process is repeated across all 71,000 haploblocks in the genome to compile the complete set of tagSNPs.

In summary, for each haploblock, the haplogroups that each accession belongs to were determined in the above HapMap analysis. We further calculated the distances at all SNPs between each pairwise haplogroups. The SNP with the maximum distance were chosen as the tagSNPs that distinguishes these two haplotypes.

### Development of a *Triticum aestivum* Next Generation Genotyping Array (*Ta*NG)

Single nucleoptide polymorphism calls for 208 elite lines and 111 landraces from the Watkins “core collection” were selected from the entire skim sequence dataset (Supplementary Table 52), having excluded varieties with >= 1% heterozygous loci. Selection of SNPs with an even distribution across the genome was performed as follows (https://github.com/pr0kary0te/GenomeWideSNP-development). SNPs were initially filtered to have a maximum of 0.5% heterozygous calls among the 315 selected varieties, a minimum MAF of 0.01, a minimum call rate of 0.95 and a minimum mapping quality score of 5000. Two steps were then taken to filter out non-unique SNP loci. First, SNPs were removed if their flanking sequence mapped to more than one genome location in the IWGCSv1.0 Chinese Spring genome using the BWA version 0.7.12-r1039 ^51^. SNPs passing this filter were then checked by BLAST (blastn v2.6.0+)^81^ against the the IWGCSv1.0 Chinese Spring genome to exclude those that matched multiple locations. Each chromosome was then divided into 1.5 Mb intervals and a set of SNPs selected from candidates in this interval which had the highest combined varietal discriminatory power. Up to six of the highest ranked SNPs were assigned to each bin. This approach has been described previously for the selection of a minimal set of SNP markers capable of discriminating apple varieties^82^, where the SNP with the highest varietal discrimination is chosen initially, and further SNPs are selected to add the maximum additional discrimination until either all varieties are discriminated, the addition of more SNPs does not improve discrimination, or all available SNPs have been evaluated. SNPs designed with this approach were supplemented with 1974 DaRTseq markers (supplied by Susanne Dreisigacker, CIMMYT) and 6388 markers from the existing Wheat Breeder’s Axiom array ^58^ for cross-compatibility of results and to infill some spaces that became available on the array late in the design process.

In total, 44,266 SNP markers were placed on the high-density Axiom genotyping array by Thermo Fisher design team. This was designated the *Ta*NG (*Triticum aestivum* Next Generation) genotyping array. Initial screening of the array with a diverse set of elite and landrace lines showed that a high proportion of the SNPs tiled on the array (over twelve thousand in total) failed to convert to polymorphic SNP assays. These were replaced with 13,065 SNPs taken from our existing 820K wheat genotyping array ^13^ by comparing genotyping data on a common set of varieties genotyped on both the 820K and TaNG design and selecting those that maximised the differentiation of varieties in 1.5 Mb bins as described above. This second-round empirically optimised design was designated TaNG1.1 (Axiom 384-well array Thermo Fisher catalogue number 551498), comprising 43,373 markers.

## Extended Data Figures 1–10

**Extended Data Fig. 1.**
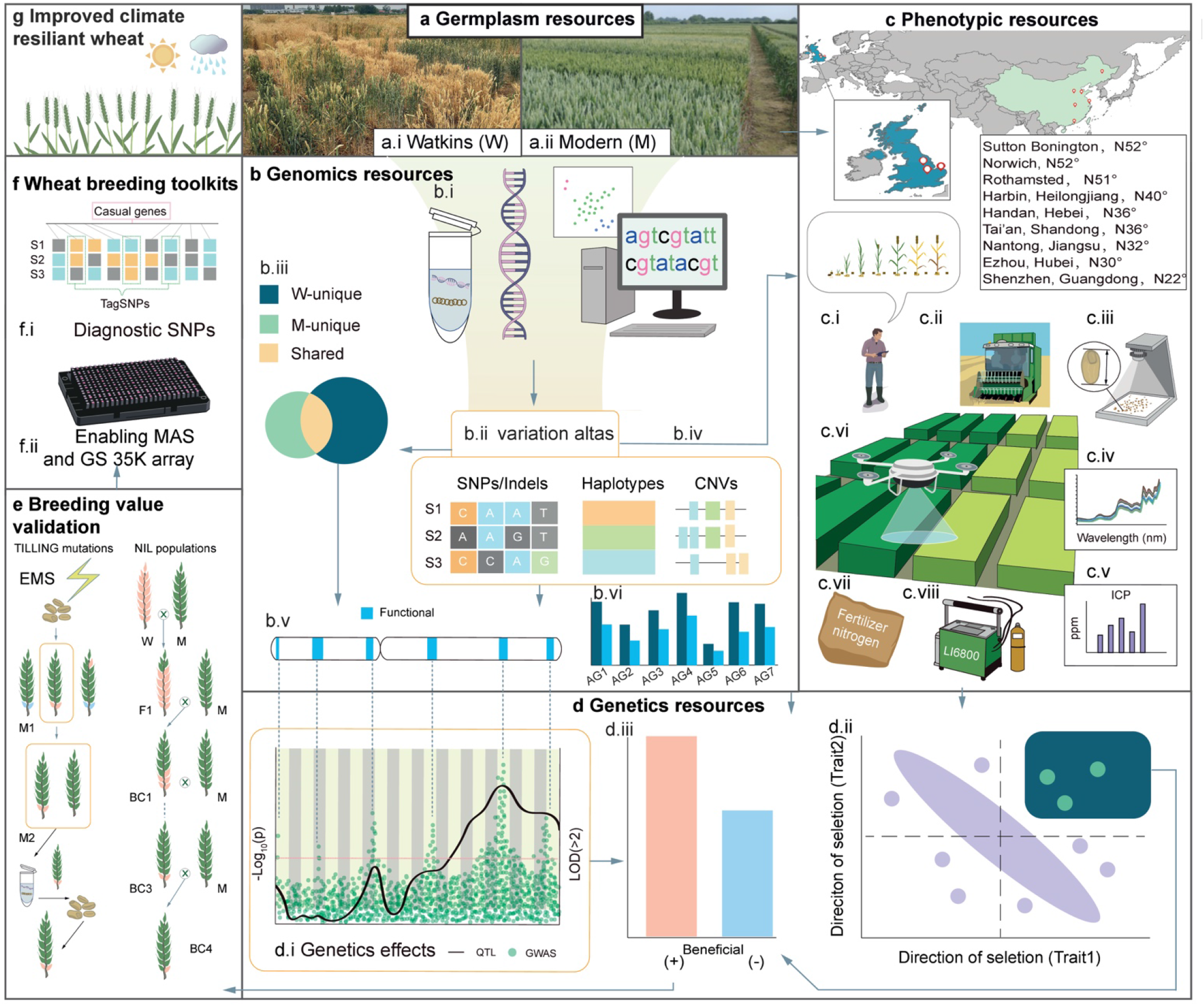
Graphical abstract and conceptual strategies. **a,** The WWWG2B strategy started with the comparison of variations between the Watkins landrace collection (W, a.i) and the modern wheats (M, a.ii). **b,** By developing an extensive genomics resource wedetermined the extent to which landraces carry variants that are not present in modern wheats. **c-d,** Natural and structured populations were then combined in multi-site field experiments to identify novel, functional and useful genetic variations not yet deployed in modern wheat. This required unprecedented genetic dissection by a combination of whole-genome resequencing (b.i), construction of variant atlas and haplotype map (b.ii) and high-throughput field-based phenotyping (c.i-c.viii) of next-generation-gene-discovery populations, the NAM RIL segregating populations combined with GWAS (d.i), bi-parental QTL mapping, and haplotype analysis enabled by the development of the advanced genomics and genetics resources shown in b and d**. e,** Alleles with high breeding values and their phenotypic effects (determined by calculating the AMMI means for their selection) were validated and delivered for use in breeding. **f,** Diagnostic SNPs and KASP markers were designed for assisted molecular breeding. **g,** The overall objective of the Watkins Worldwide Wheat Genomics to Breeding (WWWG2B, http://wwwg2b.com/) consortium is to enable the development of a new generation of modern elite wheat cultivars that are climate resilient. The flow of useful genes and alleles through this process was gated by three prioritisation criteria: 1) the alleles or haplotypes are novel or unique to Watkins landraces or are at very low frequency in modern wheat (b.iii); 2) the Watkins-unique alleles or haplotypes are functional with significant genetic effects (target QTL and MTA) (d.i); 3) haplotype analysis showed that the Watkins-unique functional alleles or haplotypes are beneficial with increasing effects based on our understanding of physiological trait relationships (d.ii and d.iii). These prioritisation selection criteria are used to choose alleles to build new modern wheat cultivars while avoiding negative trade-offs. Features highlighted within each section are as follows: **a**, Germplasm resources. Comparison between the Watkins landrace collection (W) and the modern wheat cultivars (M). **a.i,** the A.E. Watkins landrace collection (*n* = 827), which is the source of plant genetic resource variation for the WWWG2B project. **a.ii,** selection of 208 modern elite wheat cultivars selected from the public 35K Axiom genotyping data hosted at CerealsDB (https://www.cerealsdb.uk.net/cerealgenomics/CerealsDB/indexNEW.php). **b**, Genomics resources. Exploitation of the genomics for target breeding values. **b.i,** Whole-genome re-sequencing and alignment of reads to the Chinese Spring reference genome (IWGSC v1.0) to identify single nucleotide polymorphisms (SNPs), copy number variations (CNVs) measured by read depth variants (RDVs) and haplotypes based on SNP linkage disequilibrium (LD), (**b.ii** and **b.iii**). **b.iv,** Whole-genome imputation of 73 RIL populations (6,762 RIL genotypes). **b.v,** Local details of the target genomic interval with dissection of local haploblocks and haplotypes of the allelic series, the chromosome distribution and frequency across sub-populations of haplotypes were estimated, specifically for those that are specific to modern wheat or Watkins wheat (**b.vi**). **c**, Field trial and phenotyping. **c.i**, Manual collection of field phenotypes such as height and phenology. **c.ii,** Plot combine harvester for measuring yield and biomass components. **c.iii,** Digital seed analysis (such as grain length, width, area and number). **c.iv,** Near-infrared reflectance (NIR) and CHN elemental analysis for nitrogen distribution. **c.v,** Inductively coupled plasma optical emission spectrometry (ICP-OES) and X-ray fluorescence (XRF) for mineral analysis. **c.vi,** Unmanned aerial vehicles (UAVs) capturing normalised differential vegetative index (NDVI). **c.vii,** Nitrogen fertiliser treatment and measurement of nitrogen component traits. **c.viii,** LI-6800 portable photosynthesis system for examining leaf physiological traits such as CO2 respiration and leaf chlorophyll content. **d,** Exploitation of the genetics resources for target breeding values. **d.i,** Genome scanning of genetic effects by approaches combined GWAS-, NAM GWAS- and biparental QTL analysis. **d.ii,** direction of selection (DOS) for a trait of interest. The green rectangles represent field plots of each RIL. **d.iii,** calculation of phenotypic effects (AMMI mean values) to quantify breeding value based on the genetic effects identified by QTL and GWAS, together with trait performance and trait relationships., Quantification of favourable or unfavourable haplotypes based on QTL/MTA analyses; the breeding direction towards beneficial alleles is determined based on breeding purposes. **e**, Breeding value validation. Prioritised WWWG2B alleles were tested in modern wheat background by producing Near Isogenic Lines (NILs) or TILLING mutants. **f**, Development of the breeding toolkits. **f.i**, Full exploitation of WWWG2B outputs required the development of new selection tools including the *Triticum aestivum* Next Generation high density genotyping array (*Ta*NG) and KASP assays based on haplotype-specific tagSNPs. **f.ii,** The development of the diagnostic SNPs; experiments for the delivery of WWWG2B outputs were designed in cooperation with wheat breeders using precompetitive platforms. **g**, Vision and mission: advanced outputs from WWWG2B represented in modern wheat cultivars.

**Extended Data Fig. 2.**
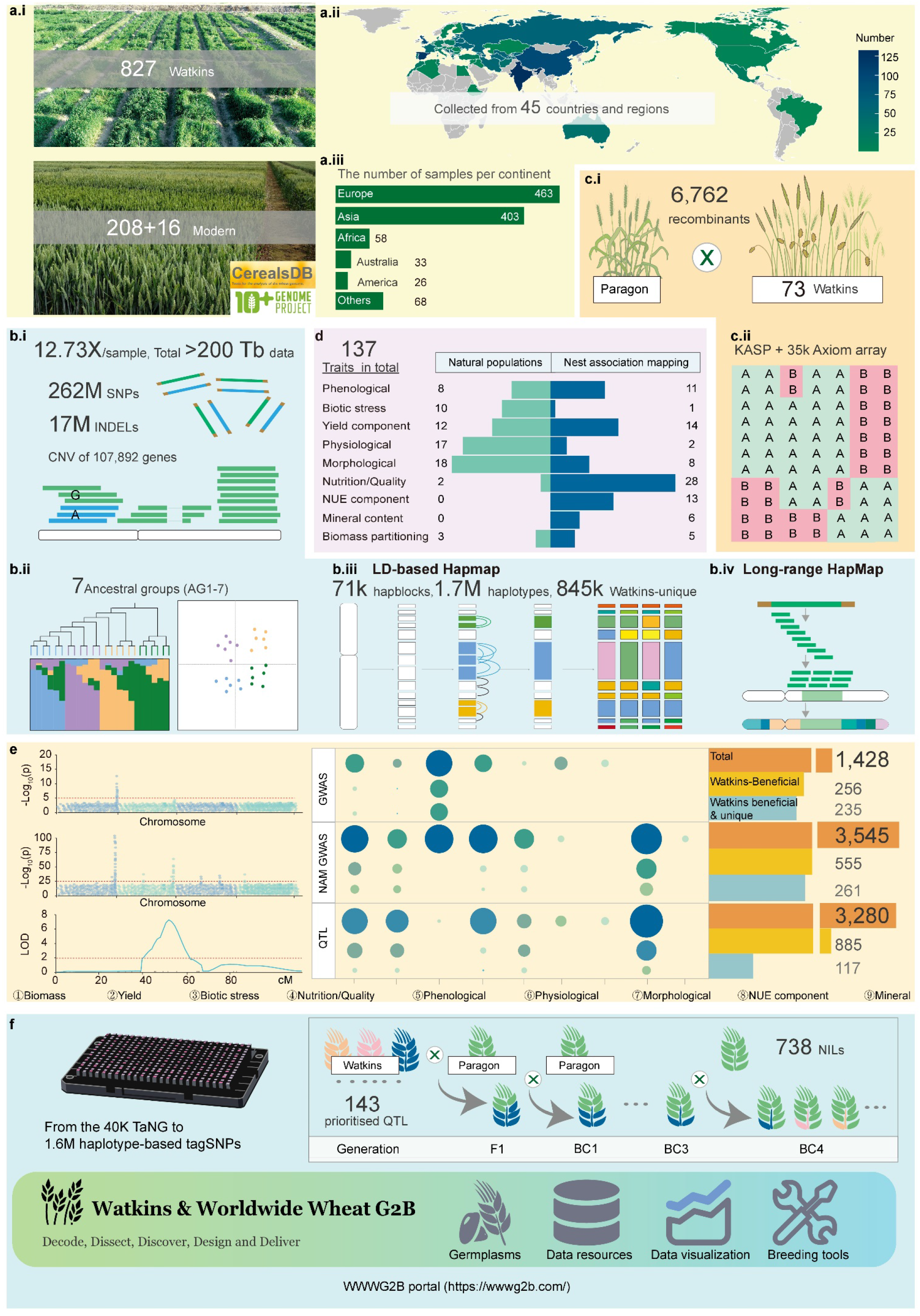
WWWG2B roadmap visualised by numbers to summarise the datasets in this study. **a,** Germplasms used in this study, including the entire A.E. Watkins Landrace Collection (827 accessions), 208 Modern elite wheat cultivars and the 16 modern accessions used in the 10+pan-genome project including Chinese Spring (in total 1051 accessions, **a,i**) originated from 45 countries and regions (**a.ii, a.iii**). **b,** Development of the genomics resources, including whole-genome re-sequencing, with 12.73X genome coverage per accession, resulting in >200 Tb raw data in total, including ∼ 262 million high-quality SNPs (after inbreeding co-efficiency quality control), 17 million short insertions and deletions (<50 bp), and quantification of copy number variations for each protein-coding gene (**b.i**); population structure analysis: seven ancestral groups (AGs) of the 827 Watkins landraces, together with modern wheat, were used to construct a framework for comparison across the 1051 wheat accessions (**b.ii**); SNPs-based haplotype map based on linkage disequilibrium (LD was calculated by Plink, **b.iii**), and a *k*-mer IBS based long-range (>1 Mb window) haploblock map (**b.iv**). **c**, Development of the genetics resoures, including a 73 RIL populations with parents selected to represent the maximised diversity from the seven ancestral groups, forming a structured NAM population (**c.i**); In the F4:5 NAM populations, genotyping of the populations/recombinants was performed using both KASP markers and the 35K Axiom array (**c.ii**). **d**, Development of phenotyping resources, including 70 traits surveyed from the Watkins collection diversity panel grown in both the UK and China over the past 17 years and 88 traits from nine categories in the NAM populations (F4:5). In total, 137 traits were surveyed in this project. **e,** the development of the academic toolkits. Thousands of significant QTL and MTA were identified and catalogued for the nine trait categories, by approaches of whole genome-wide association study (GWAS), bi-parental QTL mapping study (QTL mapping), and NAM GWAS after NAM imputation, the number statistic of the identified genetic effects for each approach was given. **f**, Development of the breeding toolkits. 143 prioritised QTL segments were selected and introgressed into Paragon by marker-assisted backcrossing to test their genetic effects and measure the phenotypic effects of each QTL segment by growing the introgressed lines in different fields. The new 40K *Ta*NG array was designed and tested based on the high-quality SNP datasets; diagnostic SNPs were selected based on the haplotype map; a WWWG2B database and pre-breeding portal were built.

**Extended Data Fig. 3.**
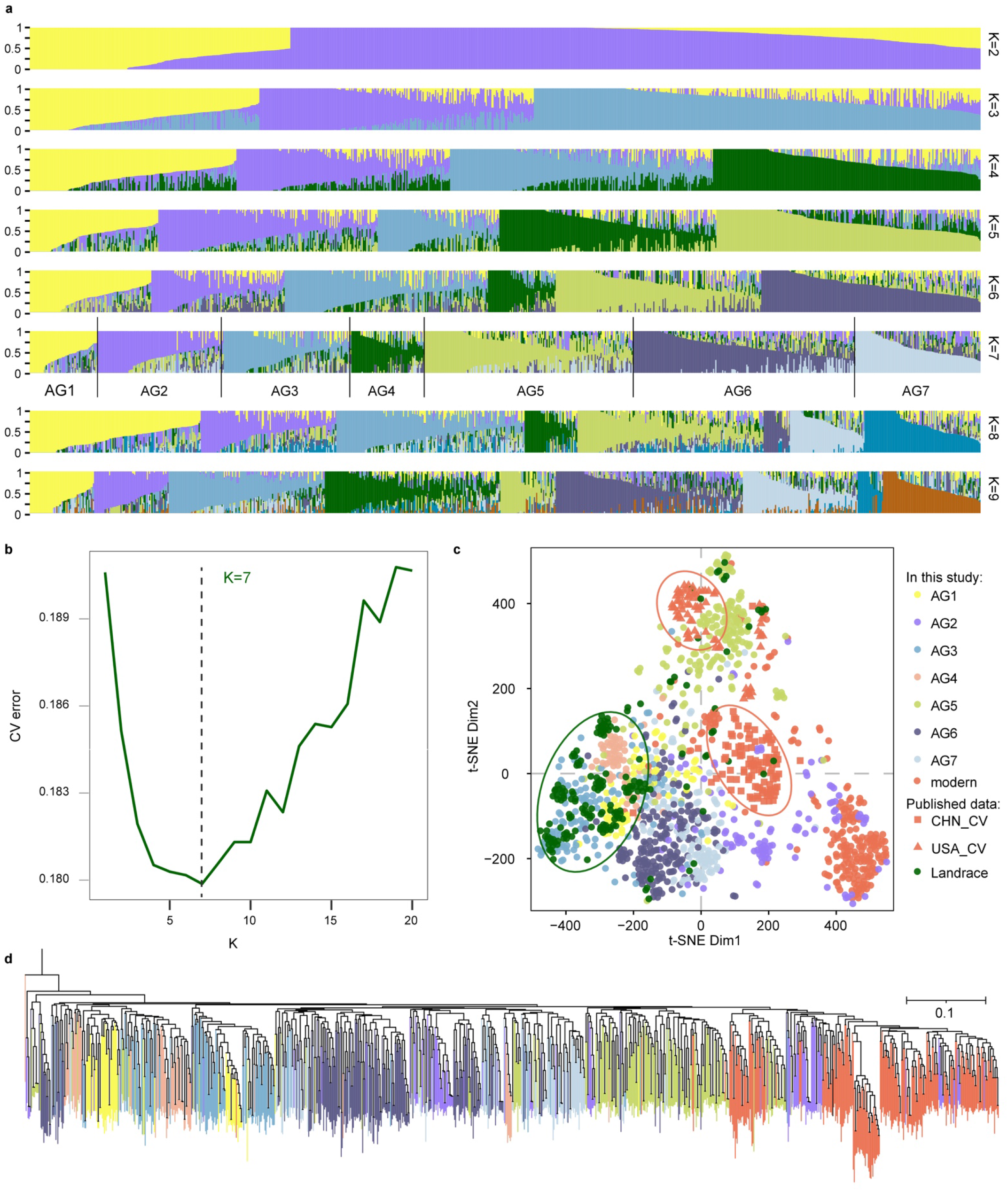
Population structure and phylogenetic analysis of Watkins landraces and modern wheat cultivars. **a,** Genome-wide ADMIXTURE results for 827 Watkins landraces from K=2 to K=9. Each colour represents an ancestral population. The length of each segment in each vertical bar represents the proportion contributed by ancestral populations. **b,** Estimated CV error for different K values (from K=2 to K=20) in the ADMIXTURE analysis with a K of 7 designating the ancestral groups (AGs) identified here. **c,** dim1 and dim2 plots of t-SNE results using Plink haplotypes, for the merged variation matrix (4 M shared SNPs) between the SNP matrix built in this study (the 10M core SNPs, see Methods) and the published dataset (76 M SNPs) from Niu et al.^14^. **d,** Phylogenetic tree of a set of 1,687,965 tagSNPs using rapidNJ with 1000 bootstrap replicates. The seven AGs are indicated by different colours (as in **c**).

**Extended Data Figure 4.**
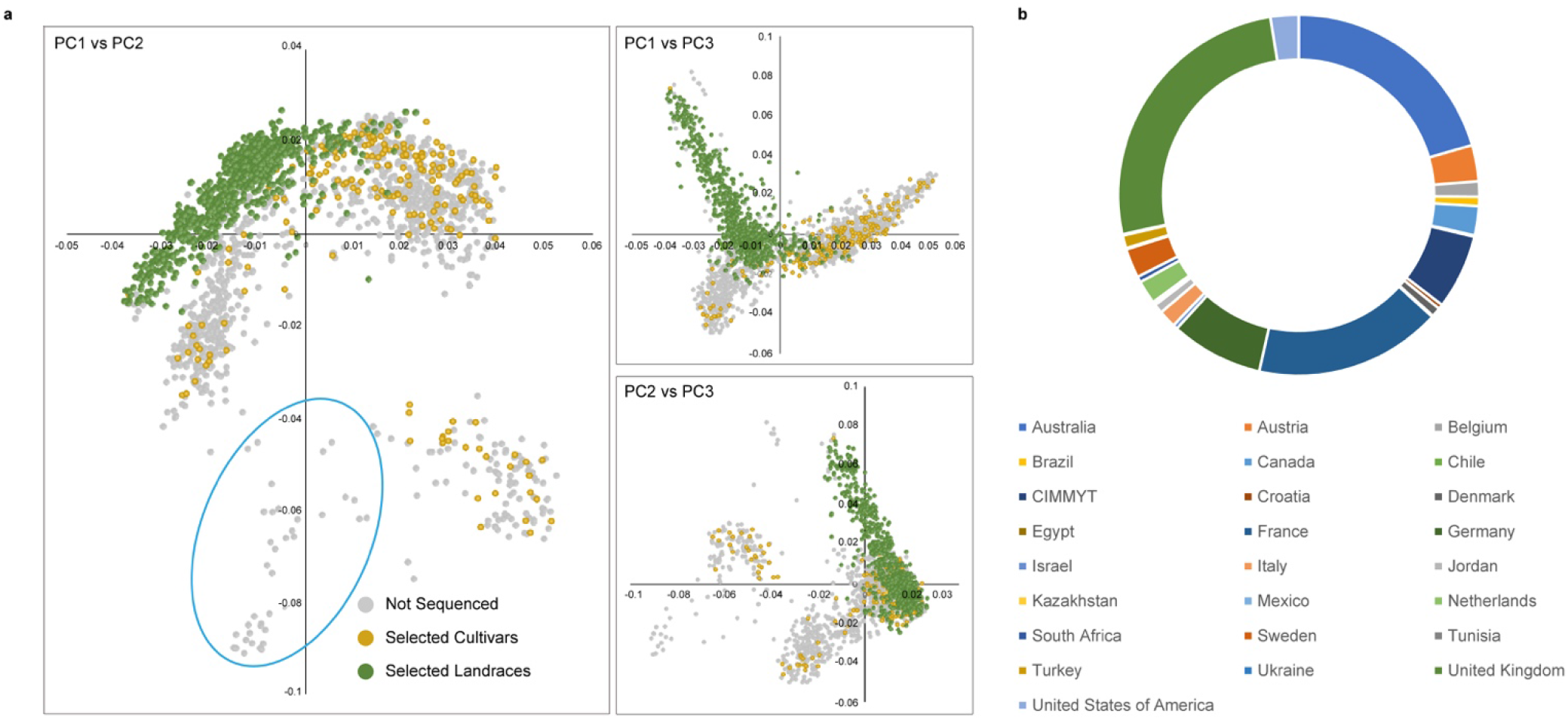
Choice of modern wheat cultivars and comparison with the Watkins collection. **a,** Principal component analysis based on Axiom 35K genotypic data for 3500 cultivars hosted by CerealsDB (https://www.cerealsdb.uk.net/cerealgenomics/CerealsDB/indexNEW.php) when selections for sequencing in this project were made. Selected cultivars are shown in gold, Watkins in green and cultivars not selected for sequencing in grey. Representative cultivars from the blue circled group in PC1 vs. PC2 were not selected because synthetic wheat derivatives are highly represented in this group. **b,** Countries of origin of the 3500 cultivars genotyped.

**Extended Data Fig. 5.**
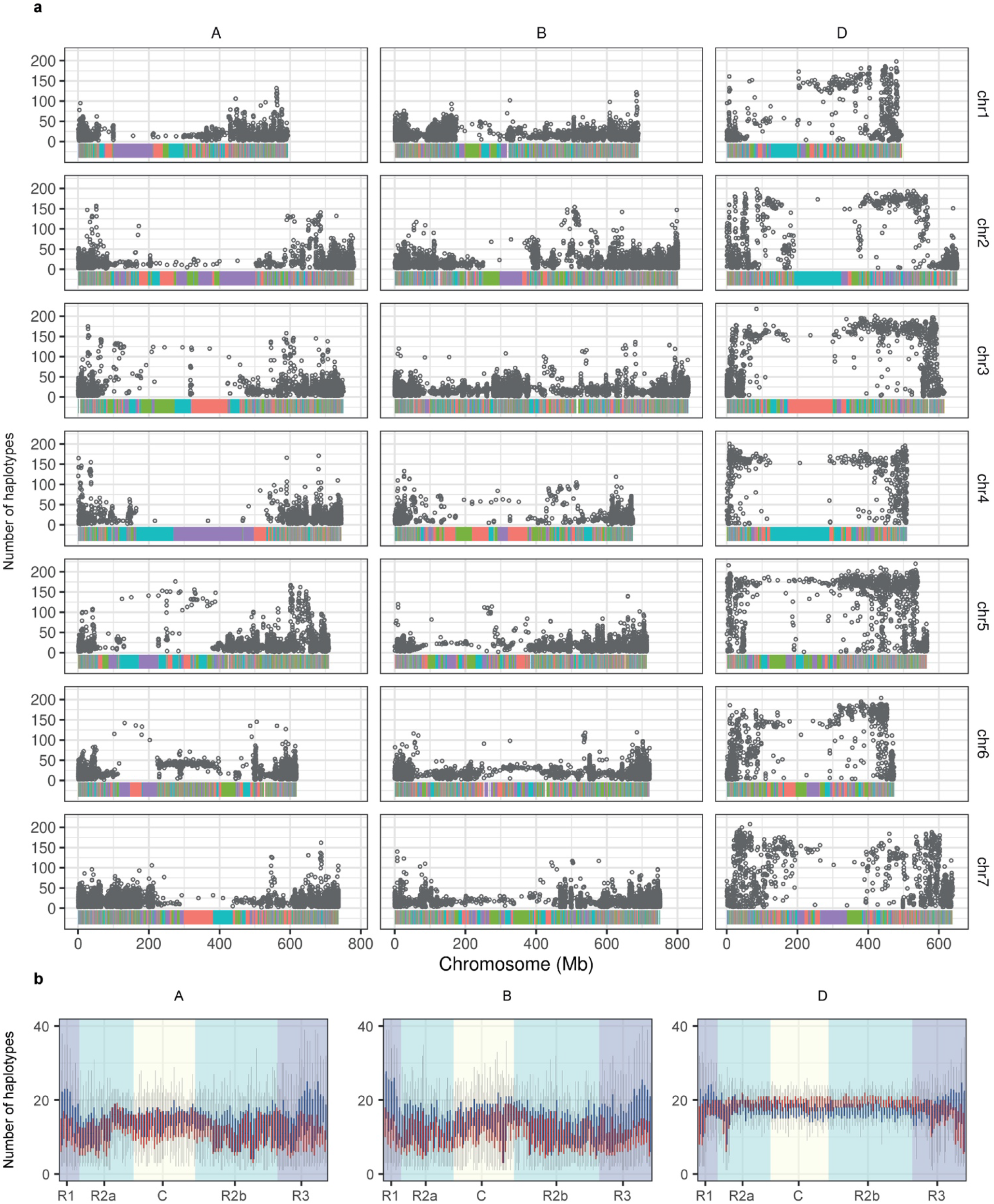
Wheat LD-based haplotype map. **a,** The multicoloured horizontal bar at the bottom along each chromosome shows the genome structural distribution of the haploblocks and a comparison of the A, B and D subgenomes; colours are used to distinguish adjacent haploblocks. The grey circles show the distribution of the number of haplotypes for each haploblock. **b,** Distribution of the number of haplotypes in 1Mb windows in the A, B and D subgenomes, where the background colour represents different chromosomal regions. Each chromosome were separated into 100 bins averagely, resulting in several 1MB windows per bin. One box represents one bin (window), red box for Modern, and blue box for Watkins.

**Extended Data Fig. 6.**
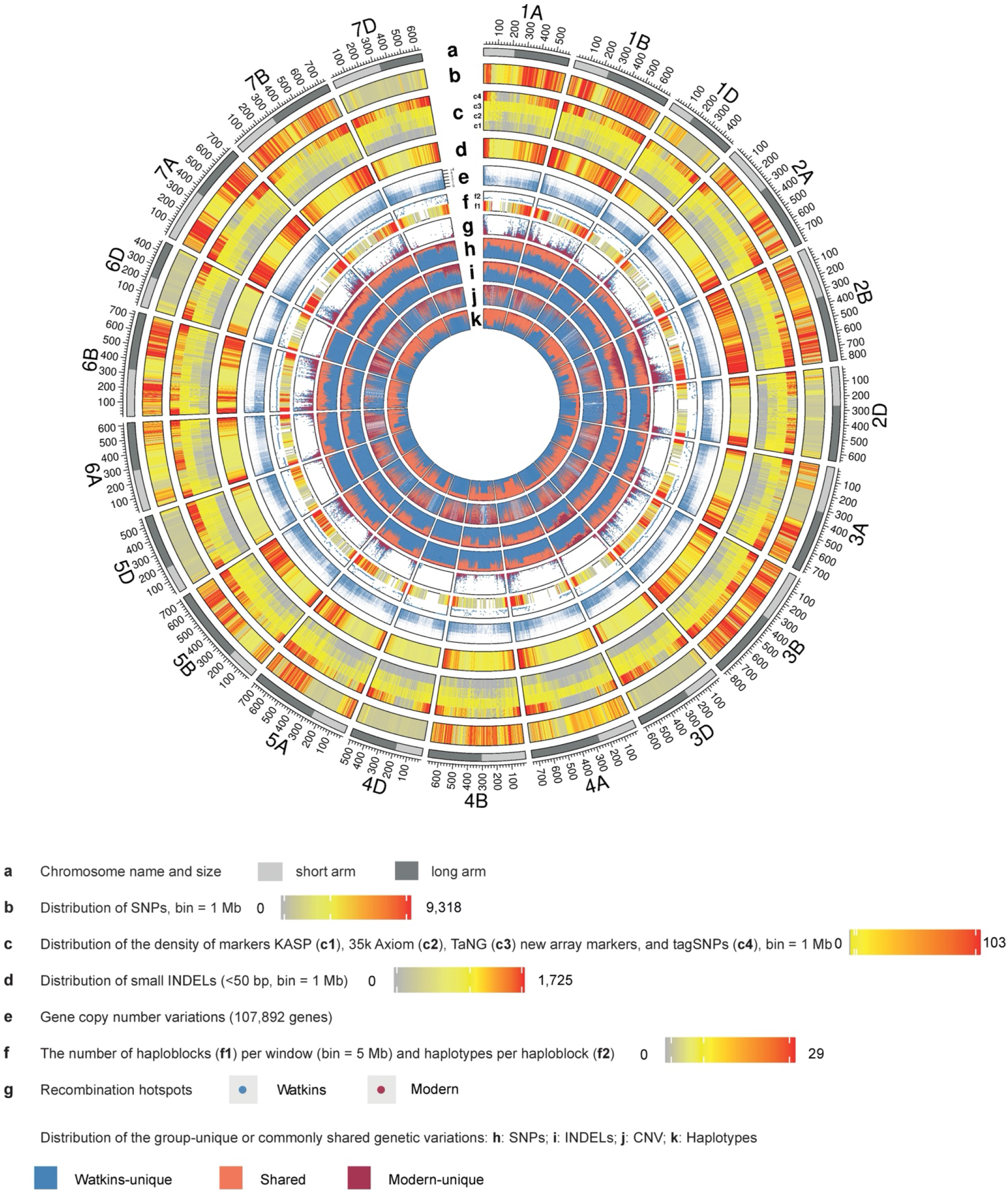
Genomics resources and the genetic diversity landscape. **a,** Chromosome names and sizes with short/long arms highlighted (bin = 100 Mb; light grey: short arm; dark grey: long arm). **b,** Distribution of high-quality SNPs identified in this study. **c,** Distribution of the density of genetic markers; circles (from the inner to the outer layer) represent the following: KASP markers (c1), 35k Axiom markers (c2), *Ta*NG markers (c3) and tagSNPs (c4; Bin = 1 Mb), each of which has a KASP marker designed for marker-assisted haplotype selection. **d**, Distribution of short INDELs (<50 bp) identified in this study. **e,** Gene copy number variation for each of the wheat protein-coding genes. **f,** Wheat LD-based HapMap: the heatmap shows the density distribution of haplotype blocks along the chromosome (f1, bin = 5 Mb), and the points (f2) represent the median haplotype number on the chromosome (5 Mb window). **g,** Distribution of recombination hotspots on chromosomes. **h,** Distribution of SNPs that are unique to either Watkins landraces or modern wheat cultivars or are shared. **i,** Distribution of small INDELs (<50 bp) that are unique to either Watkins landraces or modern wheat cultivars or are shared. **j,** Distribution of gene copy number variations that are unique to either Watkins landraces or modern wheat cultivars or are shared. **k,** Distribution of haplotypes that are unique to either Watkins landraces or modern wheat cultivars or are shared.

**Extended Data Fig. 7.**
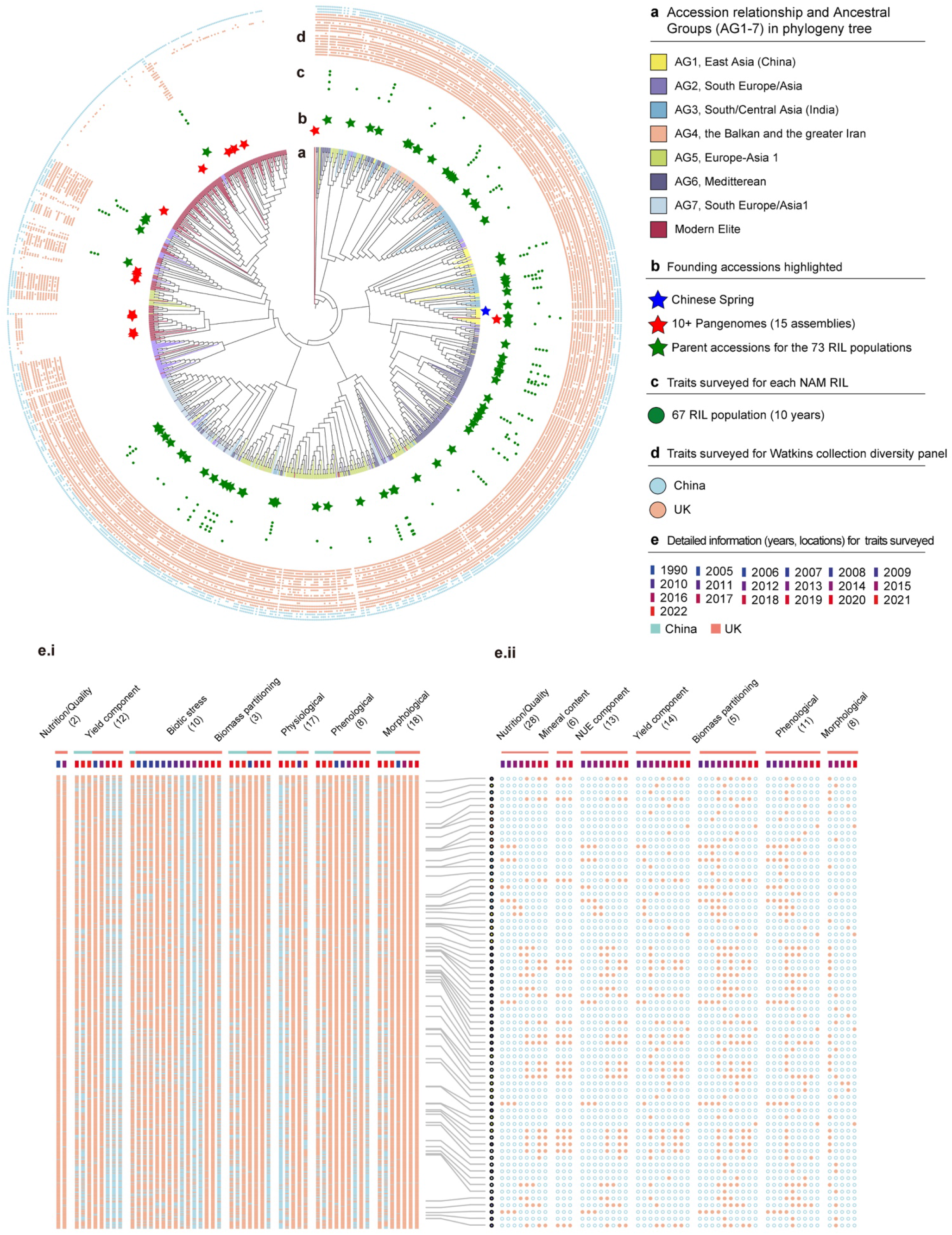
Watkins collection and mapping populations, phenotypic resources and traits surveyed summarised in a phylogeny-based circular diagram. **a,** Phylogenetic tree of the wheat accessions examined in this study. The phylogenetic tree was constructed using a set of 133,222 four-fold degenerate sites using rapidNJ with 1000 bootstrap replicates. The seven ancestral groups (AG1–7) and modern wheats are colour-coded as in Fig. 1a. **b,** The founder parents (green stars) of 73 Watkins x Paragon RIL populations are marked, the 15 pan-genome lines (red stars) and Chinese Spring (blue star) are indicated on the phylogenetic tree. **c,** Traits surveyed in multiple environments and multiple years for each of the NAM RIL populations, in which the corresponding Watkins line was used as the non-common parent. Each track represents a year (total of 10 years: 2011, 2012, 2013, 2014, 2015, 2016, 2017, 2018, 2019 and 2020). **d,** Traits surveyed in multiple environments and multiple years for the Watkins collection diversity panel (natural populations) grown in five geographic locations across China (blue circle, four years: 2020, 2021, 2022, 2023) and the UK (orange circle, 16 years: 1990, 2005, 2006, 2007, 2008, 2009, 2010, 2011, 2012, 2013, 2014, 2015, 2018, 2019, 2020 and 2021). **e**, Magnified view of data from the detailed field experiments, traits and phenotyping datasets as indicated in track d, including the traits surveyed in the Watkins collection diversity panel in multiple environments (e.i) and in multiple years (left) and in the RIL populations (NAM RILs) in multiple environments and in multiple years (e.ii).

**Extended Data Fig. 8.**
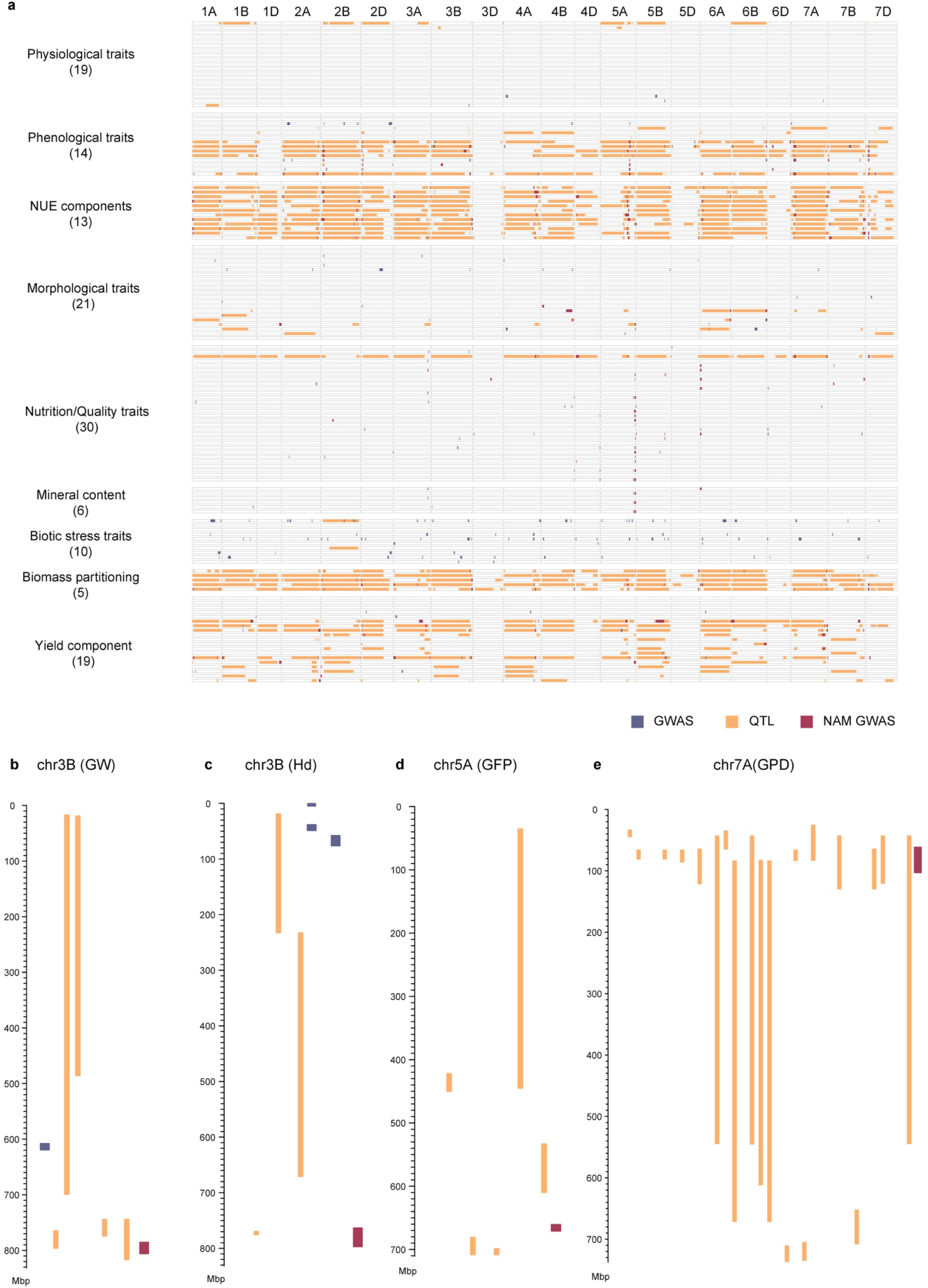
Summary of all the genetic effects identified, including QTLs detected by biparental QTL mapping, marker–trait associations (MTAs) from GWAS of the Watkins collection and MTAs from the NAM RIL populations (NAM GWAS). **a,** Whole-genome distribution of all the genomic intervals indicative of the identified QTL (orange) and the prioritised MTAs either from GWAS or NAM GWAS in this study. **b,** Magnified view of the examples with genetic effects detected for grain weight (GW) on chromosome 3B (Chr3B); **c**, heading date (Hd) on chromosome 3B; **d**, grain filling period (GFP) on chromosome 5A (Chr5A); **e,** and grain protein deviation (GPD) on chromosome 7A (Chr7A).

**Extended Data Fig. 9.**
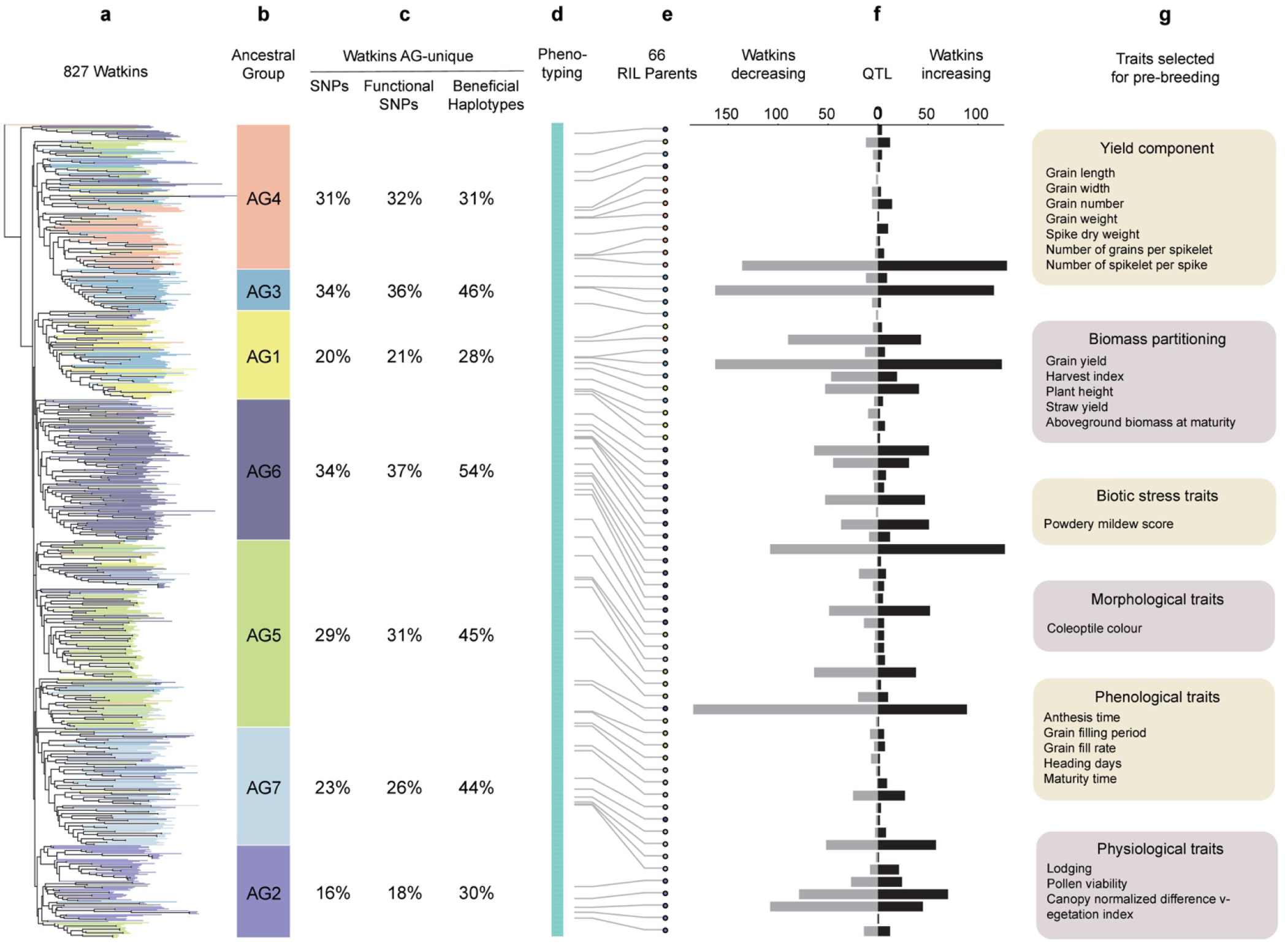
Information flow from novel and functional genetic diversity derived from Watkins landraces to the quantification of the beneficial increasing QTL allele associated with target traits. **a,** Phylogenetic tree of the 827 Watkins accessions, which were roughly clustered into the seven ancestral groups; these groups are roughly colour-coded in panel **b**. **c,** Percentage of AG-unique genetic diversity for SNPs, functional SNPs and beneficial haplotypes. **d**, Phenotyping of the Watkins collection including data collected in China and the UK and data for the 66 RIL populations in panel **e**, corresponding to Extended Data Figure 7. **f**, distribution of the number statistic of QTLs detected from the biparental QTL mapping populations, comparison was made for the beneficial QTL with increasing effects (right) and decreasing (left) in Watkins. **g**, Prioritised QTL and the major traits selected for introgression into Paragon via backcrossing to test their phenotypic effects for pre-breeding.

**Extended Data Fig. 10.**
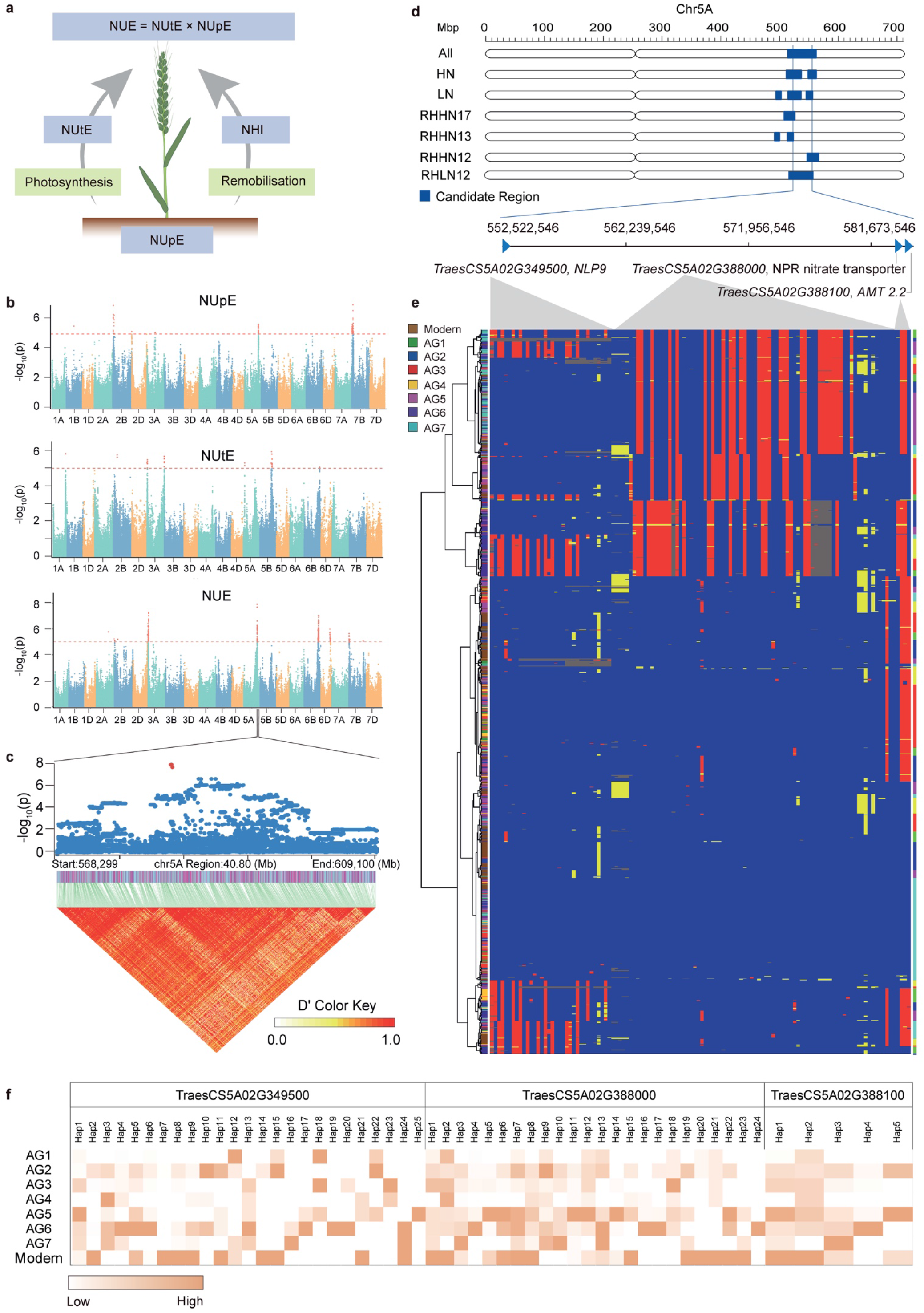
Trait dissection and genes discovery for nitrogen component traits. **a,** Schematic illustration of nitrogen component traits and their relationships: Nitrogen Use Efficiency (NUE), Nitrogen Utilisation Efficiency (NUtE) and Nitrogen uptake Efficiency (NUpE). **b,** Manhattan plots for NUE, NUtE and NUpE from a phenotyping dataset obtained in 2012. **c,** Local Manhattan plot with visualization of the LD haploblocks of the candidate genomic interval of the target peak located on chromosome 5A (Chr5A). **d,** An example to show the overlapped genomic interval of the genetic effects (QTL analysis from bi-parental mapping populations, MTA from NAM GWAS) detected on Chr5A, for NUE; three nitrogen-related transporters are highlighted. **e,** Visualisation of allelic diversity and haplotype clustering for the “nitrogen transporter region” with the three key genes indicated in the main text. **f**, Distribution and frequency of the different haplotype clusters for the “nitrogen transporter region”. Similar analyses were performed for other traits (see WWWG2B website, the genetic resources: https://wwwg2b.com/).

