## Supplementary Notes and Figures for "Harnessing Landrace Diversity Empowers Wheat Breeding for Climate Resilience"

### Supplementary Note 1: Watkins Bread Wheat Germplasm Collection

#### 1.1 Historic overview of the A.E. Watkins landrace collection

The A.E. Watkins landrace collection was assembled by Arthur Ernest Watkins during the 1920s and 1930s and originated mostly from local markets across 32 countries. A.E. Watkins accomplished this by enlisting the assistance of the London authorities to deploy their extensive network of consuls and agents. These agents visited local farms and grain markets on his behalf, gathering grain samples to be sent to Cambridge. Additionally, part of the collection came from contributions by colleagues including N. Vavilov in St. Petersburg, O. Frankel in New Zealand, and T.H. Shen in Shanghai, the later who also introduced 1700 wheat accessions into China in 1932 through John Percival (Percival collection)[1]. The collection, at its height in the 1930s numbered several thousand accessions, including diploid, tetraploid and hexaploid forms of *Triticum*. At present, 827 pure lines, derived from the hexaploid bread wheat section of the Historic Watkins collection are conserved in the Germplasm Resources Unit in John Innes Centre. Those lines are utilised, and further developed by various institutes, including, Rothamsted Research, Nottingham University and Bristol University, as part of the UKs institute strategic programmes: Wheat prebreeding LOLA, Wheat Institute Strategic Programme (WISP), Designing Future Wheat (DFW) and currently Delivering Sustainable Wheat (DSW).

In 2018, the 827 pure lines (the entire Watkins-derived bread wheat collection) was introduced to China, accompanied by 208 modern wheat cultivars assembled to capture the global modern wheat diversity. This effort marked the initiation of the Watkins Worldwide Wheat Genomics to Breeding project (WWWG2B, https://wwwg2b.com/). Within this initiative, more than one thousand wheat accessions underwent whole-genome re-sequencing, comprehensive phenotyping, and thorough analysis at the Agricultural Genomics Institute at Shenzhen, a part of the Chinese Academy of Agricultural Sciences (CAAS). Of the 827 lines, 118 were originally collected in China one century ago, and hence repatriated as part of WWWG2b collaboration following a century long of ex-situ conservation in the UK. These lines were originated from diverse geographic locations across China, such as Chongteh (West of Tungting lake), Hills near Chungking, Sichuan, Shanghai (Tsao-Ka-Doo), Tung An Men Ta-Chieh, Peking, Mukden, Hankow, Tsinan district, and Tungting lake. At present, 300 core Watkins accessions have been identified as potential parent lines for crossbreeding with various Chinese wheat cultivars including YangMai20/22 and ZhongMai578.

The Watkins collection also includes 354 tetraploid (Durum) wheat, assembled globally in the same manner and at the same time. Following the success of WWWG2B collaboration work on bread wheat, newly generated tetraploid derived lines were also introduced into AGIS CAAS, China, with a vision to couple high resolution genomics and phenomics to enhance durum wheat breeding outputs.

#### 1.2 Germplasm Development and Availability

All the germplasm used in this study is conserved in the UKRI-BBSRC Germplasm Resources Unit (GRU) National Capability at JIC, as well as at Agricultural Genomics at Shenzhen (AGIS), Chinese Academy of Agricultural Sciences (CAAS), a newly established germplasm team is maintaining and developing the WWWG2B germplasms, for its basic research and breeding effort throughout China.

The seed and passport data are now available on https://www.seedstor.ac.uk/ and at Shenzhen. Compiled accession lists were assembled according to the Ancestral Groups (AG1-AG7) reported in this study, representing a landmark in crop germplasm management, relying on high resolution genomics rather than on the accepted historic geographic origin data. https://www.seedstor.ac.uk/search-custom.php. The progenitor landrace populations (Watkins Historic Collection) are available on https://www.seedstor.ac.uk/search-browseaccessions.php?idCollection=4. The derived 827 illumina sequenced Watkins landraces seed stocks are available on https://www.seedstor.ac.uk/search-browseaccessions.php?idCollection=39. The 208 Modern elite wheats used were sourced from various SeedStor collections and can be viewed and ordered collectively as a Compiled Accession List at https://www.seedstor.ac.uk/search-custom.php

Pure stocks of the seed harvested from a single DNA sequenced plant (gold standard stocks) are kept for reference in the GRU and were also maintained and developed at AGIS CAAS, China. Progeny of these were multiplied in greenhouse with bagged ears for preventing cross pollination and handled following international Genebank Standards (Rome 2014). These progeny are available for the global wheat community. Novel plant genotypes deposited in GRU as part of this study include the BBSRC Designing Future Wheat - Recombinant Inbred Lines (RILs) Nested Association Mapping panel (DFW - NAM) comprising 8359 and the DFW Wheat Academic Toolkit pre-breeding germplasm collection, currently comprising 1845 lines. Germplasm or DNA sample can be ordered here:
DFW-NAM: https://www.seedstor.ac.uk/search-browseaccessions.php?idCollection=47; DFW-ATK: https://www.seedstor.ac.uk/search-browseaccessions.php?idCollection=40

The DFW – NAM collection are RILs from a total of 91 bi-parental mapping populations (including the 73 populations used in the study). Each population originates from a cross between the UK spring wheat Paragon and 91 hexaploid wheat landraces, members of a genetically diverse core set of the A.E Watkins Stabilised Collection. The DFW ATK is a set of newly developed pre-breeding germplasm lines containing the next generation of key traits. The ATK includes near isogenic lines (NILs) or equivalent material from different wheat diversity sources, e.g., landraces, mutation screening, ancient wheat relatives and new synthetic wheats. ATK lines are assessed for traits of interest within the DSW programme (http://wisplandracepillar.jic.ac.uk/toolkit.htm) and are genotyped using the 35k Axiom Wheat Genotyping Breeders Array. Particularly promising lines are assessed in multi-site trials by commercial breeders and scientific institutions as an annually changing Breeders Diversity Toolkit (BTK) series. Paragon has been used as a background parent for many lines (Wild Relative, Mutation, Landraces) along with Robigus (Synthetics). Further background parents have been more recently added such as Freiston (F), Gleam (G), Silverstone (S) and Weebill (Wee) Genotype data and genetic maps are available at: http://wisplandracepillar.jic.ac.uk/results_resources.htm and QTL discovered at: http://www.cerealsdb.uk.net/cerealgenomics/CerealsDB/select_QTL.php

**Supplementary Note 2: A WWWG2B database and breeding portal**

Here, we present WWWG2B platform (https://wwwg2b.com), a free, integrative web-based, and user-friendly wheat research and breeding portal. A variety of featured and practical toolkits were provided, offering convenience for the further study of wheat functional genomics, selection of academic toolkits and breeding toolkits. The detailed description of this platform is provided in the following sections.

**2.1 Database structure and content**

Our database mainly includes the following five components. (1) Breeding Strategy: this part mainly described the breeding strategy and key breeding methods. (2) Display of Watkins collection diversity: this part shows the diversity of Watkins wheat germplasm resources, including the descriptions of different Watkins accessions and genotypes, phenotypic characteristics, and possible genetic potential on breeding. (3) Diversity database: this part contains SNP, InDel, Haplotype, and the identified beneficial alleles superior to modern wheat, supporting gene discovery and diversity analysis in wheat breeding research. (4) Breeding and analysis tools: this section integrated breeding and analysis tools, allowing for data processing, data query, statistical analysis, and visualization. (5) People and contact information: this section contains the contact information for members of the breeding team, partners, and the data manager of the WWWG2B project.

**2.2 Data deposition and access**

The WWWG2B contains germplasm information, and the genetic variation information for SNPs, InDels, SVs of all the accessions in this study. In addition, we showed the identified haplotypes for each gene based on SNPs, and all the QTLs and MTAs at multiple locations in multiple years. These various datasets were all organized and stored in MySQL, each with a query web page. For data access and query, we provide a web-based user interface (https://wwwg2b.com). On this interface, users can order, query, browse, and analyze data online. For the data stored in the database, users can specify the query conditions using the specific query interface or tools provided by us to obtain the corresponding results.

#### 2.3 Breeding portal

The breeding portal is the core component of the WWWG2B database, which provides breeder friendly links to the highly complex genomics resources and genetics resources developed through this study. The Breeding portal was divided into three parts: 1) Academic Toolkits; 2) Breeding Toolkits; 3) Mapping Toolkits. The data will continuously be growing in the future. We provide an intuitive interface where breeders can search traits or the relevant QTLs and MTAs of interest, or the target genes, alleles or haplotypes of interest; or breeders can design PCR primers, KASP markers for assisted molecular breeding, or sgRNA for target genome editing. Based on the data and algorithms implemented in the WWWG2B database, the breeders can obtain the results in tables, graphs or visualizations.

#### 2.4 Usage

WWWG2B can be accessed through a user-friendly interface. We provide the 'Manual' in 'Documentation' page for users to access the database. There are many functionalities implemented in WWWG2B, such as IGVBrowse, BLAST, Get Sequence, Interval Tool, ID-Converter, Coordinate transformation, Gene Expression, Primer design such as haplotype-specific KASP assay design, sgRNA design, etc., and more datasets will be stored and new tools to be developed. Here are a few of examples explained below.

***BLAST***

This tool allows users to search nucleotide sequence and proteins sequence in different bread wheat assemblies, including CS Refseq v1.0, v1.1 and v2.1. Max score, total score, query coverage, e value and identity are provided for each alignment. For protein sequence or nucleotides sequence of genes, the annotation information is also provided. Please be minded that, the searching process online might be slow because of alignment of your query sequence to the large wheat genome will require considerable computational resource. In the future, more functionalities will be realized, such as linking the target sequence to the allelic diversity and haplotype clustering across the WWWG2B wheat populations

***Get Sequence***

Users can query by gene ID or genomic intervals. The nucleotide sequence and protein sequence of CS RefSeq v1.0 and v2.1 are provided.

***IntervalTool***

We developed a web interface (IntervalTool) to access the location, functional description, and protein domain of a gene by submitting the gene ID/genomic region.

***ID converter***

There are several different assemblies and corresponding annotations of bread wheat, resulting in different genes IDs for one gene. To make the conversion of gene

IDs between these different versions convenient and efficient, we built a conversion tool (IDConverter) that can convert different versions of gene IDs in 10+pangenome [2], CS1.0[3] and CS2.1 to CS1.1 version.

***Gene Expression***

We have built a gene expression database using 59 RNA-Seq datasets, which were divided into four groups including multiple developmental stages, biotic and abiotic stresses, and others. The expression levels of a specific gene are graphically shown when querying with gene ID(s).

***Primer design tool***

More efforts and time are generally required to obtain sequence-specific primers in polyploid species than in diploid species. We implemented 3 user-friendly and efficient wheat primer design tools (PCR primers Design, KASP primers Design, and sgRNA Design) respectively, which integrated primer design and specificity check into one step. However, what are still in development are the linkage and association analysis of haplotypes for target traits, to empower breeders to obtain useful information, like: 1). What haplotypes are associated with trait of interest; 2). What are the frequencies and distribution of these haplotypes in wheat generally and in Watkins. 3). What specific markers can breeders use to track these haplotypes in breeding.

***IGVBrowse***

Integrative Genomics Viewer (IGV) is a high-performance, easy-to-use interactive tool for the visual exploration of genomic data. We present annotation information for the genomes on this viewer, at this stage, variant information for the 1051 wheat data.

### Supplementary Figures

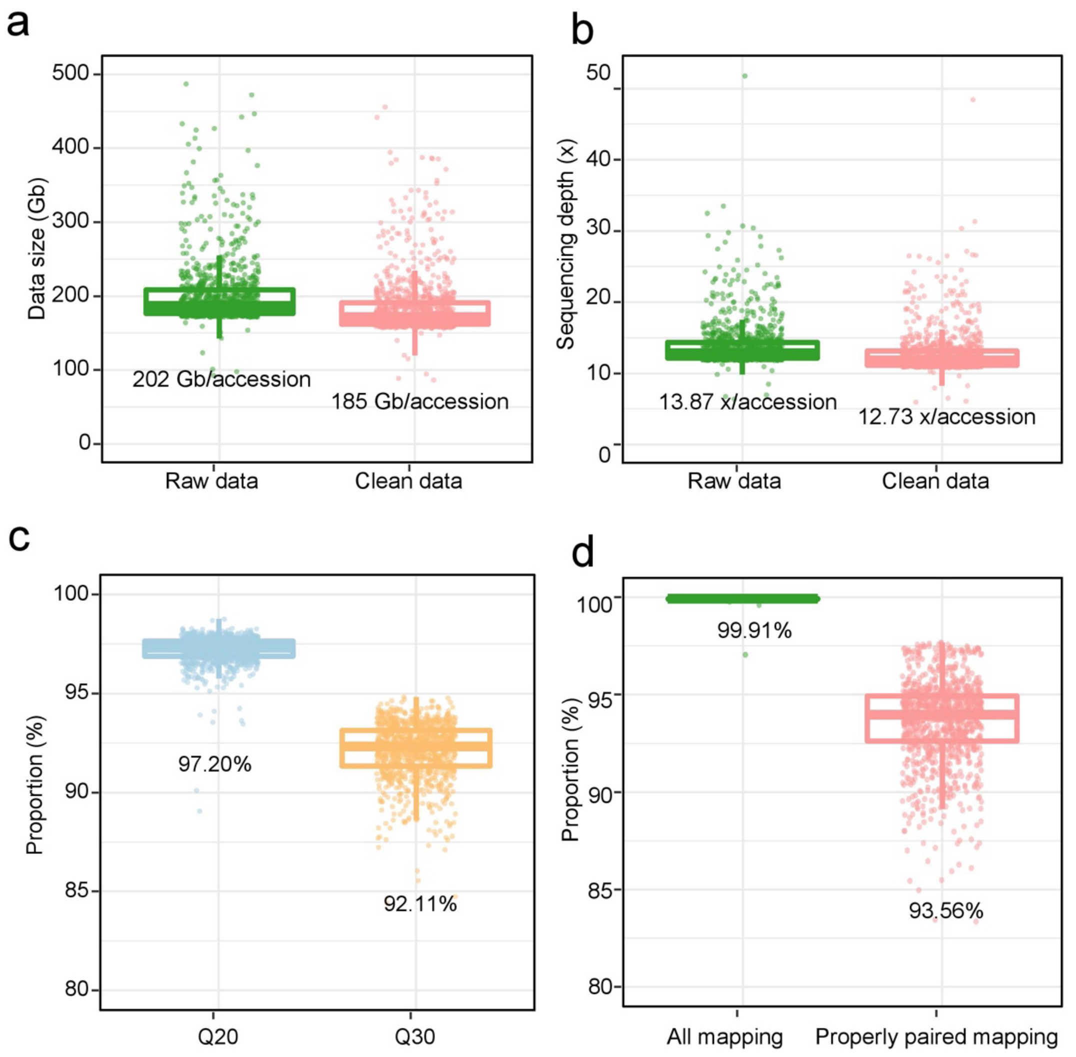

**Supplementary Fig. 1 | Summary statistics of sequencing dataset and quality evaluation for all 1,051 sequenced accessions in this study.** a, The distribution of the total sequenced base number of each wheat accession. b, The distribution of the sequencing depth based on the reference genome size (IWGSC RefSeq v1.0 assembly). c, The distribution of Q20 and Q30 rate of the clean read datasets. d, The mapping rates of 1,051 (including Chinese Spring) accessions based on individual read mapping (all mapping) or properly paired mapped reads. This figure links to Supplementary Table 1.

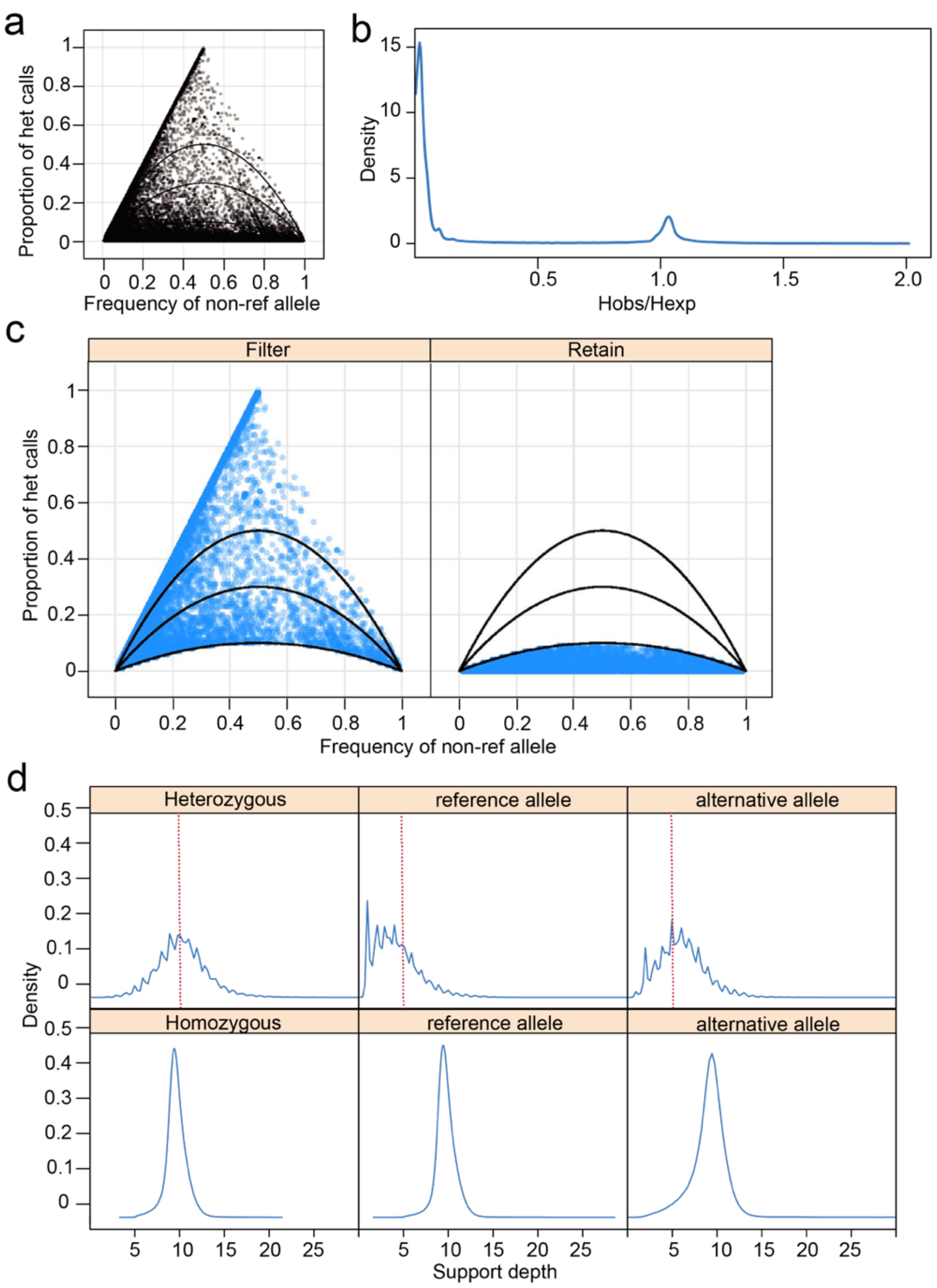

**Supplementary Fig. 2 | SNP quality control through inbreeding coefficient data training**. **a,** Proportion of heterozygous calls versus allele frequency. Each dot represents a SNP from a random sample of 200,000 SNPs on chromosome 3A (5<DP<20, MAF>0.01, Missing<0.1). The three curves show the theoretical values for Hardy–Weinberg equilibrium, inbreeding coefficient of 0.4, and inbreeding coefficient of 0.8. **b,** The distribution of Hobs/Hexp, showing a bimodal peak, as the data with Hobs/Hexp greater than 1 generated mostly due to alignment errors. Hobs is the frequency of heterozygous calls, and Hexp = 2p(1-p), which p is the frequency of non-reference allele (or reference allele). **c,** Comparison of discarded and retained data after filtering using the Hobs_max = 10*(1-Fmedian)*Hexp. The median value of F (Fmedian) for each chromosome is calculated using the SNP site of F>0 and MAF>0.05. **d,** Support depth distribution of heterozygous and homozygous calls of retained dataset. This figure links to Supplementary Table 2.

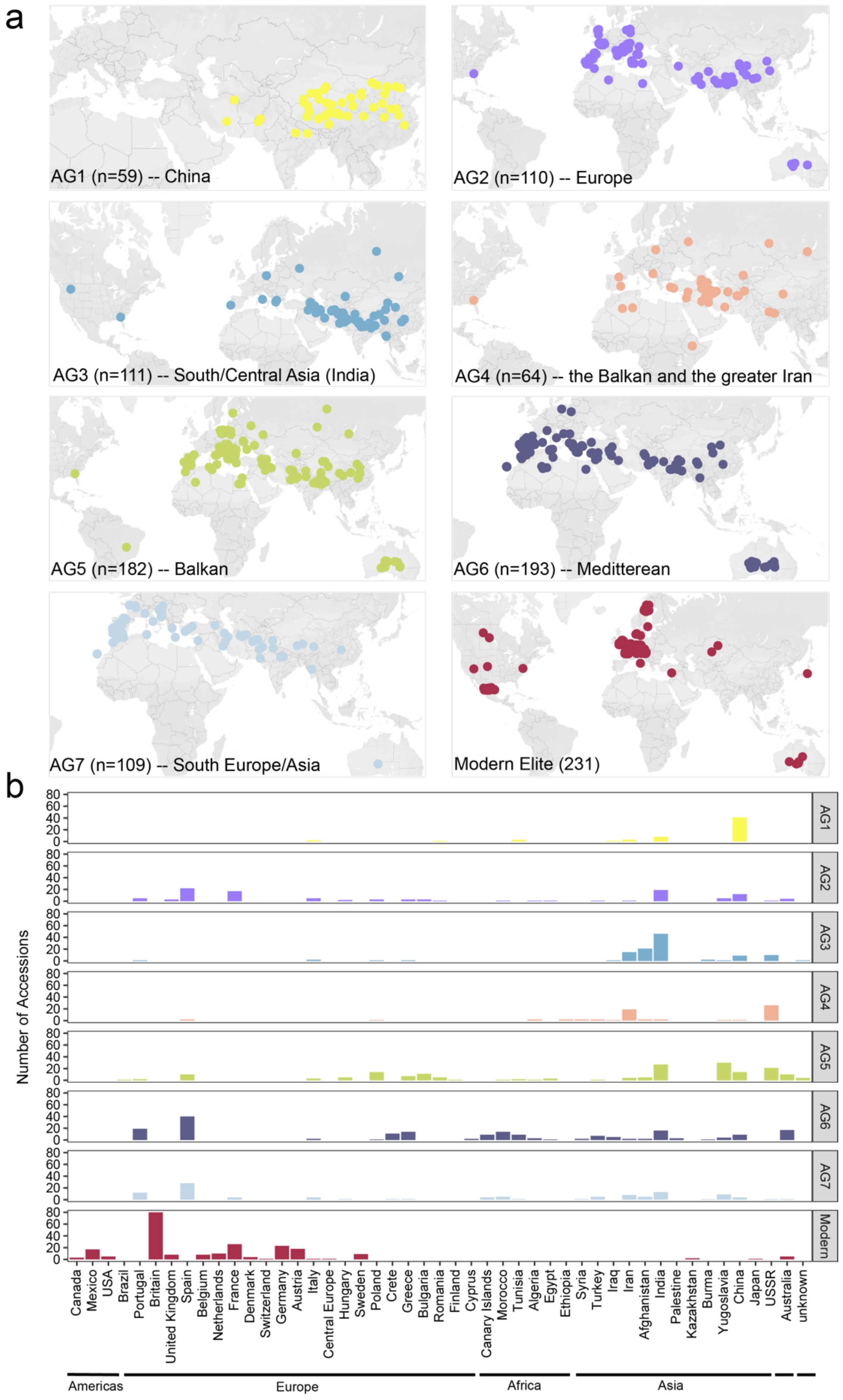

**Supplementary Fig. 3 | Geographic distribution for all the sequenced accessions in this study. a,** World map illustration of the population subgroups (Watkins AG1-7) and Modern cultivars. **b,** distribution of the number of accessions collected from different countries for each population group. This figure corresponds to Fig. 1a.

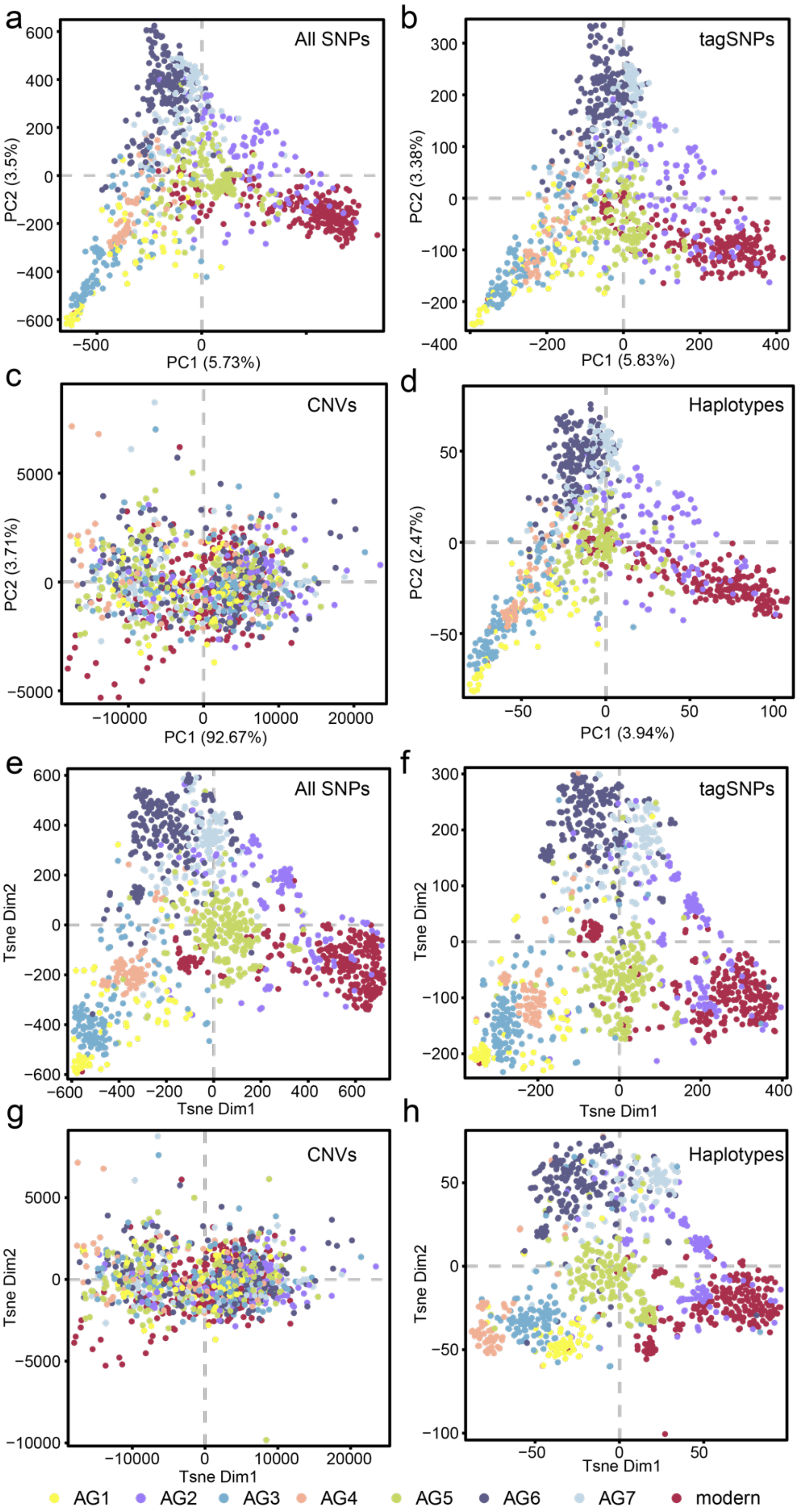

**Supplementary Fig. 4 | The pc1 and pc2 plots of PCA and the dim1 and dim2 plots of tSNE.** a-d, are the PCA results of SNPs, tagSNPs, CNVs, and Haplotypes, respectively. e-h, are the tSNE results of SNPs, tag SNPs, CNVs, and haplotypes, respectively. This figure corresponds to Fig. 1b.

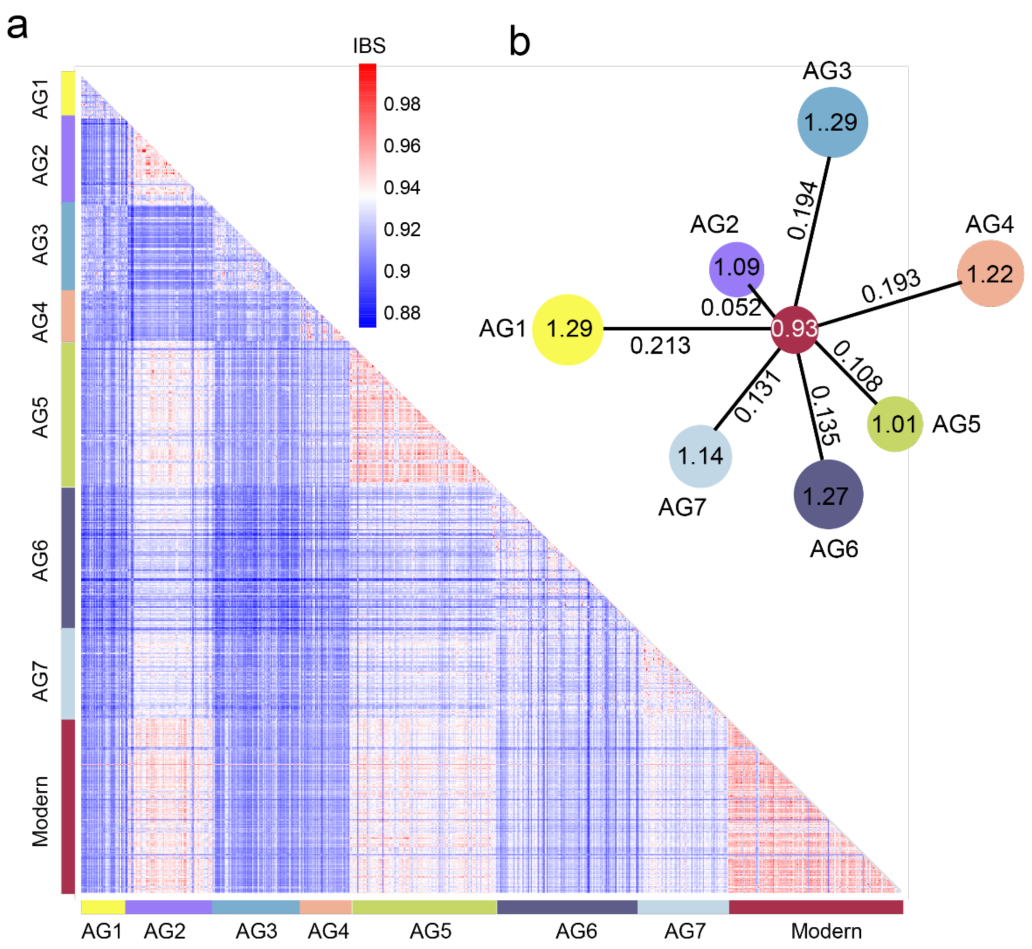

**Supplementary Fig. 5 | The genetic distance between the 7 AGs and modern wheat.** **a,** Genome-wide IBS (identity by state) distance of each pairwise accessions. Values represent the pairwise relationship with red colors representing closer relationships than blue colors. Modern accessions are genetically closer to AG2 and AG5 Watkins groups than to other AGs. **b,** Nucleotide diversity and population divergence (*F*st) across AGs and modern wheat. The value in each circle represents the nucleotide diversity ($\pi \times 10^3$), and the value on each line indicates population divergence between each pairwise population. This figure corresponds to Fig. 1b.

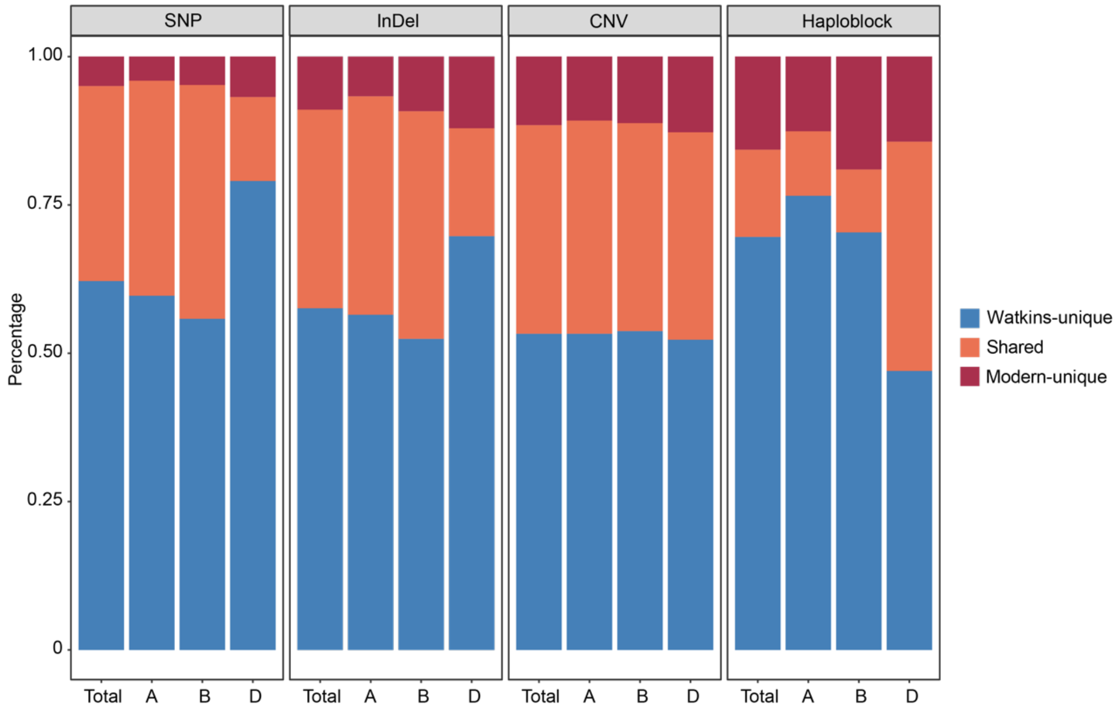

**Supplementary Fig. 6 |** The percentage (%) of the Watkins-unique, Modern-unique and the Shared variations on the whole genome (Total), and the A, B and D subgenomes. **a, b,** and **c,** Percentage of Watkins-unique (the allele frequency of modern =0), shared and Modern-unique (the allele frequency of Watkins =0) SNPs and InDdels. **d,** Percentage of haploblocks without Modern-unique haplotypes (blue), wiht Modern-unique haplotype ≤ 5% (orange), with the percentage of Modern-unique haplotype >5% (red). This figure corresponds to Fig. 1c and Supplementary Table 7.

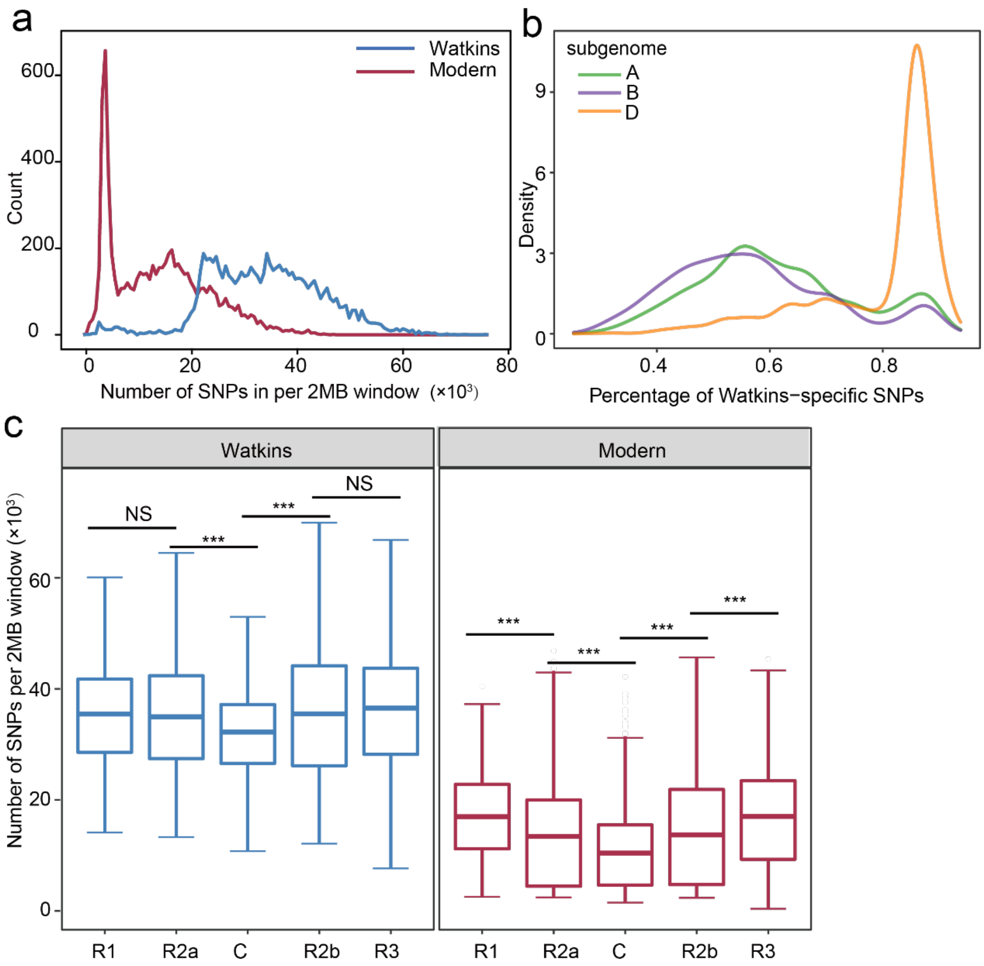

**Supplementary Fig. 7 | Overall summary and comparison of the density and distribution of SNPs between Watkins landraces and modern varieties, between the A/B/D subgenomes and between the five major chromosomic regions.**

**a,** Density distribution and comparison of SNPs between Watkins landraces and modern cultivars. **b,** The percentage of density of Watkins-specific SNPs and compared across the A, B and D subgenomes. **c,** The numbers of Watkins-specific SNPs in the five major chromosomic regions across the genome, R1 and R3 for distal telomeric on short and long arm, R2a and R2b for interstitial regions on short arm and long arm, C for centromere regions. Number of SNPs are calculated in non-overlapping 2MB windows. This figure corresponds to Fig. 1c.

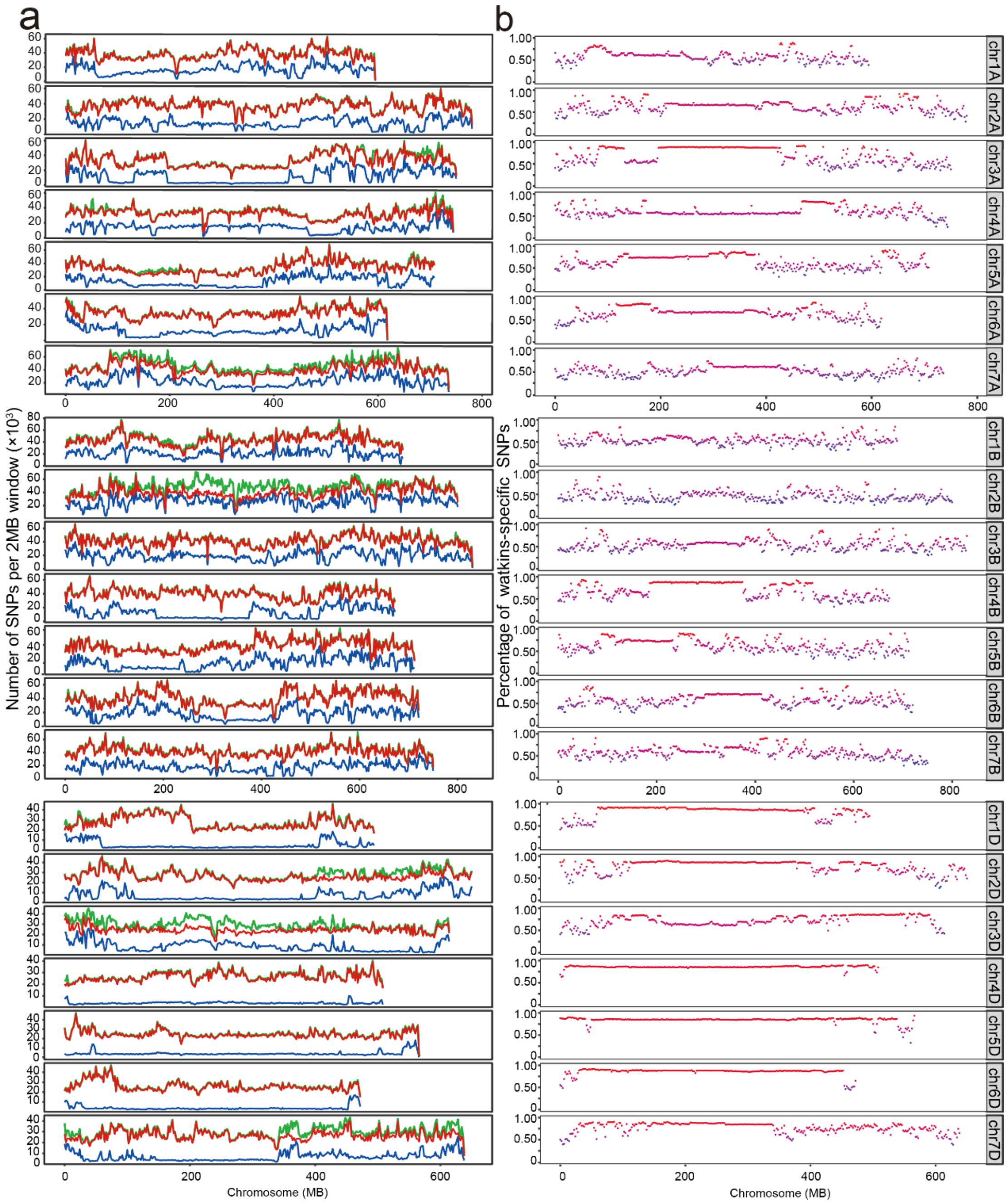

**Supplementary Fig. 8 | Distribution of SNPs and diversity comparison between Watkins landraces and Modern cultivars across chromosomes.** The number of SNPs (x $10^3$) was counted in each 2 MB window along each chromosome. **a,** The distribution of the number of SNPs are colour-coded for Watkins landrace (red), modern cultivars (blue) and all accessions (green). **b,** The percentage distribution of the Watkins-specific SNPs along the 21 chromosomes. This figure corresponds to Fig. 1c.

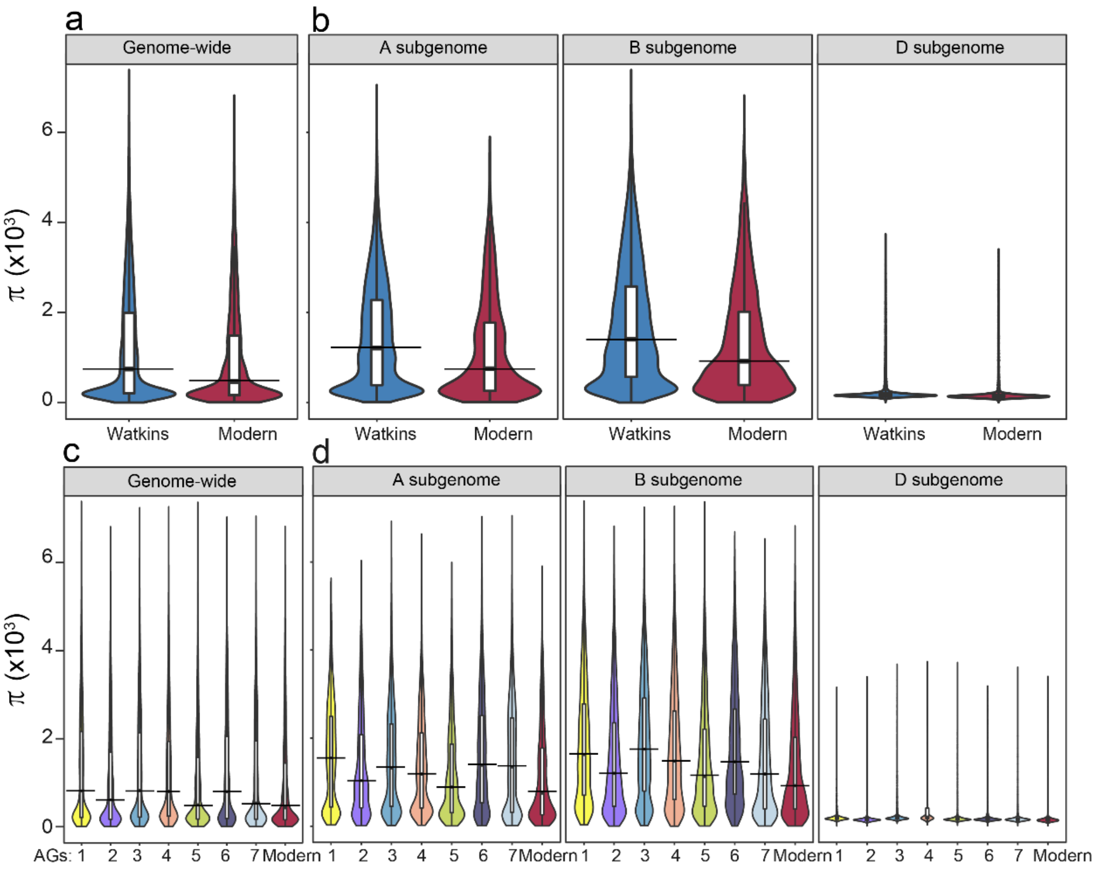

**Supplementary Fig. 9 | Comparison of the nucleotide diversity between Watkins landraces and modern varieties. a,** Genome-wide comparison of nucleotide diversity between Watkins and modern. **b,** Subgenome (A, B, D) comparison of nucleotide diversity between Watkins and modern. **c,** Genome-wide comparison of the nucleotide diversity across the 8 AG populations. **d,** Subgenome (A, B, D) comparison of nucleotide diversity between Watkins and modern. This figure corresponds to Fig. 1c.

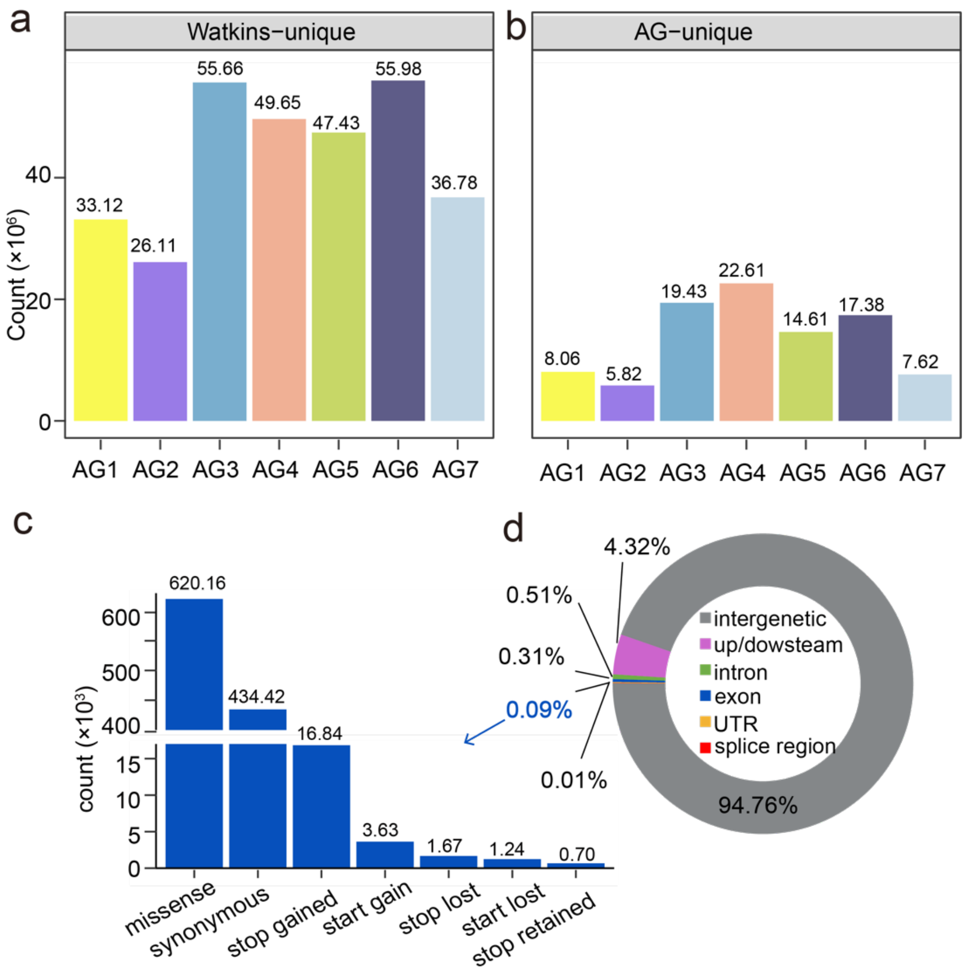

**Supplementary Fig. 10 |** Distribution of count of the Watkins-unique SNPs (**a**) and count of the AG-unique SNPs (**b**) in the seven ancestral subgroups (AG1-7) **c,** Distribution of exon-derived Watkins-unique SNPs, with different functional significance. **d,** Percentage (%) of the Watkins-unique SNPs located in varied genomic regions (intergenic, up/downstream, intron, exon, UTE, splicing regions).

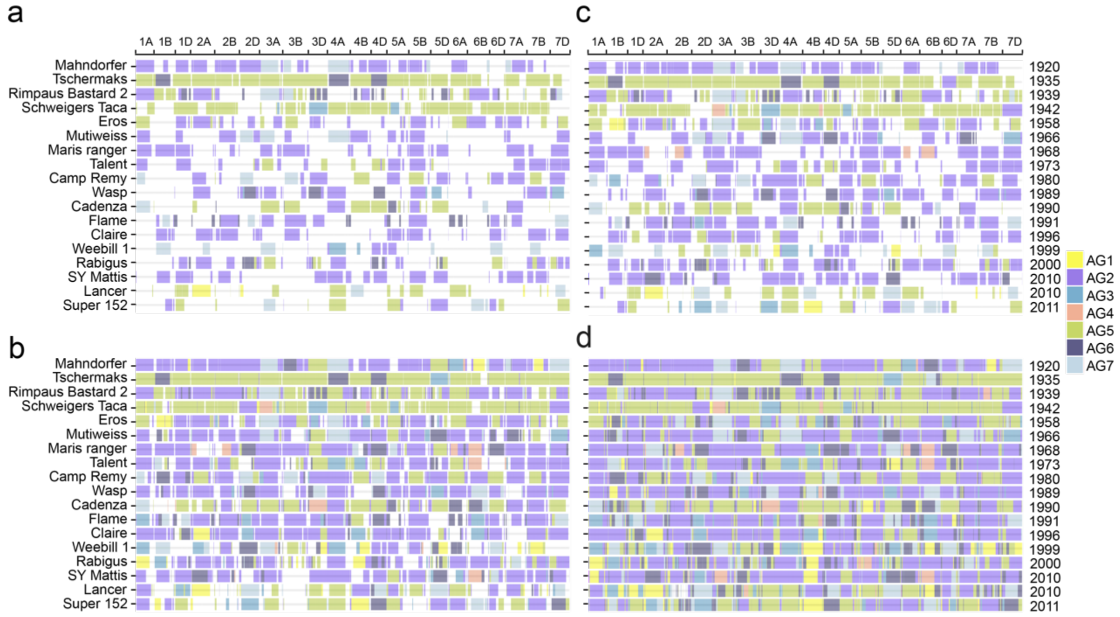

**Supplementary Fig. 11 |** The mosaic structures of the selected 18 modern varieties (20$^{th}$ century) with the conserved genomic composition of the (**a**) top 5, (**b**) top 10, (**c**) top 26 and, (**d**) all of the Watkins landrace accessions. This figure corresponds to Fig. 1d.

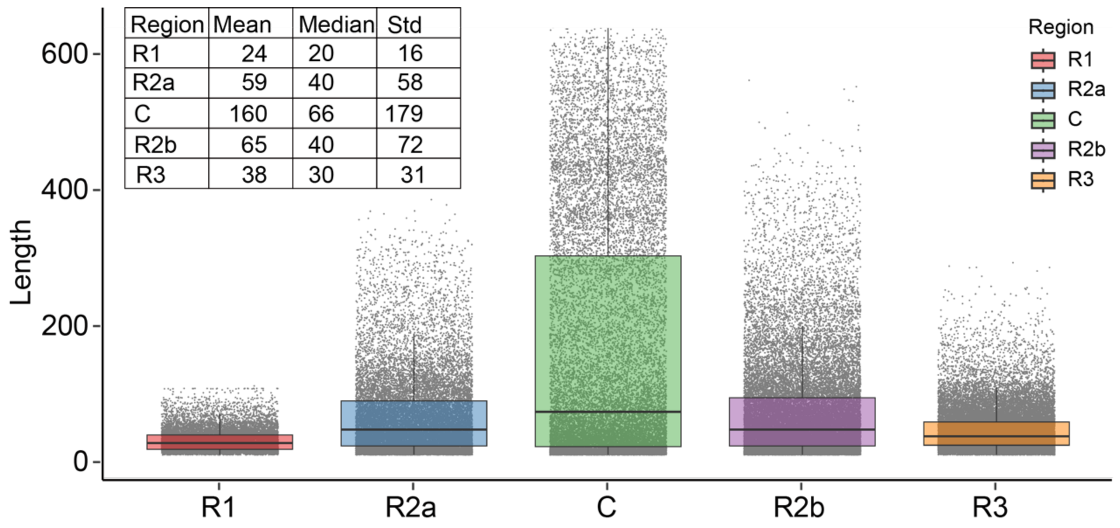

| Region | Mean | Median | Std |
|---|---|---|---|
| R1 | 24 | 20 | 16 |
| R2a | 59 | 40 | 58 |
| C | 160 | 66 | 179 |
| R2b | 65 | 40 | 72 |
| R3 | 38 | 30 | 31 |

**Supplementary Fig. 12 | Distribution of the intact IBS segments across the different five major chromosome regions (R1, R2a, C, R2b, and R3).** The centromeric regions maintain longer segments whereas the more recombinogenic distal regions maintains shorter IBS segments. The mean and median values of the segments is shown in the top left hand table (Std: standard deviation). This figure corresponds to Fig. 1d.

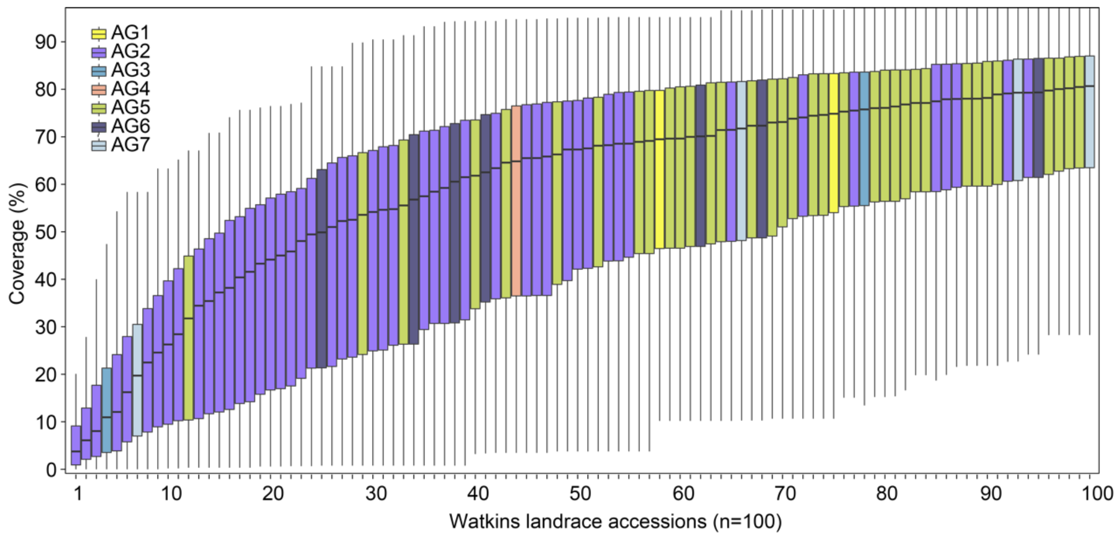

**Supplementary Fig. 13 | The cumulative contributions of Watkins to modern wheat.** The percentage of the Top100 Watkins accessions which contribute to each modern accession was calculated. The boxplot shows the median cumulative contribution, for example WATDE0352 (#1) is responsible for a median of ~3.75% of any modern line, then add WATDE0791 (#2) and this goes up to 6%. The names of the Top100 Watkins accessions are provided in Supplementary Table 10.

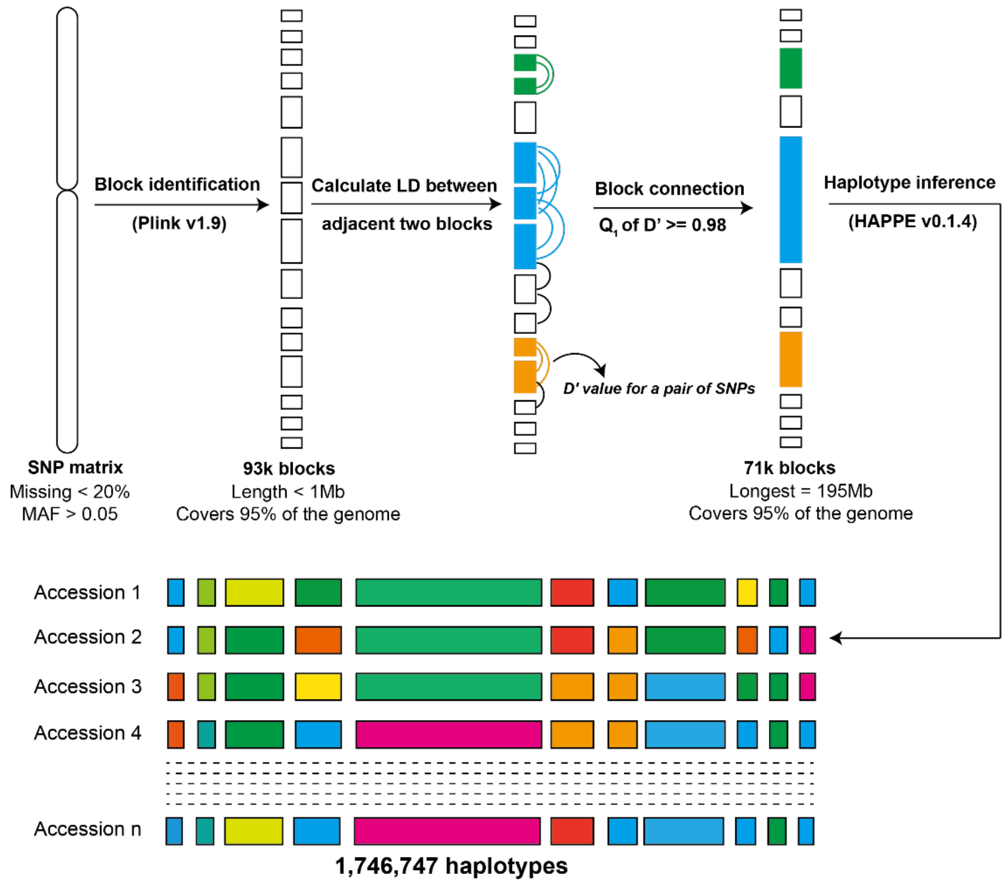

**Supplementary Fig. 14 | Flowchart for the construction of Wheat haplotype block map (HapMap) based on linkage disequilibrium (LD).** HAPPE[4] was a tool developed to visualize the allelic diversity and haplotype clustering. This figure corressponds to Fig. 1e.

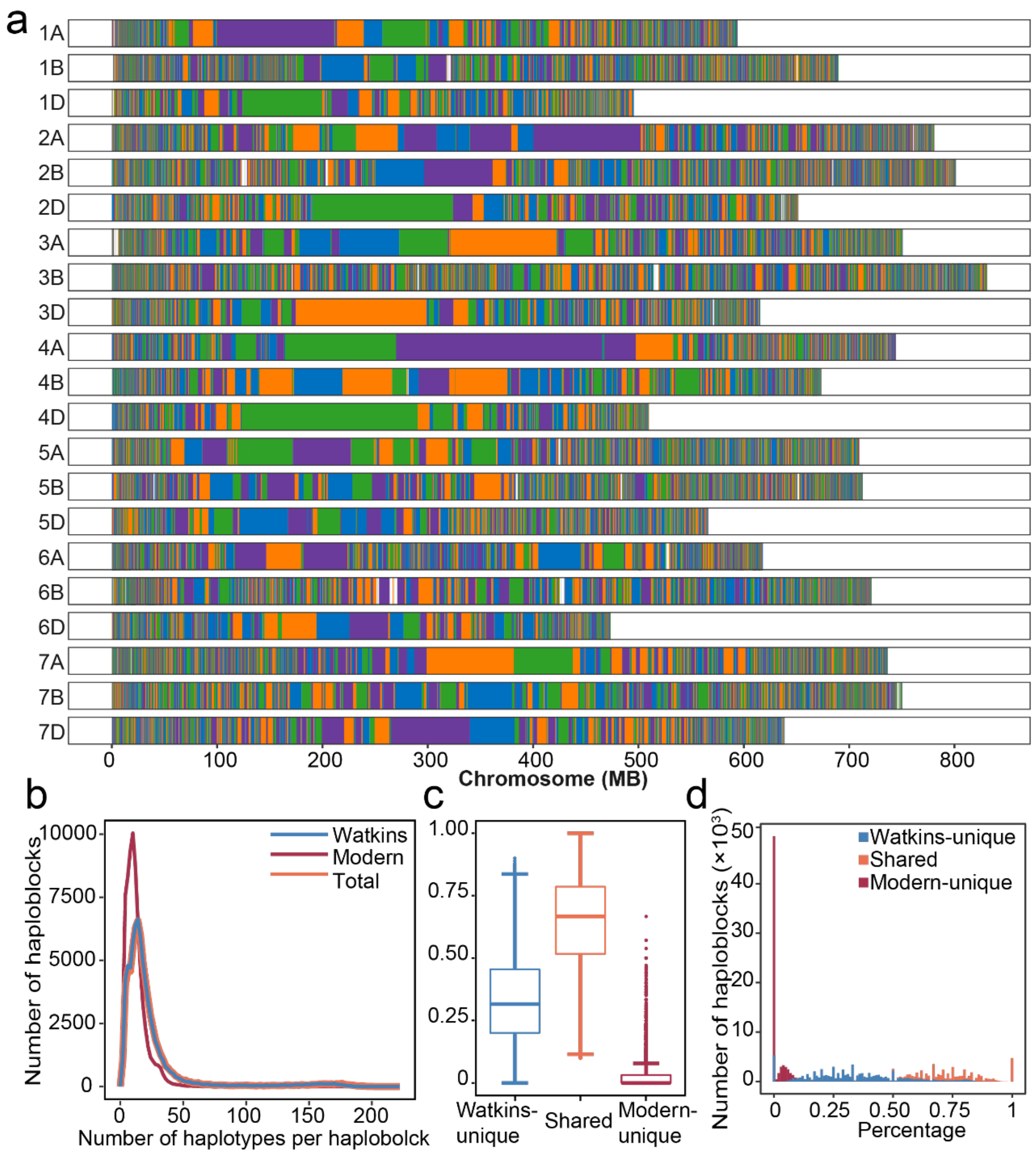

**Supplementary Fig. 15 | The distribution of the 71,282 haploblocks along the 21 chromosomes, and comparison of haplotype diversity between Watkins landraces and modern cultivars. a,** Each chromosome is colored in segments representing a haploblock estimated in this study. **b,** The distribution of number of haplotypes. **c,** Percentages of population-specific and shared haplotypes. Each dot represents a haploblock. **d,** Distribution of percentages of population-specific and shared haplotypes.

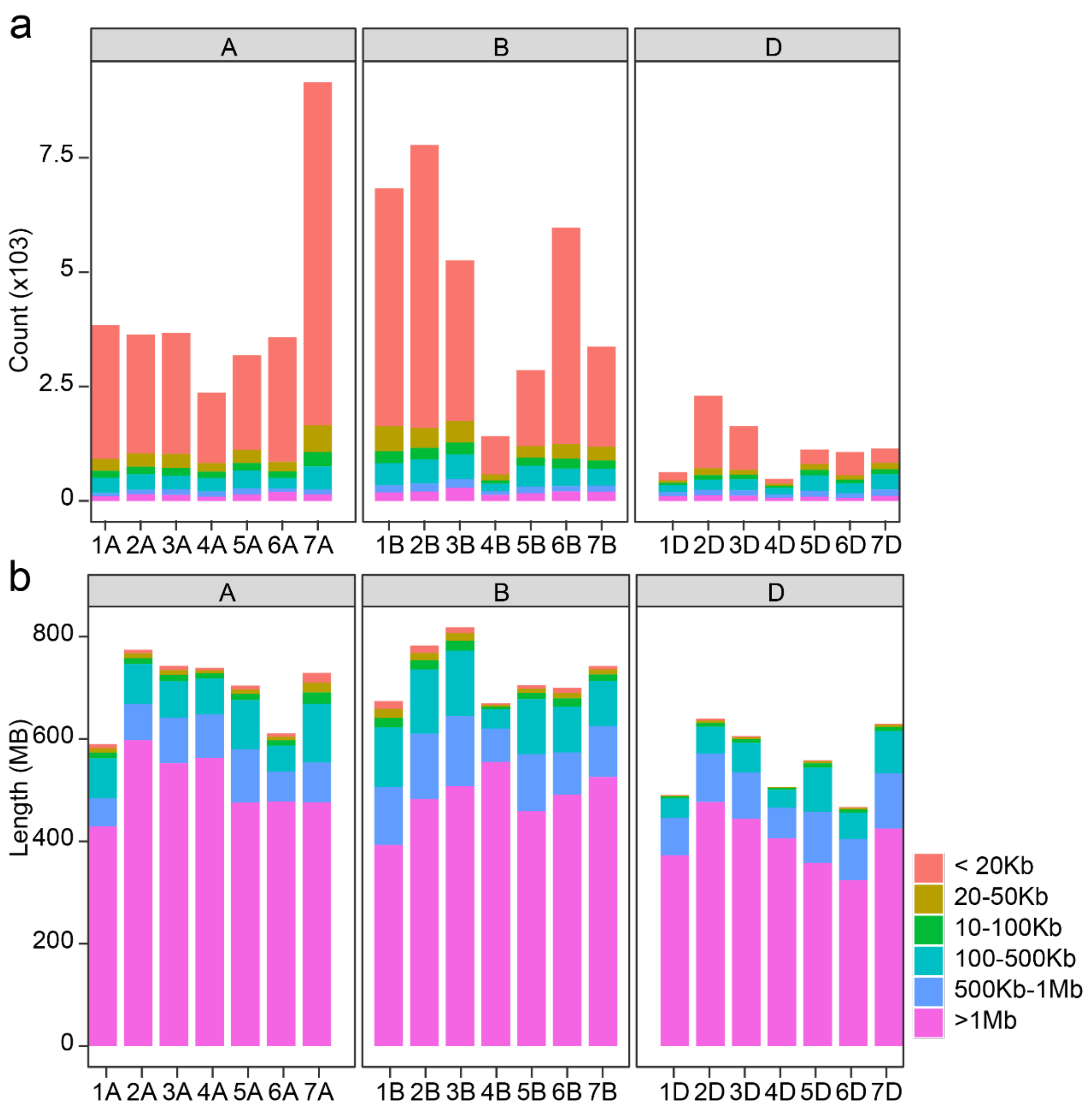

**Supplementary Fig. 16 |** The distribution of the total number (**a**) and the total length (**b**) of different haploblocks, categorized by size, across the 21 wheat chromosomes.

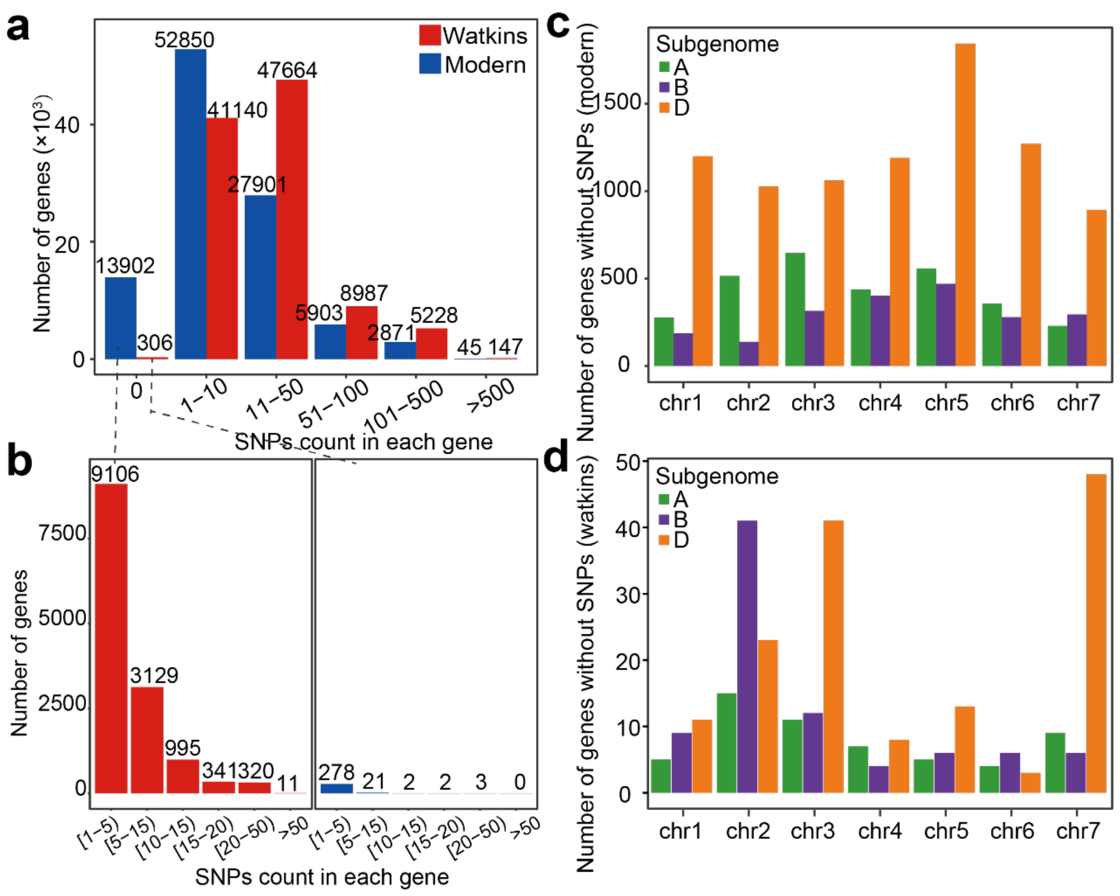

**Supplementary Fig. 17 | The summary of genetic diversity and SNP distribution for the 107,892 high-confidence wheat genes. a,** Genes grouped according to the number of identified genic SNPs. **b,** The distribution of the number of SNPs for Watkins landrace in 13,902 genes that are monomorphic in modern varieties (left), and the number of SNPs for modern varieties in 306 genes that are monomorphic in Watkins landraces (right). **c,** The chromosomal and subgenomic distributions of 13,902 monomorphic genes in modern varieties. **d,** The chromosomal and subgenomic distribution of 306 monomorphic genes in Watkins landraces.

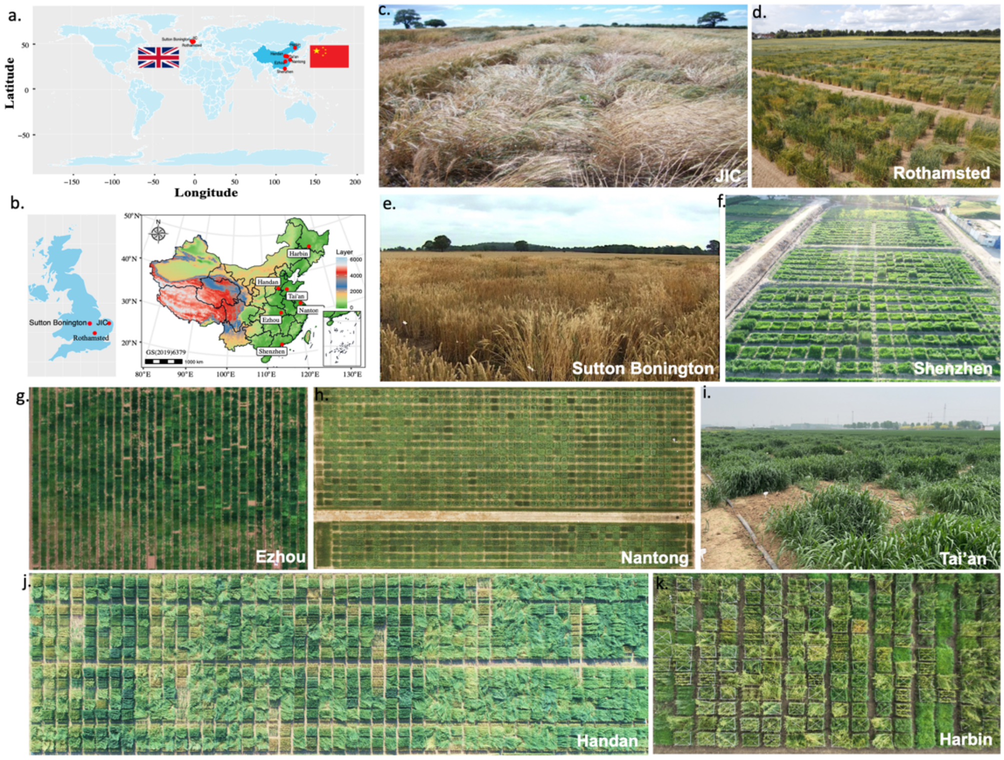

**Supplementary Fig. 18 | Field trial and phenotyping of the Watkins collection. a-b,** Locations of experimental stations across the UK (**c-e**) and China (**f-k**).

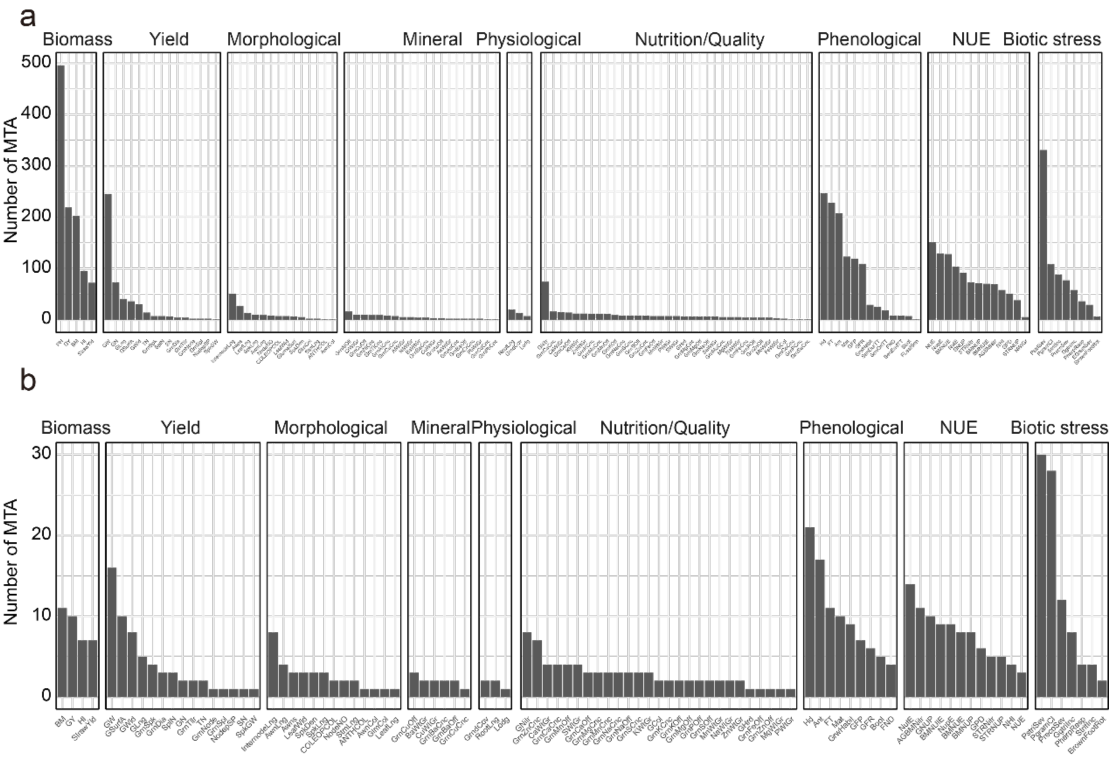

**Supplementary Fig. 19 |** The number of genetic effects (significant peaks) associated with 137 traits, the marker-trait associations (MTAs) detected from GWAS/NAM GWAS, signal peaks as shown from the Manhattan plots. **a**, all MTAs. **b**, prioritized MTAs.

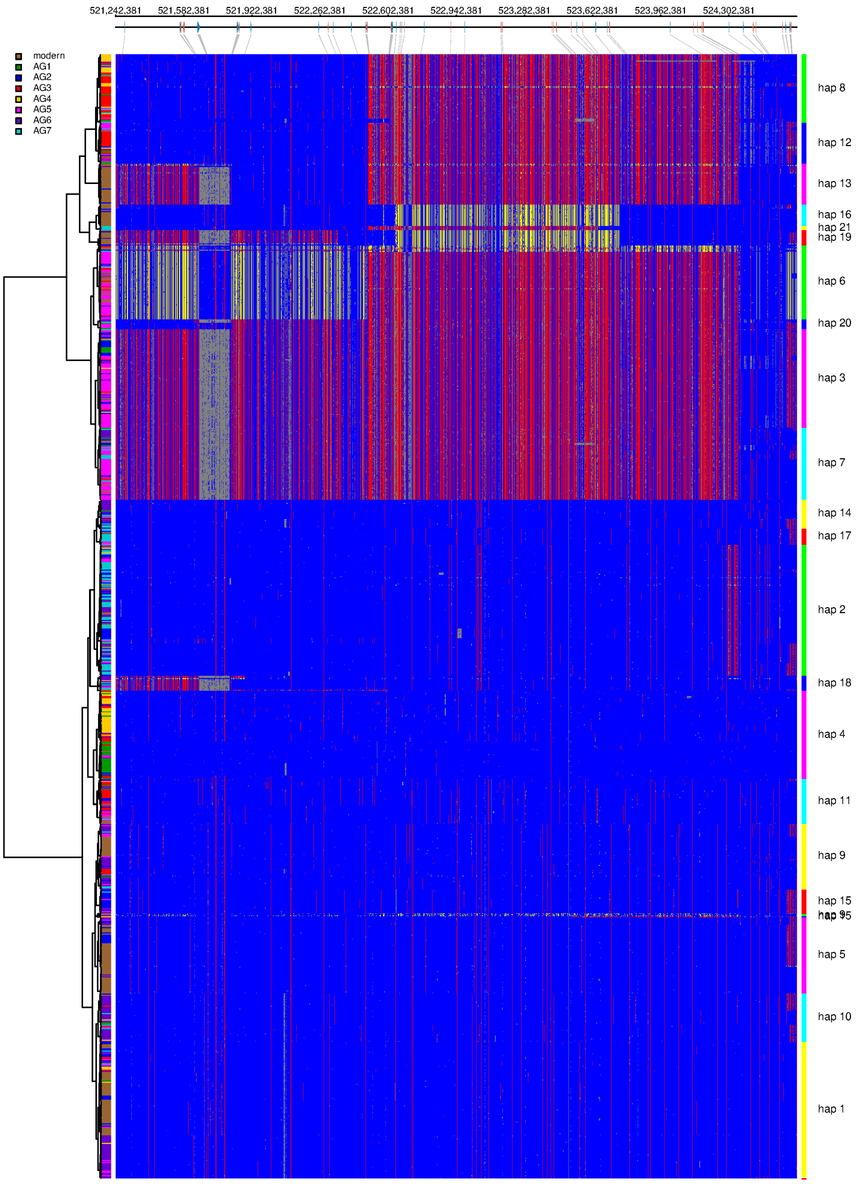

**Supplementary Fig. 20** | An example of the allelic diversity and haplotype clustering across the 1051 accessions for candidate genomic region responsible for cold resistance of the *CBF* cluster on chromosome 5A. HAPPE[4] was used for the haplotype identification and clustering. Blue: reference alleles; red: alternative alleles; yellow: heterozygous alleles; grey: missing. The bars along the phylogenetic tree (left) were color-coded based on the seven ancestral groups (AG1-7) and the modern wheat. The right side indicates the different haplotype clusters (hap: haplotype clusters).

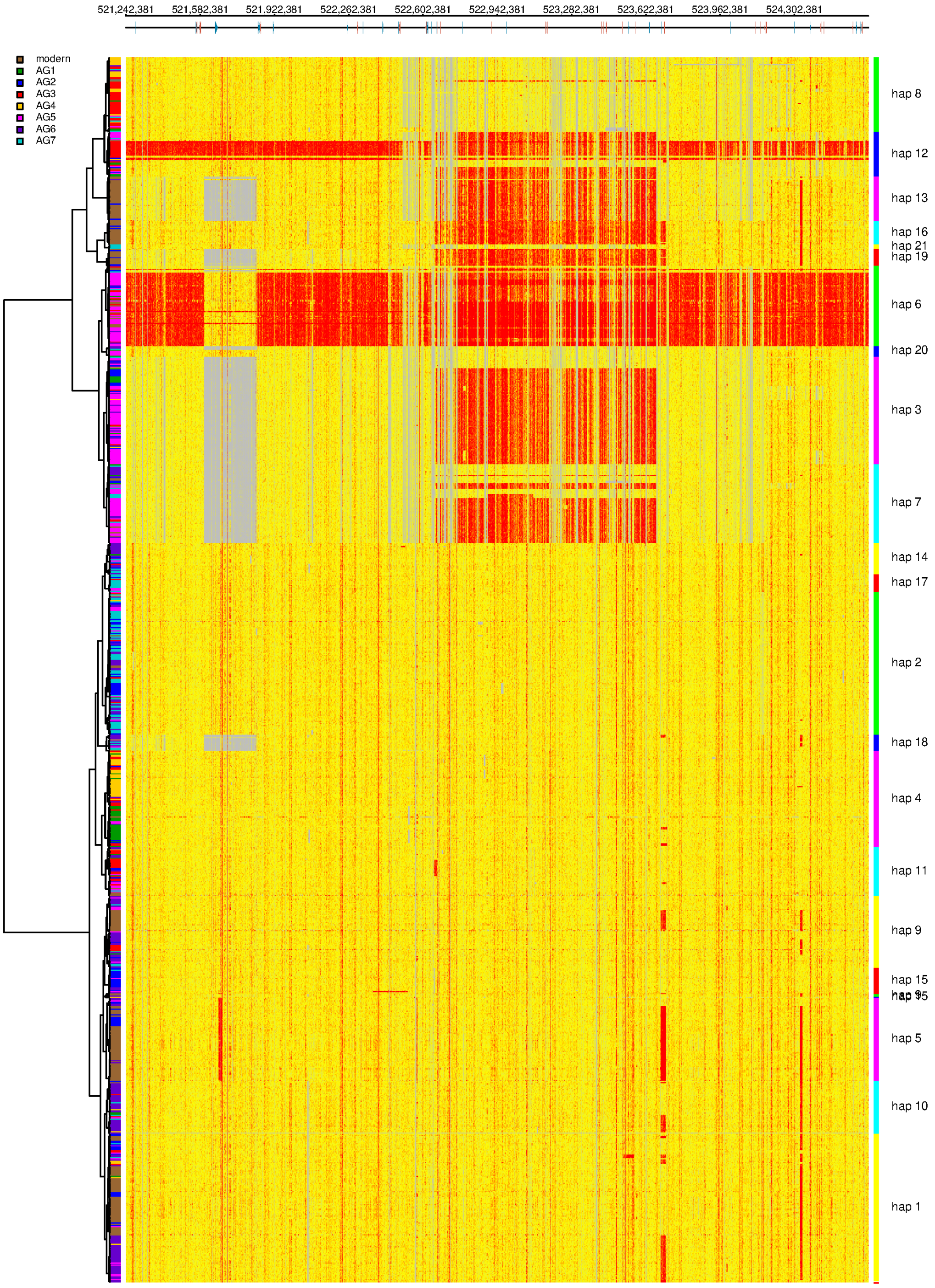

**Supplementary Fig. 21** | The read depth variation across the *CBF* cluster candidate genomic region on chromosome 5A underlying cold resistance. This was used to estimate the copy number variation across the 1,051 accessions. Accessions are sorted based on the haplotype clusters. Gene copy number was color-coded in the heatmap, grey: 0; yellow: 1; red: >=2.

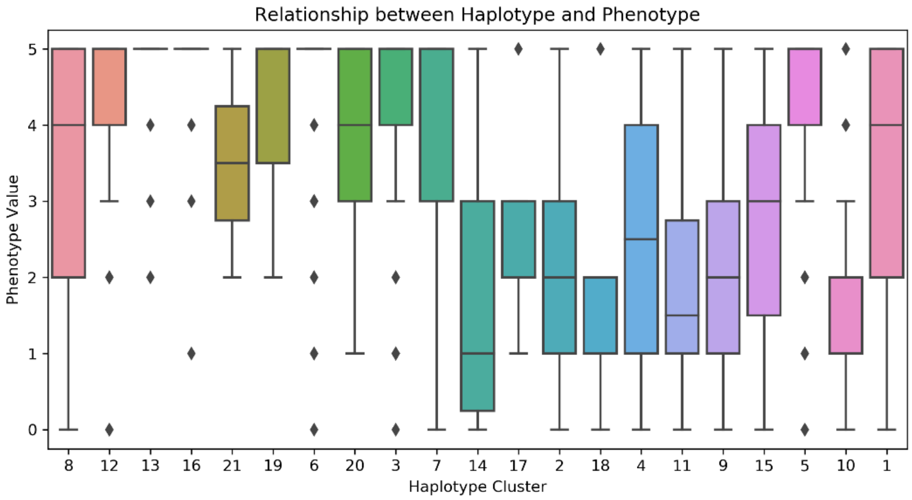

**Supplementary Fig. 22 | Association between the haplotypes of CBF cluster genes and phenotypic variations. Haplotypes with multiple copies of the CBF gene are more resistant to cold.** The higher the value, the better performance in frost resistance. All of the accessions of hap 12, 13, 16, 21, 19, 6, 3, 7, and part of the accession of hap 5 showed multiple copies of CBF gene. This figure corresponds to Supplementary Table 38.

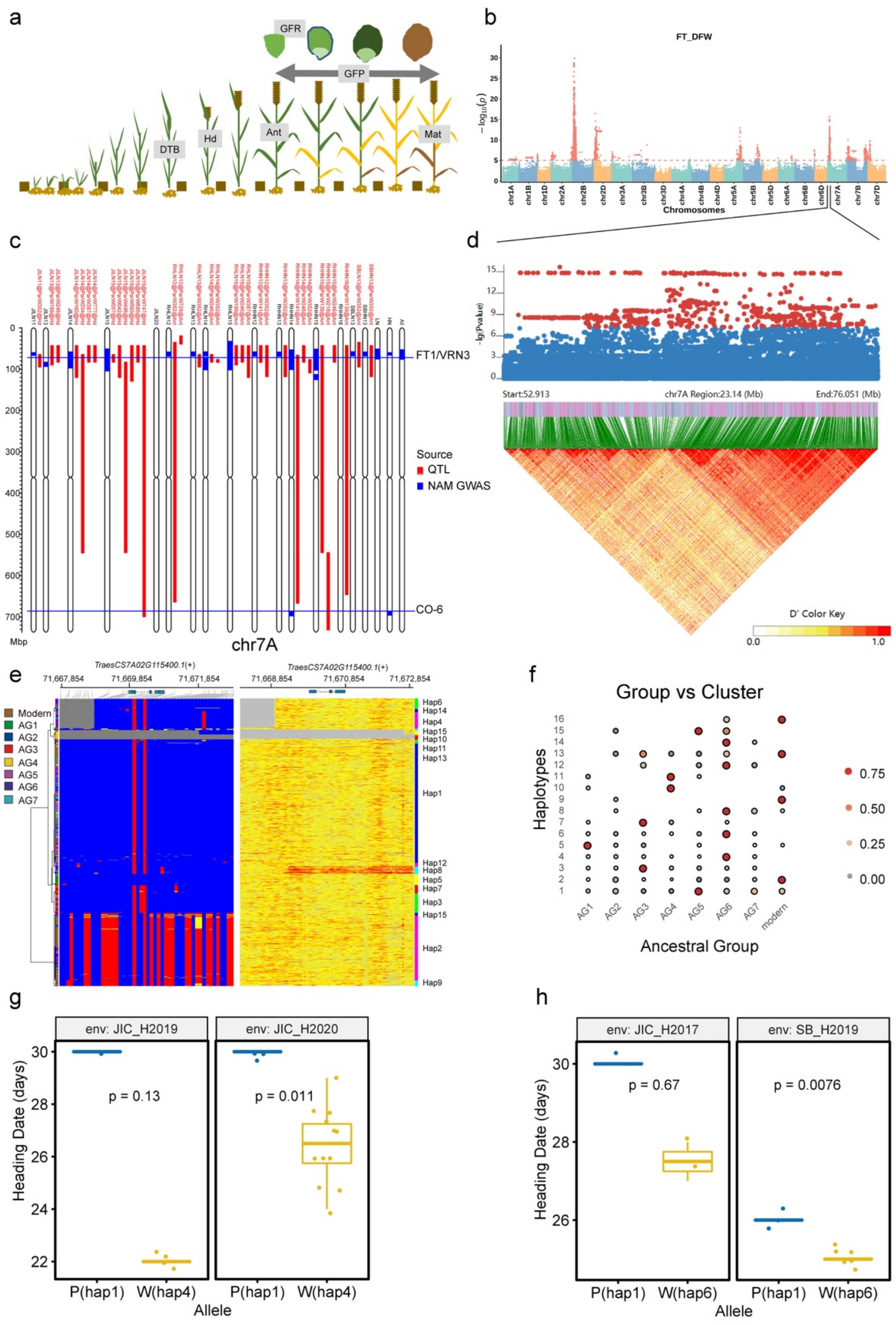

**Supplementary Fig. 23 | An example for gene discovery (*FT1*) of the flowering time related traits, similar analyses were performed for other major traits (see WWWG2B website, the genetic resources: Supplementary Fig. 24 | An example for allele discovery (*FT1*) of the flowering time related traits**. **a,** Schematic illustration of the wheat developmental stages with key transitions in flowering time (booting, heading, flowering and anthesis). **b,** NAM GWAS for the flowering time trait, an example for one of the significant peaks from chromosome 7A (Chr7A) was selected to narrow down into the local details. **c,** All the QTL and MTA from NAM GWAS detected in Chr7A are shown; two known genes located in the overlapped intervals are marked: *FT1* and *CO-6*. **d,** The local LD for the candidate genomic interval in Chr7A overlapping with *FT-A1*. **e,** Visualization of the allelic diversity, haplotype clustering, and read depth variations for the candidate gene *FT-A1* and its up-/down-stream 2 kbp regions; **f,** The distribution and frequency of the different haplotypes clustered in **e**, across AG1-7 and modern wheat. **g.** phenotypic effect confirmation by RIL population in which a prioritized QTL segment from WATDE0002 that carries the *FT-A1* Haplotype 4 (hap4, promoter deletion) was introgressed into Paragon (hap1), leading to earlier flowering. **h,** Phenotypic effect confirmation by RIL population in which the second prioritized QTL segment from WATDE0038 that carries the *FT-A1* hap6 (promoter deletion) was introgressed into Paragon (hap1), leading to earlier flowering. Similar analyses were performed for other major traits (see WWWG2B website, the genetic resources: https://wwwg2b.com/).

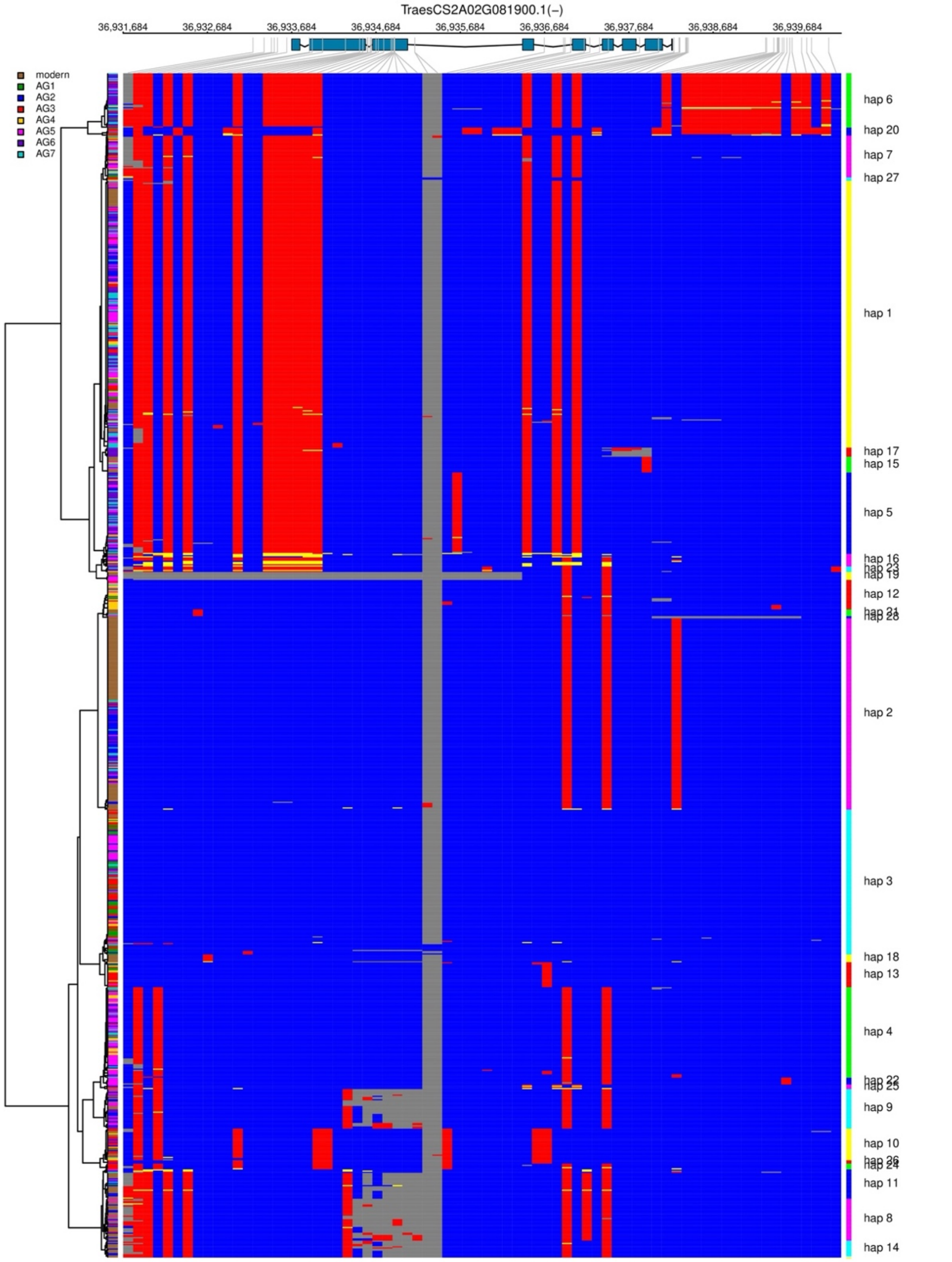

**Supplementary Fig. 24 | Allelic diversity and haplotype clustering across the 1051 accessions for *Ppd-A1* on chromosome 2A.** HAPPE[4] was used for the haplotype identification and clustering. Blue: reference alleles; red: alternative alleles; yellow: heterozygous alleles; grey: missing. The bars along the phylogenetic tree (left) are color-coded based on the seven ancestral groups (AG1-7) and modern wheat. The right side indicates the different haplotype clusters (hap: haplotype clusters).

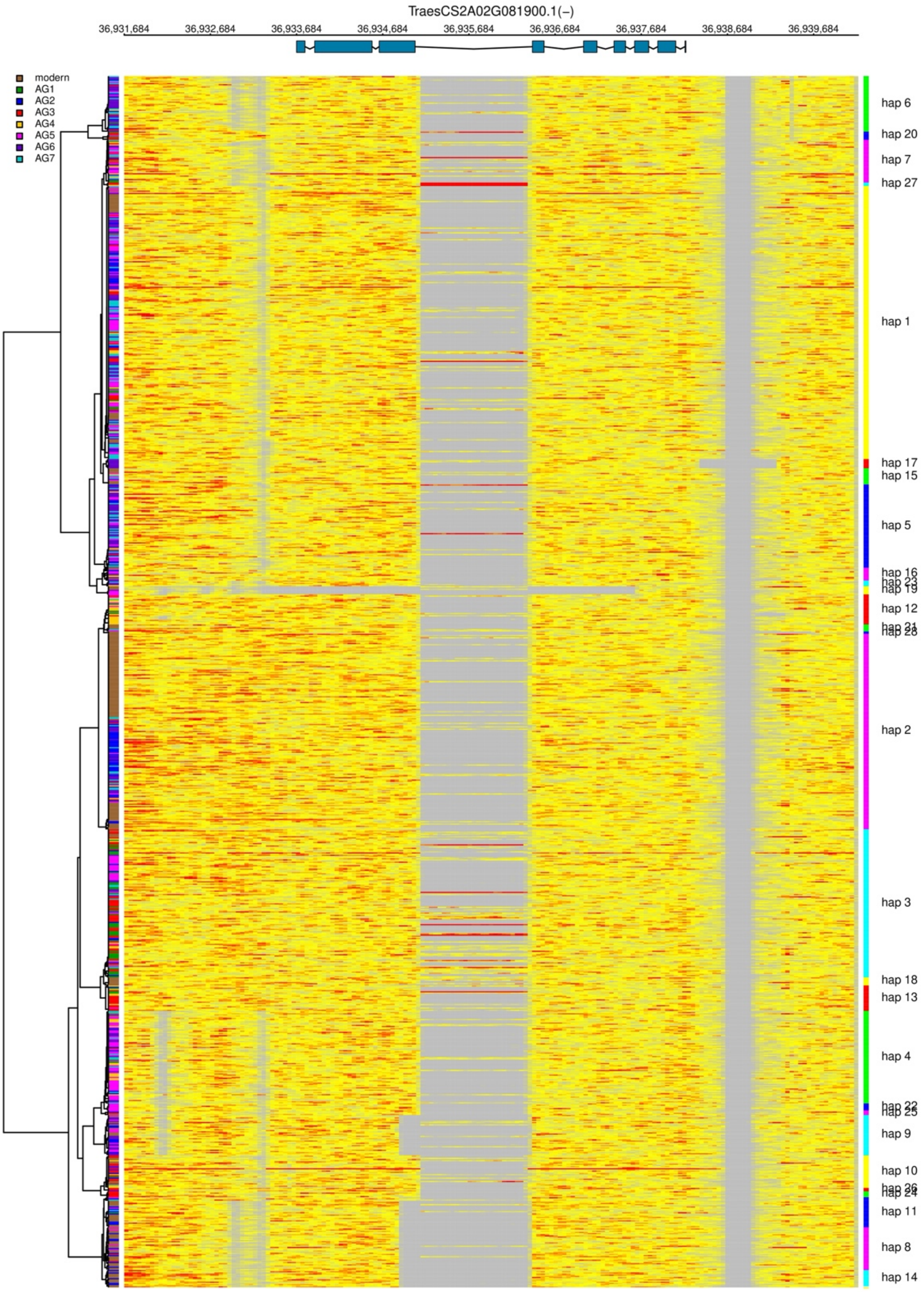

**Supplementary Fig. 25 | The read depth variation of *Ppd-A1* on chromosome 2A.** This was used to estimate the copy number variation across the 1,051 accessions. Accessions are sorted based on the haplotype clusters. Gene copy number was color-coded in the heatmap, grey: 0; yellow: 1; red: >=2.

a

b

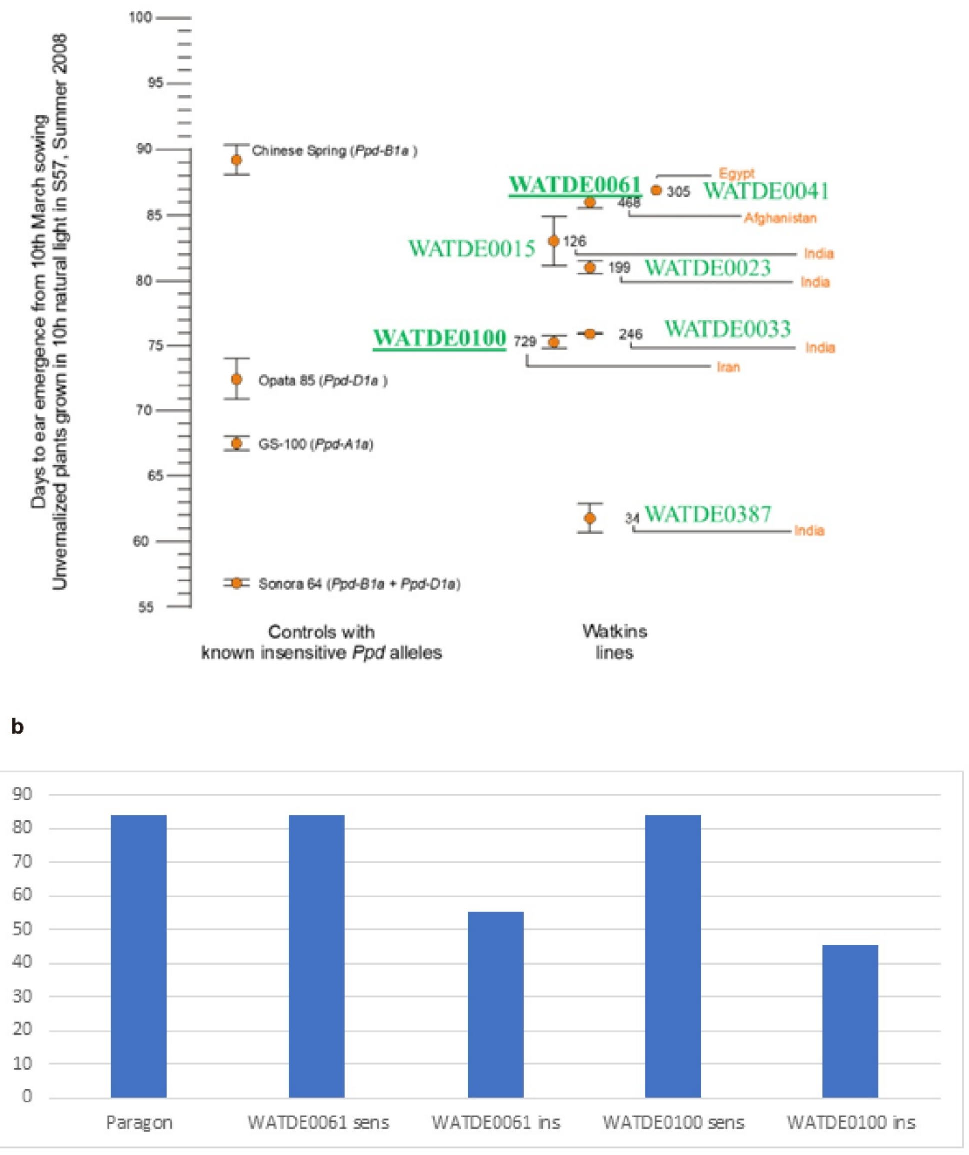

**Supplementary Fig. 26** | **a**. Early flowering Watkins lines were tested under short days and found to exhibit photoperiod insensitivity, but they did not carry known photoperiod insensitivity alleles. QTL analysis mapped these earliness effects to the same location as PPD1 (see Supplementary Table 24). **b**. In the case of WATDE0061 and WATDE0100 the QTL was located on chromosome 2A. Near Isogenic Lines carrying putative *PpdA1a* alleles were developed (see Supplementary Table 25) which were then grown under 10-hour day length in controlled environment chambers.

Paragon and NILs carrying photoperiod sensitive alleles never reached heading. Whereas NILs carrying the WATDE0061 insensitive flowers significant earlier by 21.8 days ($p < 0.02$ ) and WATDE0100 insensitive flowers earlier by 21 days ($p < 0.07$). The experiment was stopped on August 22$^{nd}$.

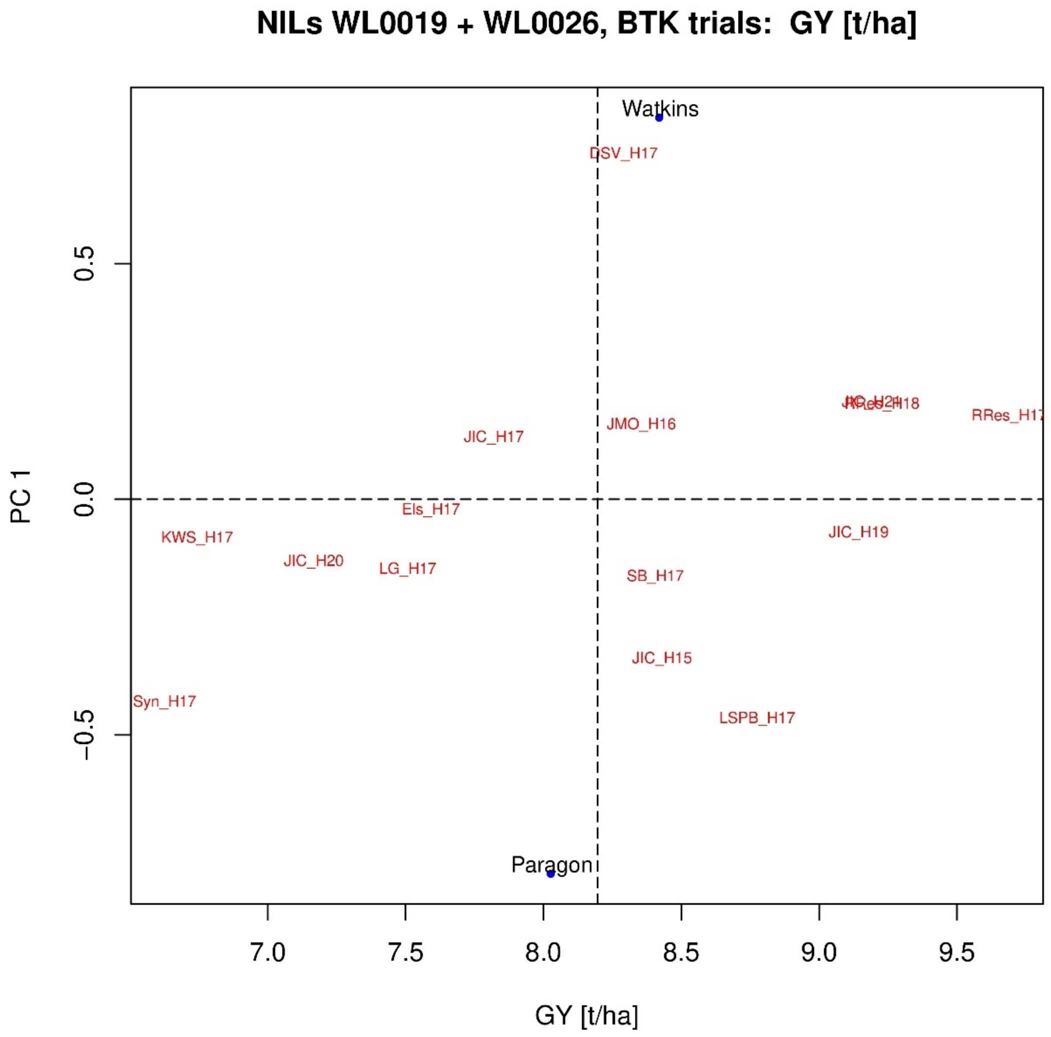

**Supplementary Fig. 27 | 7B-PH Additive Main-effects and Multiplicative Interaction (AMMI) means for GY.** AMMI bi-plot showing the GY potential of a NIL pair developed for a PH increasing QTL on chromosome arm 7BL in WATDE0018 as measured in fifteen field experiments. The GY estimator, in t ha$^{-1}$ (t/ha), is plotted against PC1 of the AMMI, also demonstrating the environmental variability of NILs and sites. The GY estimators of the two NILs are shown as a blue dot with the NILs named by the origin of the introgressed allele, either Watkins (NIL WL0019) or Paragon (NIL WL0026). The Watkins allele shows a GY advantage over the Paragon allele of 0.39 t/ha. The GY potentials of the experiments are plotted in red, using the abbreviation for site and harvest year. This figure links to Figure 3g.

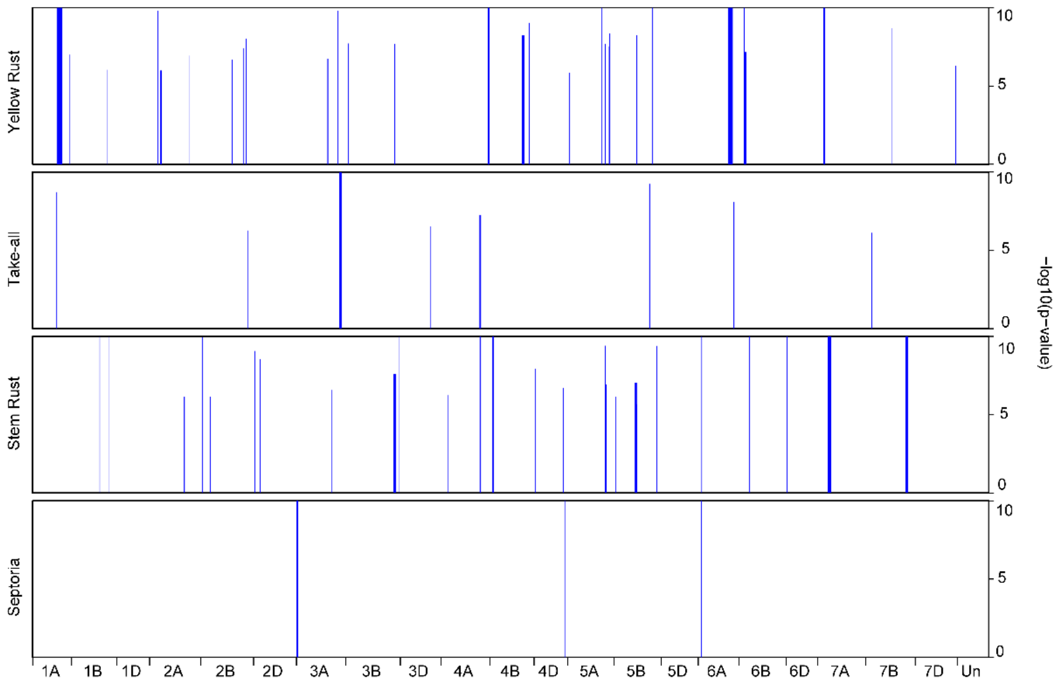

**Supplementary Fig. 28 | The distribution of 70 prioritized MTAs for major diseases, including yellow rust, take-all, stem rust, and Septoria.** The width of the blue bars represents the physical size of each MTA, while the height indicates its significance level, quantified as -log(p-value). For visualization purposes, all -log(p-values) greater than 10 are capped and displayed as 10.

| phenotype of lines* | haplotype tag KASP | trait categories | phenotype of lines* | haplotype prediction | prediction accuracy |
|---|---|---|---|---|---|
| **Coleoptile colour** | CC_083 | green | 156.00 | 151.00 | 96.79 |
| | | red | 50.00 | 42.00 | 84.00 |
| | | Total | 206.00 | 193.00 | 93.69 |
| **Awns** | AWN_744 | Absent | 173.00 | 134.00 | 77.46 |
| | | Present | 177.00 | 172.00 | 97.18 |
| | | Total | 350.00 | 306.00 | 87.43 |

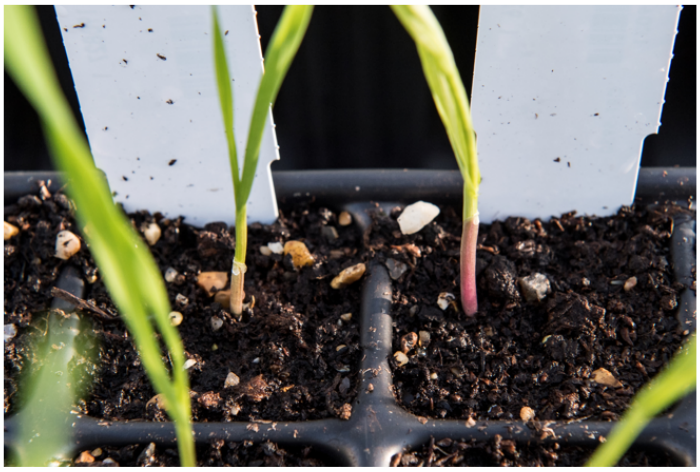

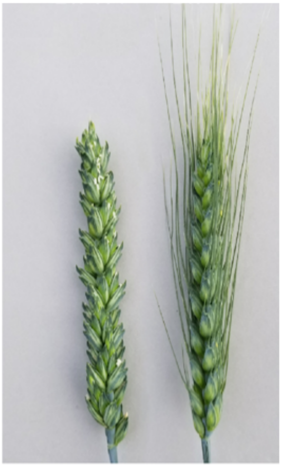

**Supplementary Fig. 29 | Phenotypic prediction using haplotype** tag KASP markers**.** A table describing the prediction accuracy by trait category count and % of accurate predictions for each of the markers used. In the lower panel, a visualisation of the two assessed traits, coleoptile colour (left) and presence/absence of awns (right).

.

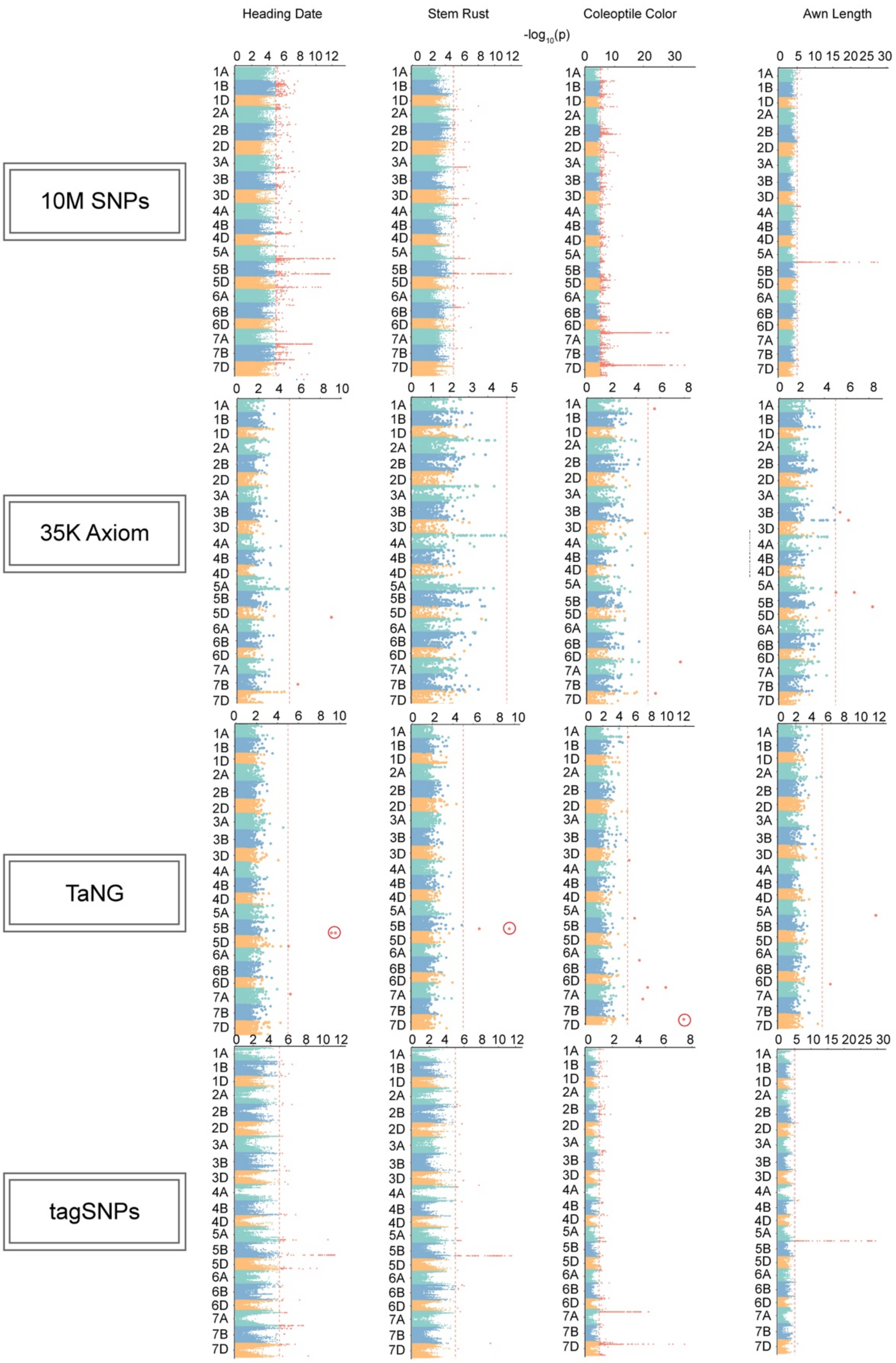

**Supplementary Fig. 30 | Comparison of the power and resolution for GWAS using different SNP datasets.** High-quality core 10 Mb SNPs, the 35K Axiom array, the newly design *Ta*NG array, and the haplotype-based diagnostic SNPs were used to test for heading date, stem rust, coleoptile color, and awn length. Red circles indicate MTA identified using the *Ta*NG array, but not the 35k Axiom array.

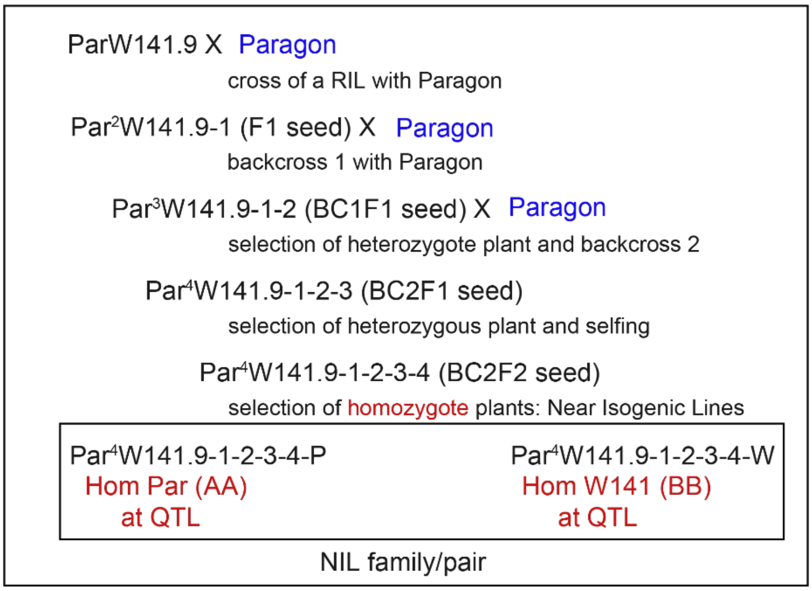

**Supplementary Fig. 31 | Near Isogenic Line development: The** crossing scheme with marker assisted selection is shown.
